# Constrained variation in the internal architecture of avian wing bones

**DOI:** 10.1101/2025.08.07.669070

**Authors:** F. Alfieri, O.E. Demuth, E.M. Steell, A.-C. Fabre, D.J. Field

**Affiliations:** Institute of Ecology and Evolution, Universität Bern, Baltzerstrasse 6, 3012 Bern, Switzerland; Department of Earth Sciences, University of Cambridge, Downing Street, Cambridge, CB2 3EQ, UK; Clare College, University of Cambridge, Trinity Lane, Cambridge, CB2 1TL, UK; Girton College, University of Cambridge, Cambridge, CB3 0JG, UK; Naturhistorisches Museum der Burgergemeinde, Bernastrasse 15, 3005 Bern, Switzerland; Department of Life Sciences, The Natural History Museum, Cromwell Rd, South Kensington, London SW7 5BD, UK; Museum of Zoology, University of Cambridge, Downing Street, Cambridge, CB2 3EJ, UK

**Keywords:** birds, ecomorphological convergence, flightless birds, forelimb, inner structure, marine birds

## Abstract

Extant birds exhibit remarkable ecological disparity accompanied by widespread skeletal convergence driven by functional adaptation. Investigations of morphofunctional associations with ecological factors have frequently focused on the external morphology of avian wing bones; however, the extent to which such associations also apply to the internal structure of the wing skeleton remains understudied. Here, we investigate disparity of the internal epiphyseal and diaphyseal structure of the avian humerus and ulna, and explore its correlates with ecology. Our dataset of 140 species spans extant bird diversity, and demonstrates that the internal structure of avian wing bones exhibits limited ecological signal beyond expected secondary trends related to flightlessness and marine habits. Our work instead shows that variation is primarily determined by body size, suggesting that functional constraints on internal wing bone structure imposed by flight are essentially universal across flying birds irrespective of most ecological habits and flight styles. Despite this broad lack of ecological signal, distinctive aspects of forelimb internal structure may facilitate the identification of flightless bird taxa in the fossil record.

## 1. Introduction

Identifying the functional drivers of phenotypic patterns requires disentangling extrinsic (e.g., ecological) and intrinsic (e.g., allometric, phylogenetic) factors (Jablonski 2022). The vertebrate skeleton exhibits traits of ecological significance (Currey 2002; Chirchir et al. 2017; Basu et al. 2019) that may be tightly linked to environmental interactions (Lieberman 1997; Ruff et al. 2006). For instance, internal bone structure (i.e. the distribution of bone tissue within skeletal elements) is associated with ecological habits in mammals, where it generally correlates with ecological parameters such as locomotor loadings, metabolism, and lifestyle (Kivell 2016; Alfieri et al. 2021, 2022; Amson and Bibi 2021). Bone structure has also been proposed to be more reflective of ecological habits than external bone morphology, both during an individual’s lifetime (Kivell 2016) and through generations (Alfieri et al. 2023). As such, bone structure may represent an informative source of data in investigations of fossil vertebrate ecology and functional morphology (Kitchell 1985; Chinsamy 2023). Nevertheless, it remains frequently overlooked in macroevolutionary studies of ecomorphology, possibly due to the time-intensive nature of data collection and analysis.

Crown birds (Neornithes) are among the most ecologically and taxonomically diverse clades of extant vertebrates. With >11,000 extant species and exceptional behavioral, morphological and size disparity (∼2 g in the smallest hummingbirds to over 120 kg in the Common Ostrich) (Wyles et al. 1983; Gill 2007; Barrowclough et al. 2016; Töpfer 2018), birds represent an ideal system in which to test relationships between ecology and bone structure, particularly as ecological and morphological convergence is abundant across avian phylogeny (e.g. Watanabe et al. 2020; Maderspacher 2022; Chen et al. 2025).

The morphology of the avian wing has been studied rigorously in the past, both in terms of external functional anatomy (e.g., size/shape, loading, aspect ratio, tip shape, e.g. Rayner 1988; Norberg 1990; Dial 2003; Wang and Clarke 2015; Baumgart et al. 2021; Beauchamp 2023) and internal anatomy (e.g., soft tissues and bones, e.g. De Margerie 2002; Nudds et al. 2007; Biewener 2011; Simons et al. 2011; Serrano et al. 2020). Investigations focusing on osteology have primarily focused on external bone morphology, and have revealed generally low degrees of disparity and overlapping morphotypes at macroevolutionary scales, that, nevertheless, also capture patterns driven by ecological factors (Nudds et al. 2007; Serrano et al. 2020).

Avian wing bones are hollow, lightweight, and characterized by a narrow and dense cortex, as well as bony struts/ridges and approximately circular cross-sections, with the bones of the stylopod and zeugopod functionally optimized to resist bending and torsion (de Margerie et al. 2005; Pennycuick 2008; Dumont 2010; Novitskaya et al. 2017; Sullivan et al. 2017). While this general configuration is expected to be functionally modulated by different ecological strategies, as suggested by some aspects of wing bone structural properties (de Margerie et al. 2005; Habib and Ruff 2008; Habib 2010; Simons et al. 2011), these patterns have only been explored in studies that are phylogenetically restricted (Simons et al. 2011; Frongia et al. 2018), relatively species-limited (de Margerie et al. 2005; Habib and Ruff 2008), or focused on specific ecological adaptations (Habib 2010). Moreover, to date these studies have predominantly focused only on the structure of the central shaft (‘diaphysis’) of wing long bones (de Margerie et al. 2005; Habib and Ruff 2008; Simons et al. 2011). The structure of the wing bone extremities (‘epiphyses’, i.e. ‘trabecular bone’), which in mammals are assumed to be more ecologically driven than diaphyseal structure (Kivell 2016), remains understudied in birds and has only been investigated on very restricted datasets (Kish 2011; Louis et al. 2022). Hence, a broad comparative investigation of links between avian ecology and wing bone diaphyseal and epiphyseal structure is needed. The absence of such work has, to date, precluded an understanding of macroevolutionary patterns of bird wing bone structure that may enable the identification of structural changes associated with ecological transitions through avian evolutionary history.

Here, we aim to determine how avian wing bone structure is associated with ecological diversity on a macroevolutionary scale. We studied the internal bone structure of the two primary wing-supporting elements, the humerus and ulna (de Margerie et al. 2005), across a broad phylogenetic sample (Fig. 1, Fig. 2). Flight style was expected to be the primary determinant of wing bone morphology because various flight behaviors are assumed to impose different constraints and/or loadings on wing bones, and, consequently their internal structure (Habib and Ruff 2008; Tobalske 2016). However, it has been hypothesized that wing anatomy may also reflect other ecological factors, such as migratory distance and associated mobility requirements, habitat and its intrinsic physical characteristics (e.g., air currents), or foraging strategy and related demands for agility (Baumgart et al. 2021; Beauchamp 2023). Thus, to broadly describe ecological diversity our dataset prioritized capturing variation in flight style, habitat, lifestyle, migration and feeding habits.

**Fig 1.**
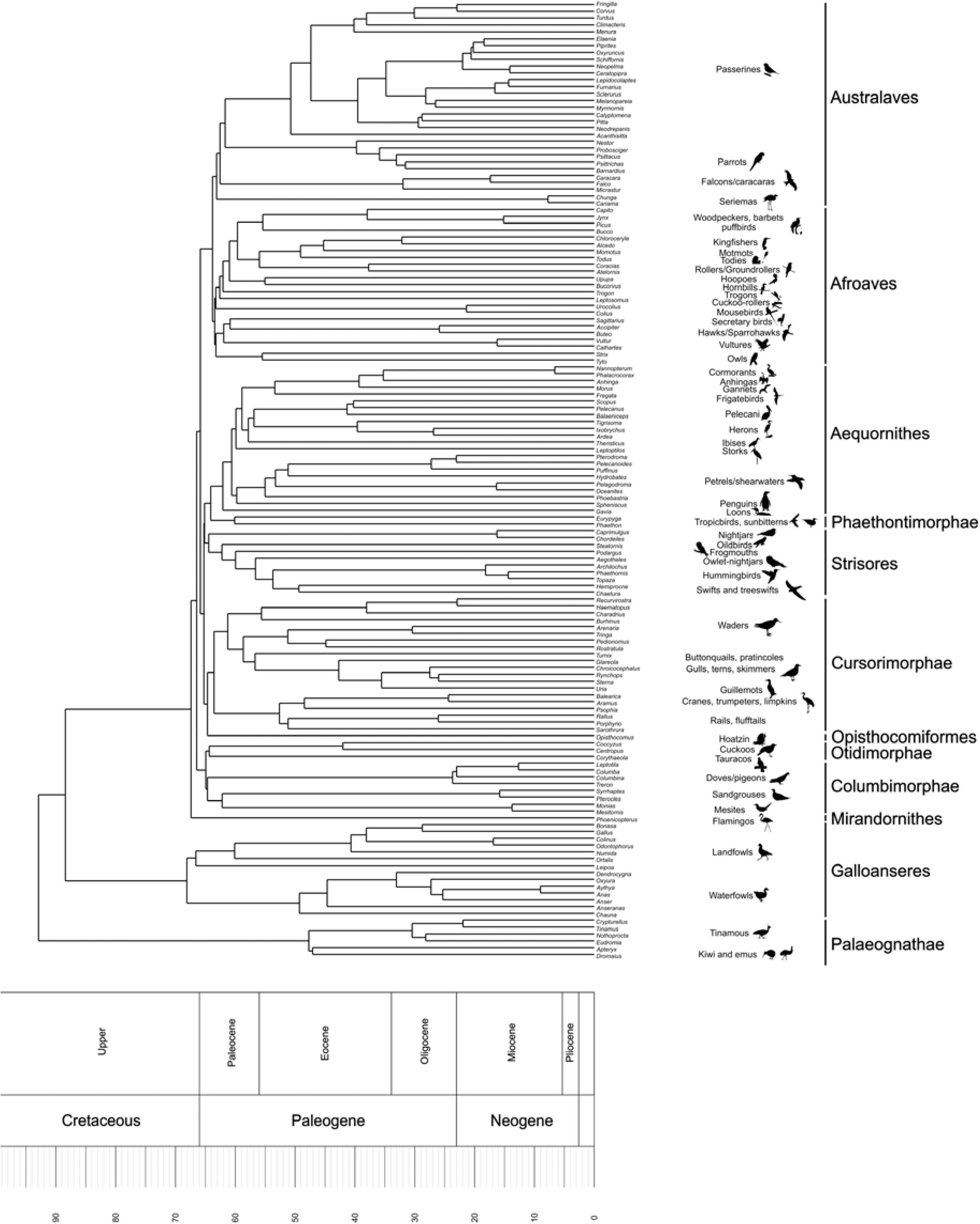
Taxa investigated in the present work (140 species), and their phylogenetic interrelationships (based on Stiller et al. 2024) (see Methods)

**Fig 2.**
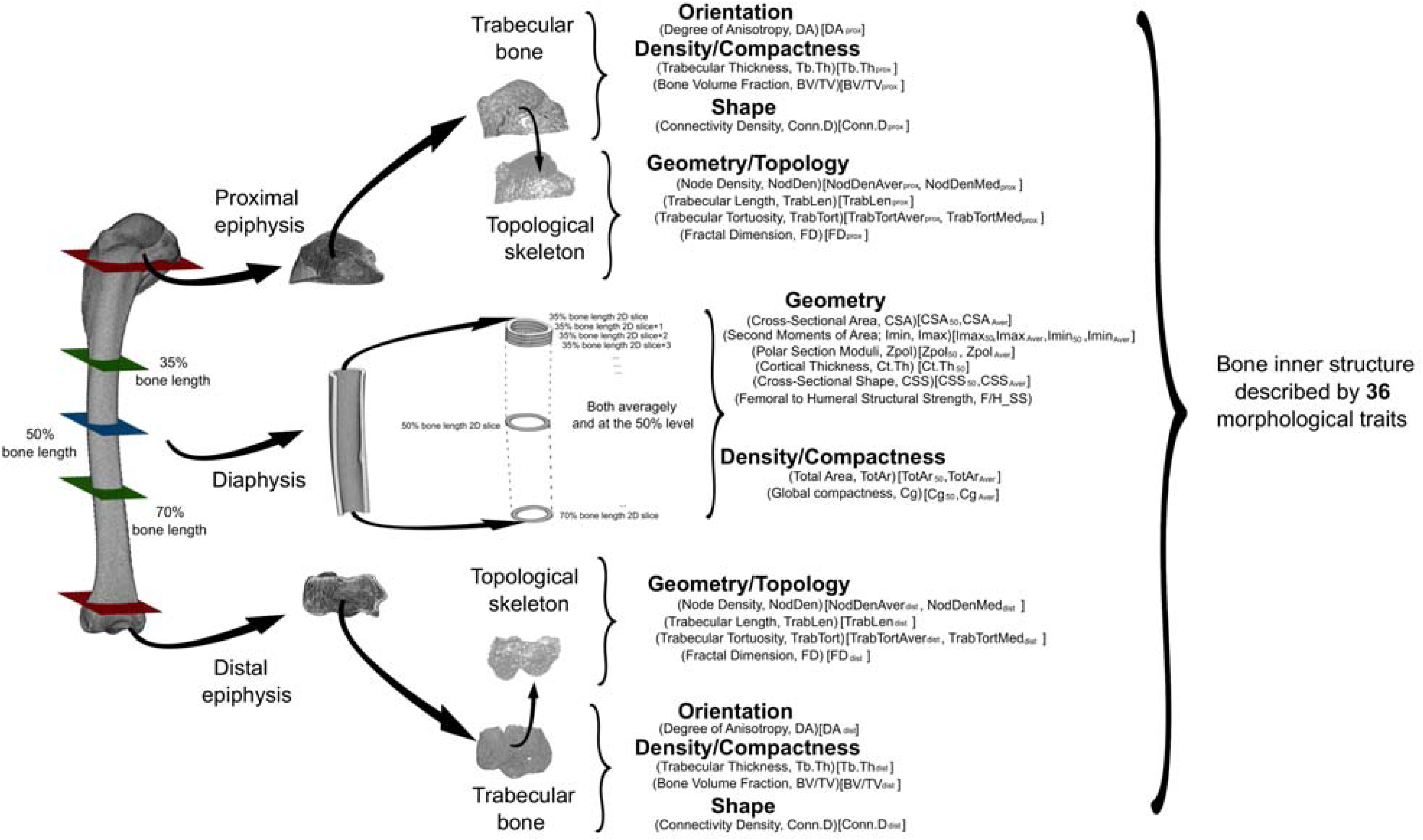
Summary of structural parameters extracted from bird wing bones in the present work, using the humerus of *Anser fabalis* NHMUK 1895-2-6-9. Physical properties (in bold) and associated structural parameters (in round and square brackets) are indicated for each anatomical region investigated. Further details provided in Methods, and Supplementary S2-S4.

## 2. Materials and Methods

### 2.1. Wing bone structure data collection

We collected micro-computed tomography (μCT) scans de novo by scanning museum specimens (Table S1, Section S1, Supplementary Material), and downloaded data from MorphoSource (https://www.morphosource.org; projects ID: 00000C420, Bjarnason and Benson 2021,; ID: 00000C580).

Our sample comprises adult skeletons of 140 extant species (and genera) (Table 1) spanning the phylogenetic diversity of crown birds, encompassing 98 family-level clades and 37 order-level clades (Fig. 1). μCT voxel sizes ranged from 0.0146 to 0.127 mm (average 0.059 mm). For each taxon, we studied one humerus, one ulna, and one femur (to assess another potentially informative trait from the diaphysis; see below).

**Table 1.** Replication statement.

| Scale of inference | Scale at which the factor of interest is applied | Number of replicates at the appropriate scale |
| --- | --- | --- |
| Species (but representative of macroevolutionary patterns) | Species (n=140, representative of 140 bird genera, spanning extant bird diversity) | NA (the work being framed in a macroevolutionary context, it was not possible to have replicates) |

Using VGSTUDIO MAX 3.5.4 (Volume Graphics), we aligned the bones in a standardized orientation to study anatomically homologous traits across our taxon sample (Section S2). The oriented bones were imported in FIJI (Schindelin et al. 2012), where they were partitioned into their proximal/distal epiphyses and diaphysis. These anatomical regions were defined in standard ways: the diaphysis was defined by isolating the central 35% of the entire bone length, and the epiphyses by identifying specific anatomical markers (Section S3) (Fig. 2). Epiphyses were pre-processed and binarised into ‘bone’ and ‘non-bone’ fractions in FIJI (Section S4), and imported in R (R 4.4.1; R Core Team 2024), where a Region of Interest (ROI) including only trabecular bone was isolated. We quantified the following initial suite of trabecular parameters (TP) from the ROIs using the FIJI plugin *BoneJ2* (Domander et al. 2021): ‘degree of anisotropy’ (DA, unitless), ‘trabecular thickness’ (Tb.Th, mm), ‘bone volume fraction’ (BV/TV, unitless) and ‘connectivity density’ (Conn.D, mm^-3^). Additional trabecular variables were computed in R from topological skeletons (in turn extracted from the ROIs, in Avizo 2020.2), in which nodes and branches represent trabecular connections and trabeculae, respectively, in order to summarise the topological properties of trabecular networks (Veneziano et al. 2021; Alfieri et al. 2025). These variables were ‘node density’ (NodDen, represented by an arithmetic mean, NodDenAver, and a median value, NodDenMed), ‘trabecular tortuosity’ (TrabTort, represented by an arithmetic mean, TrabTortAver, and a median value, TrabTortMed), ‘average trabecular length’ (TrabLen), and ‘fractal dimension’ (FD). For both the humerus and the ulna, we use the subscripts TP_prox_ and TP_dist_ to denote proximal and distal epiphyseal TPs, respectively.

In FIJI, we extracted the following diaphyseal cross-sectional properties (CSP) (Fig. 2): ‘global compactness’ (Cg, %), ‘total cross-sectional area’ (TotAr, mm^2^), ‘cross-sectional area’ (CSA, mm^2^), ‘second moments of area’ (around the minor, Imax, and major axes, Imin; mm^4^), ‘polar section modulus’ (Zpol, mm^3^), ‘cross-sectional shape’ (Imax/Imin, unitless) and ‘perimeter’ (mm). The perimeter was only used to estimate a body mass proxy (see below). These CSPs were computed as both mean (i.e. averaged across the entire diaphysis) and mid-diaphyseal (i.e. from the 50% level of the bone length) values, referred to as CSP_Aver_ and CSP_50_, respectively. At the mid-diaphysis, we additionally measured the humeral and ulnar ‘cortical thickness’ (Ct.Th_50_, mm) and the ‘relative femoral to humeral structural 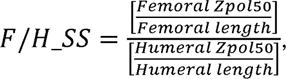, computing the latter as previously proposed (Habib and Ruff 2008). For further details on trabecular bone isolation, TPs and CSPs quantification, see Sections S5-S6, and further references (Veneziano et al. 2021; Alfieri et al. 2025).

Overall, we collected 36 and 35 structural traits to represent humeri and ulnae, respectively (ulnar traits do not include F/H_SS) (Table S1). As summarized in Fig. 2, these traits inform on several physical properties of bone structure and were previously proposed as reflecting diverse ecological aspects in tetrapods, as locomotor behaviour (through differential bone response to biomechanical loadings), lifestyle (e.g., terrestrial *vs.* aquatic) or other related aspects (e.g., metabolic rates, energy saving strategy) (e.g. de Margerie et al. 2005; Habib and Ruff 2008; Padian 2011; Mitchell 2016; Chinsamy et al. 2020; Alfieri et al. 2021, 2022, 2023, 2025; Veneziano et al. 2021 and references therein). We split the traits according to the anatomical regions that they represent, obtaining six multivariate datasets (three per bone): humeral TP_prox_, humeral CSP, humeral TP_dist_, ulnar TP_prox_, ulnar CSP and ulnar TP_dist_. From humeral/ulnar TP_prox_ and TP_dist_ we discarded the specimens with a too low number of trabeculae (see Section S5).

### 2.2. Ecological data collection

To capture broad aspects of locomotor ecology across our taxon sample, we collected data on flight style, habitat, primary lifestyle, migration, trophic level and trophic niche. Individual bird species can exhibit multiple flight behaviours; therefore, flight style was characterized into 10 discrete, non-mutually-exclusive variables (Table 2). With the only exception of the Infrequent Flight category, which has three levels, for the other flight behaviours taxa were scored with binary variables based on whether they have been observed/reported to exhibit the behaviour (score: 1; i.e., presence) or not (score: 0; i.e., absence). The dataset was compiled using data from the Birds of the World database (Billerman et al., 2025) and primary literature (see Section S7).

**Table 2.** Flight categories applied in the present work. Description of locomotory categories adapted from Baliga et al. (2019) and Lowi-Merri et al. 2021)

| 860 | Flight category | Description |
| --- | --- | --- |
| 861 | **Hovering** | Ability to hover without needing to rely on wind conditions, i.e., active, unassisted and muscle driven |
| 862 | **Intermittent flight** | Includes both bounding - i.e. short bursts of flapping interspersed with short intervals where the wings are folded against the body - and flap gliding - short bursts of flapping interspersed with short intervals of gliding where the wings are held in extended position |
| 863 |  |  |
| 864 | **Gliding** | Wings are held in extended position for considerable amount of time without flapping, often resulting in decreasing altitude |
| 865 | **Continuous flapping** | Flight that primarily involves flapping continuously without interspersed episodes of gliding/soaring or bounding behaviors |
| 866 | **Soaring** | Includes thermal soaring - i.e. using rising thermals to increase altitude -, slope soaring - i.e. increasing altitude due to harnessing rising air along a slope - and dynamic soaring- i.e. maintaining or increasing altitude through the use of wind speed gradients, e.g., over a large body of water, and own momentum |
| 867 | **Stooping** | Rapid, controlled, head-first dives, often with tucked-in wings |
| 868 | **Foot-propelled swimming** | Feet but not wings are used to propel and steer underwater |
| 869 | **Wing-propelled swimming** | Wings but not feet are used to propel and steer underwater |
| 870 | **Burst flight** | Explosive burst of flapping in takeoff (often to escape danger) usually followed by short gliding. Not to be confused with escape flights of generally poor fliers |
| 871 | **Infrequent flight** | Level 0 = flying; level 1 = any behaviour that includes mostly ground-dwelling, seasonable incapability of flight due to molting or other factors, aversion to flight, and/or poor flyer; level 2 = permanent flightlessness |

The other ecological variables were included as categorical variables from the AVONET2_eBird dataset (Tobias et al. 2022), which was preferred over AVONET1 and AVONET3 as it is the only one without missing values for the taxa in our dataset. Overall, our dataset (Table S2) comprises 15 binary/categorical/ordinal variables that collectively provide an extensive description of avian ecology. All analyses were performed in the R statistical environment (R Core Team 2024).

### 2.3. Body mass proxy computation

Structural variables are known to relate to body mass (e.g. Doube et al. 2011), therefore we extracted a body mass proxy (BMp) for each sampled species to account for BMp-related patterns. We selected skeletal correlates over average species mass, as the latter may not fully account for intraspecific variability. Specifically, we used a proxy directly informing on the specimens we examined to extract TPs and CSPs (although mass values and skeletal correlates often strongly co-vary, see below). Hence, we took the humeral perimeter (unit: mm, exploiting Perimeter_50_ from CSP computation, see above), which is a strong skeletal correlate for body mass in birds (Field et al. 2013), and log10 transformed it, to account for outliers. Since the predictive power of humeral perimeter is limited to flying birds (Field et al. 2013), humeral log10-Perimeter_50_ for flying birds (i.e., IF=0, as defined in Table 2, ca. 76% of the studied taxa, Tables S1-S2) was regressed against the log10-transformed respective species mass (unit=g, averaged from AVONET1_BirdLife, AVONET2_eBird and AVONET3_BirdTree, Tobias et al. 2022). Humeral log10-Perimeter_50_ and log10-mass strongly co-vary (R^2^= 0.90, *p*<0.001) through the equation ‘log10- Perimeter_50_= 0.29 + (0.35930 * log10-mass)’ (both *p<*0.001). The linear model built for flying birds was then used to predict (‘predict.lm’ function) log10-Perimeter_50_ for the other birds, yielding BMp for the full sample.

### 2.4. Statistical analysis

#### 2.4.1. Ecological clustering

On a dissimilarity matrix (‘daisy’ function, metric=‘gower’, ‘cluster’ package, Maechler et al. 2022)), in turn derived from the ecological dataset, we ran hierarchical clustering (‘hclust’ function, method=’ward.D2’). We quantified the optimal number of ecological clusters (‘eco-clusters’, hereafter) through the gap statistic (‘clusGap’ function, ‘cluster’) (Fig. S1) using 500 bootstrap replicates whereby variation in dispersion within clusters was compared to a null distribution (Tibshirani et al. 2001; Baliga et al. 2019). The optimal eco-cluster number was used to cut the hierarchical cluster tree (‘cutree’ function) and to identify which taxa were assigned to each eco-cluster by the hierarchical clustering. This was visualized through a Principal Coordinate Analysis (PCoA, ‘cmdscale’ function) PCo1-PCo2 plot (Fig. S2). Then, we computed correlations between PCos and ecological variable levels to visualize how ecological variables contribute to discriminating eco-clusters (Fig. S3). This allowed us to assess which ecological features primarily define each eco-cluster and, when needed, to re-assign taxa to alternative eco-clusters (as detailed and justified in Section S8) (Fig. 3, Fig. S4). Our data represents 140 different bird genera. We considered interspecific differences within the same genus as negligible across the phylogenetic scale under investigation.

**Fig 3.**
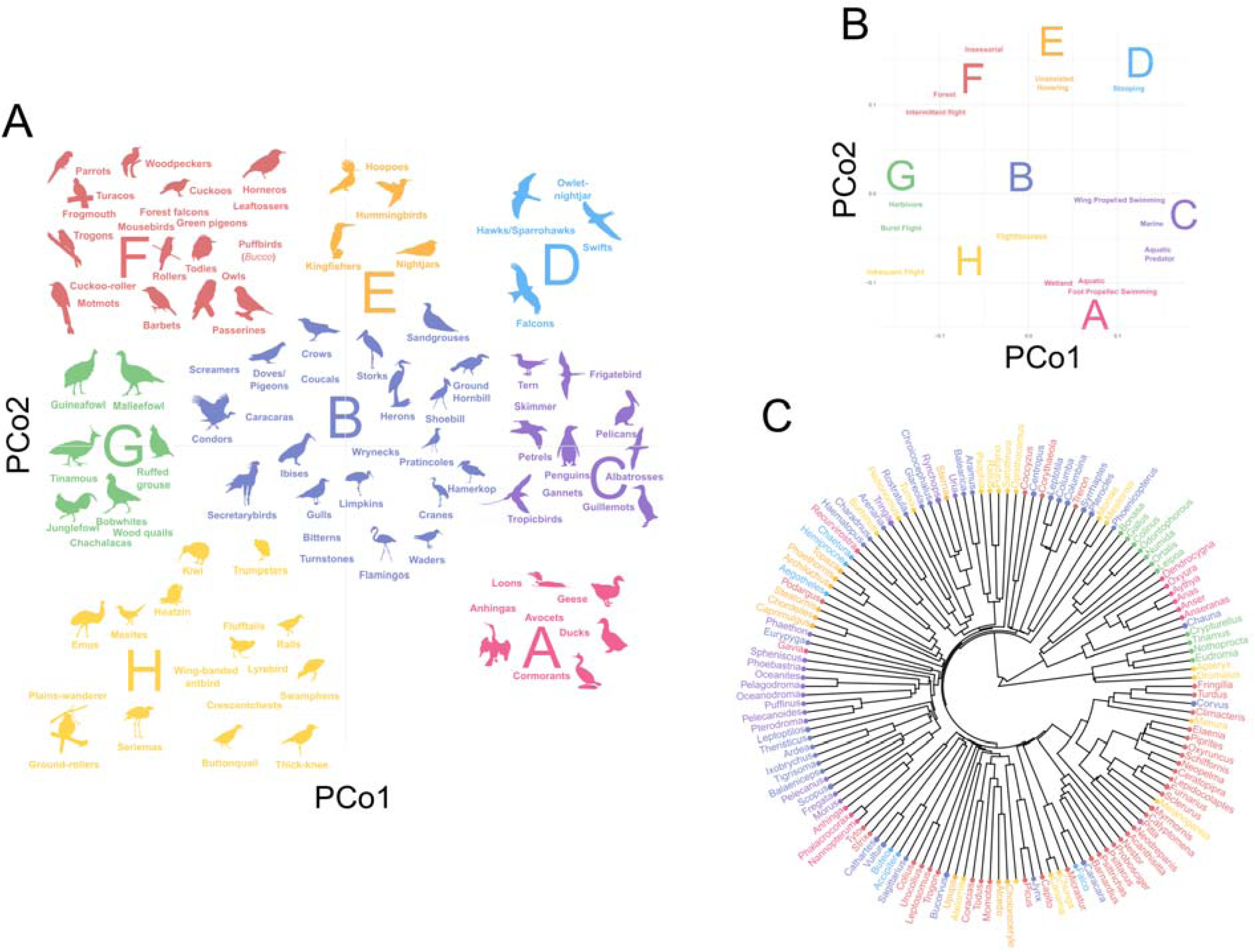
A. Simplified version of the PCo1-PCo2 scatterplot, showing the bird groups represented within each eco-cluster (Exact position of silhouettes and names do not represent precise PCo scores, which are shown in Fig. S4). **B.** Plot indicating the influence of particular ecological variables on separation of eco- clusters (deriving from Fig. S3). Only the ecological variables defining each eco-cluster are shown (as detailed in Section S8). Variable text positions indicate how strongly a specific variable contributes to eco-cluster separation (i.e. for peripheral clusters, more peripheral variable levels contribute more strongly to cluster differentiation). **C.** Illustration of the phylogenetic distribution of the eight eco-clusters, following the topology of Stiller et al. 2024 (subsequently adapted through inference from other works, as detailed in Section S9)

#### 2.4.2. Ecological influences on wing bone structure

We analysed the six structural datasets separately to identify differential ecological signal in epiphyses *vs.* diaphysis in humeral/ulnar structural variability. All the inferential tests were phylogenetically informed. Phylogenetic relationships and divergence times for the species in our dataset were primarily based on Stiller et al. (2024). Species present in Stiller et al. but not in our sample were removed and replaced with those represented in our sample when the same genera were represented (details in Section S9). Multivariate analyses were undertaken to allow us to identify strong interrelated variables (a frequent condition for structural traits, Cotter et al. 2009) and to detect patterns potentially obscured by univariate analyses (Ryan and Shaw 2012; Alfieri et al. 2025). Before running multivariate analyses, we visualized the distribution of residuals of a series of phylogenetic Generalized Least Square regressions (PGLS) (‘gls’ function, ‘nlme’ package (Pinheiro et al. 2020) of each single trait against eco- clusters. If the distribution of residuals in the respective PGLS strongly deviated from normality, the trait was log10-transformed. After scaling and centering each multivariate structural dataset, we tested for ecological effects while accounting for allometric and phylogenetic influences. To do so, we regressed the datasets against eco-clusters, with BMp as a co-variate, through multivariate PGLSs (mvPGLS) (‘mvgls’ function, ‘mvMORPH’ package, Clavel et al. 2015). BMp was not significantly related to eco-clusters (*p*=0.1; pANOVA) and, accordingly, ecological and allometric effects are separated in each mvPGLS. Thus, no interaction effects needed to be tested. For each structural dataset, we preliminarily fitted four mvPGLSs, each assuming one of the following evolutionary models: Brownian Motion (BM), Ornstein-Uhlenbeck (OU), Early Burst (EB) and Pagel’s lambda. The best fitting model for each anatomical region was identified through Extended Information Criterion (EIC, ‘mvMORPH’) (Table S12) and then used to fit the mvPGLS employed in the following steps. Ecological and allometric effects on humeral/ulnar structural datasets were tested through phylogenetic MANCOVAs (pMANCOVA) (test = ‘Pillai’, type = ‘II’) and quantified through measures of effect size (both through the ‘mvMORPH’ package). In the case of a significant relationship between structure and eco-clusters, we performed pairwise comparisons to identify which eco-clusters were significantly different (‘mvMORPH’, test = ‘Pillai’, adjust="Bonferroni", nperm=1000) (Table 3, Tables S13-S17).

**Table 3.**
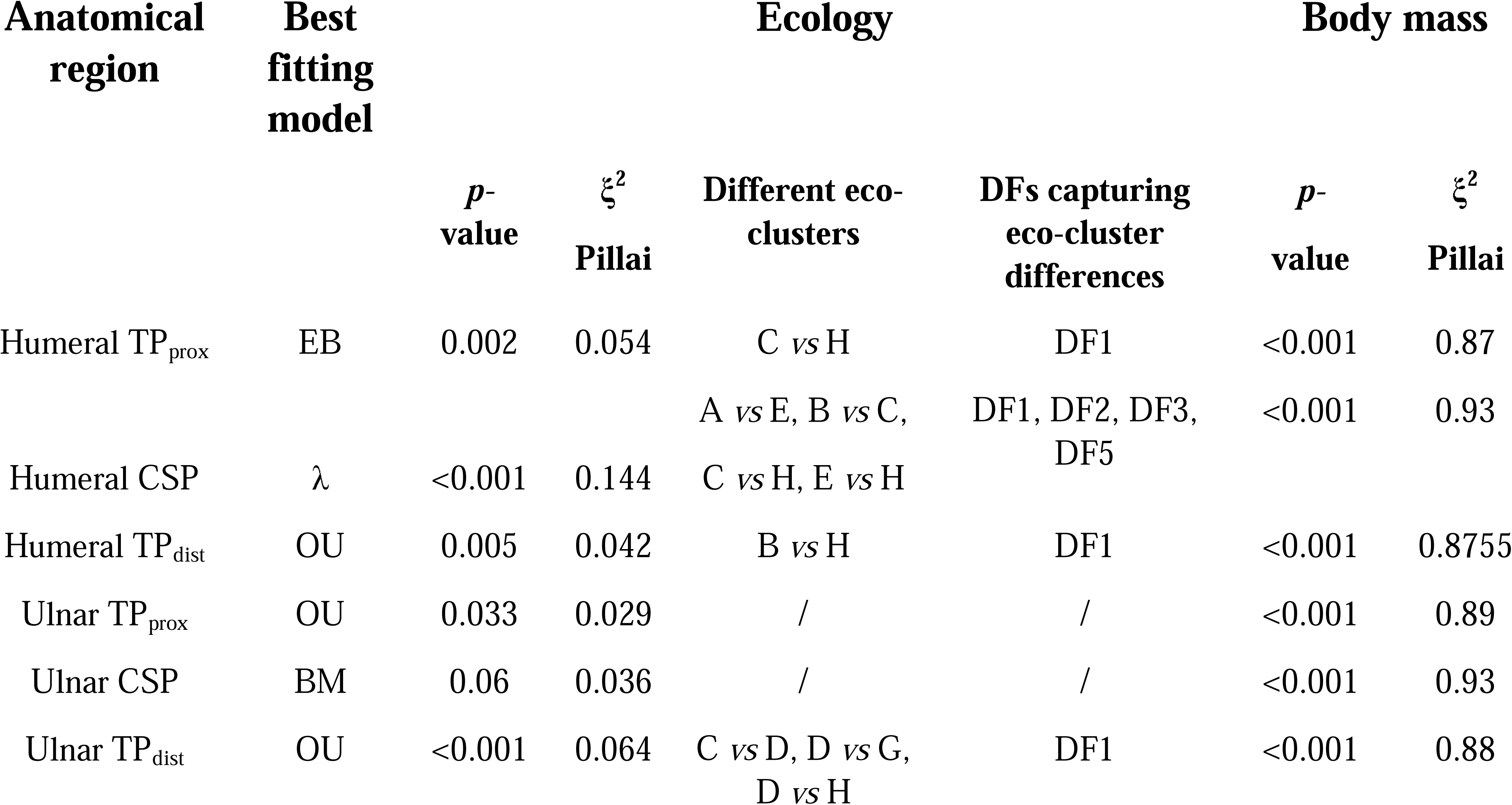
Analytical Results. For each of the six anatomical regions investigated, the following parameters are shown: the best fitting evolutionar model, results for multivariate PGLS regressions against eco-clusters and body mass proxy (‘Ecology’ *p*-value and ‘Body mass’ *p*-value, significant if *p*-value <0.05), effect size measures (ξ914 2 Pillai, informing on how strongly ecology and body mass contribute to wing structural patterns), eco-clusters yielding significance after pMANCOVA pairwise comparisons, and which DFs (deriving from DFA) discriminate among those eco-clusters (determined through DFs *vs.* eco-clusters pANOVAs).

To assess structural variability across eco-clusters, we used the respective mvPGLS for each anatomical region to run a phylogenetic Discriminant Function Analysis (pDFA) (‘mvMORPH’). For each of the six pDFAs, we generated biplots with the first three Discriminant Functions (DF), i.e. DF1, DF2 and DF3, mapping eco-clusters onto them (Figs. S5-S13). We tested whether and how the first three DFs are related to BMp (Table S18) to assess how much allometric variation they include. We focused on significantly different eco-clusters and identified the discriminating DFs (Table S19), then visualized patterns through DF *vs.* eco-clusters boxplots (Figs. S5-S13). This also enabled extraction of structural traits that substantially contribute to these DFs (as suggested by pDFAs standardized coefficients, Tables S20-S21). As with the DFs to which they strongly contribute, we also generated boxplots for these structural traits to visualize how they discriminate among the significantly different eco-clusters. If a structural dataset yielded no significant ecological effect or if pairwise comparisons yielded no significant differences among eco-clusters, we generated boxplots for the traits contributing most to DF1-DF3 across all eco-clusters. In addition to boxplots, we generated traitgrams, i.e. mapping the trait values on the phylogeny (‘phytools’, Revell 2012), for traits substantially contributing to DFs of interest, see above) (Figs. S5-S13). Allometric effects were separated from those of eco-clusters in all multivariate inferential analyses (see above), but this did not occur in datasets from which we extracted variables to generate univariate plots. Hence, to generate boxplots and traitgrams we used size-corrected values if the trait was significantly related to BMp (tested through PGLS). Residuals of the PGLS were used for size correction and size-corrected traits are hereafter indicated with the prefix ‘SC-’.

## 3. Results

Gap statistics yielded nine as the optimal number of eco-clusters to describe variation in our dataset (Fig. S1). We then detected which taxa belong to which eco-clusters and identified their corresponding ecological features (see PCo1-PCo2 biplot, Fig. S2, and ecological variables contribution, Fig. S3). This preliminary scheme of nine eco-clusters was edited by reassigning 19 taxa, i.e. 13.5% of the sample, to other eco-clusters, as justified in Section S8. These adjustments involved eliminating one of the nine eco- clusters, since it overlapped with other eco-clusters and was represented by only six taxa, which were reassigned to other groups. Hence, we obtained a final scheme of eight groups, i.e. eco-clusters A-H (Fig. S3), characterized by distinctive ecological features (summarized in Fig. 3B). The latter were detected for all eco-clusters except eco-cluster B (including e.g., pigeons and storks) which did not yield any unambiguously diagnostic ecological attributes (Fig. 3B). Hence, birds belonging to eco-cluster B were considered ecologically generalized, consistent with their central position in the PCo1-PCo2 biplot (Fig. S3, Fig. 3). Eco-clusters are not significantly related to BMp (see above), and their distribution on the time-tree suggests that they are also largely independent of phylogeny (e.g., eco-cluster H includes representatives of Australaves, Afroaves and Cursorimorphae) (Fig. 3C). This potentially allows us to identify morphological convergences associated with similar ecological features (Fig. 3C).

Depending on the wing element and anatomical region, different evolutionary models differentially explain structural trait diversification in birds (Table S11). All structural datasets are significantly related to BMp, and five out of six datasets (i.e. all except ulnar CSP) relate to ecology (Table 3). Allometric effects are largely dominant (BMp ξ^2^= 0.87-0.93), while eco-clusters have weak effects (ξ^2^=0.029-0.064 in five out of six regions; ξ^2^=0.144 for humeral CSP). Within the datasets significantly related to ecology, ulnar TP_prox_ does not yield any significant difference between eco-clusters after Bonferroni correction. The few comparisons yielding significance across the other four regions (ca. 6% of the total comparisons) are primarily driven by two eco-clusters: eco-cluster C (marine aquatic predators performing wing-propelled swimming, e.g., penguins and auks) and eco-cluster H (flightless/poorly flying birds, e.g., emu, kiwi; not including burst flying taxa, e.g., tinamous, that comprise a separate eco-cluster). Indeed, eco-clusters C and H are the most commonly contributing to significant differences, being involved in 4/9 and 5/9, respectively, of the comparisons yielding significance across several anatomical levels. Minor influences are probably due to other eco-clusters involved in the comparisons yielding significant differences, i.e. eco-cluster D involved in 3/9 comparisons but at one single region (ulna TP_dist_), eco-cluster B and E involved in 2/9 comparisons, eco-clusters A and G involved in 1/9 comparison (Table 3, Tables S13-S17).

In pDFA biplots for all the structural regions, most of the taxa occupy a restricted region, i.e. wing bone structure is largely homogeneous in birds, and eco-clusters overlap substantially (Figs. S5-S13). Allometric effects are substantially involved in Discriminant Functions (DF), as suggested by DFs-BMp correlations being significant for 83% of the DFs, and by BMp explaining up to the 91% of DFs variance (e.g., humeral CSP DF3) (Table S18).

In the epiphyseal regions, significant differences were captured by some DFs (Table 3, Table S19) strongly driven by NodDen. The only epiphyseal region that did not yield significant pairwise comparisons was ulnar TP_prox_, for which the first three DFs were largely driven by NodDen too (Tables S20). Significant differences for the humeral TPs are likely due to high values shown by the flightless emu (*Dromaius*) and kiwi (*Apteryx*) (Fig. S5F-G, Fig. S10D-E), both within eco-cluster H (Fig. 3). Traitgrams for humeral SC-NodDen show that high values for palaeognaths are extremely unusual across avian phylogeny, with the only possible exception being the swift *Chaetura* (Apodiformes: Apodidae). For humeral TP_dist_, a relatively high NodDen is also shown by the cormorant (*Nannopterum*) and the penguin (*Spheniscus*). The humeral TP_dist_ traitgram also enabled identification of extremely low SC-NodDen values for a hummingbird (*Topaza*) and a passerine (*Corvus*) (Fig. S6C-D; Fig. S10F-G). Only in TP_prox_, the DF responsible for significant differences is additionally driven by SC-BV/TV, SC-TbTh and SC-FD. Boxplot for SC-BV/TV does not reveal any pattern of interest, i.e. eco-cluster C’s distribution lies entirely within that of eco-cluster H (Fig. S5D). Boxplots for SC-TbTh and SC-FD show extremely low values for the emu (Fig. 5E-H).

As for the ulnar TPs, SC-NodDen boxplots evidence higher values for *Dromaius*. Since *Apteryx*’s ulnar epiphyseal data were discarded (due to an extremely low number of trabeculae, as detailed in Methods), it is not possible to know if it resembles *Dromaius* in ulnar node density, as it does for the humerus (see above). *Chaetura*, *Spheniscus* and *Nannopterum* yielded relatively higher ulnar SC-NodDen (mirroring humeral patterns, see above) (Fig. 4A, Fig. S11C-D, Fig. S13F-I).

**Fig 4.**
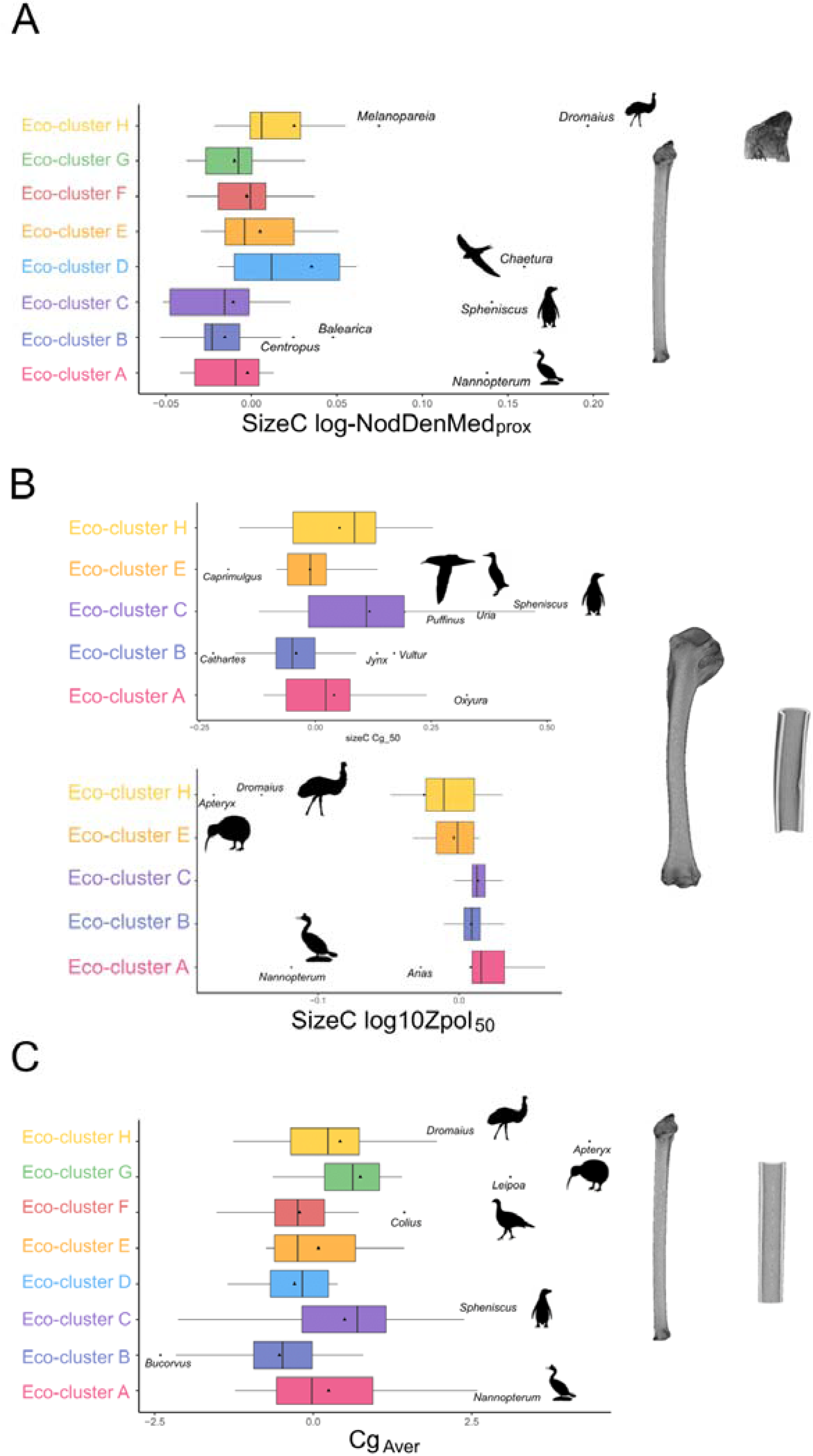
A: Boxplot showing proximal ulna size-corrected log10-NodDenMed across the eight eco-clusters. **E:** Boxplots showing humeral size-corrected Cg_50_ (above) and log10-Zpol_50_ (below) across the eco- clusters A, B, C and E; **C:** Boxplot showing ulna Cg_Aver_ across the eight eco-clusters.

In the humeral diaphyseal region, significant differences were captured by some DFs (Table 3, Table S19) which were mainly driven by six properties (TotAr, Cg, CSA, Imin, Imax and Zpol). They strongly contribute through both the mid-diaphyseal (CSP_50_) and average diaphyseal (CSP_Aver_) measures (Table S20). Through extremely low values, five variables - i.e. SC-CSA, SC-Imax, SC-Zpol, SC-TotAr and SC- Imin - primarily discriminate *Dromaius* and *Apteryx*, especially, together with *Nannopterum*, from all other taxa in our dataset. Although less distinctive, a tendency to show low values for some of these variables is also yielded by a dabbling duck (*Anas,* for SC-TotAr, SC-Imax, SC-Zpol, SC-Imin) and *Spheniscus*, for SC-TotAr and SC-Imin) (Fig. 4B, Fig. S8). This minor pattern shown by *Spheniscus* is likely not sufficient to explain the frequent significant differences between eco-clusters involving eco- cluster C (to *Spheniscus* belongs). The trait likely driving these discriminations is therefore possibly represented by humeral SC-Cg, for which eco-cluster 2 consistently shows higher values compared to the others (Fig. 4B, Fig. 8M-O). The higher values for eco-cluster 2 are caused by extremely high SC-Cg yielded by *Spheniscus*, the auk (*Uria*) and the shearwater (*Puffinus*) (see traitgram, Fig. S9B, S9D. Humeral CSS (examined due to the relatively strong contribution of this property to DF5, Table S20) reveals that *Spheniscus*, hummingbirds, *Uria* and, to a lesser extent, *Puffinus* are distinct for higher values, compared to all the other birds (Fig. S8M-N, S9L-M).

Although ulnar CSP did not yield any relationship with ecology, examining its DF1-DF3 and the traits strongly contributing to them allowed us to identify the extremely low SC-TotAr, SC-CSA, SC- Imin, SC-Imax and SC-Zpol yielded by *Apteryx*, belonging to eco-cluster H (Fig. S12). *Apteryx*’s extremely low values are consistently (i.e. for all traits) approached by *Dromaius*, *Chaetura* and *Nannopterum*, i.e. they tend to have lower values compared to other taxa, but not to the same extent as *Apteryx* (Fig. S12). The latter is also distinctive for a higher ulnar Cg, with this trend partially mirrored by *Dromaius*, *Nannopterum*, *Spheniscus* and the malleefowl *Leipoa* (Fig. 4C). A relatively high ulnar Cg is also apparent for *Columba*, but only for Cg_50,_ and it neatly contrasts with the Cg_Aver_ pattern (Fig. S12 E-F).

## 4. Discussion

### 4.1. Ecological factors do not drive the evolution of avian wing bone internal structure among crown birds

Our results show that internal bone structure in bird wings is not primarily influenced by ecological variation. Indeed, although five out of six studied anatomical regions yielded a significant relationship with eco-clusters, effect size measures indicate weak ecological effects on wing structure overall compared to allometric effects (Table 3). The relatively weak ecological effects on humeral/ulnar structure are also supported by the limited number of pairwise comparisons yielding significance (ca. 6% of the total comparisons, Table 3, Tables S13-S17) and the extensive eco-cluster overlap on our phylogenetic Discriminant Functional Analysis (pDFA) plots. Biplots built with Discriminant Functions (DFs) also illustrate the relatively restricted regions occupied by most studied taxa. This pattern points to a relatively homogeneous and conserved internal wing bone structure that is widespread across bird phylogeny irrespective of ecological habits, suggesting that morphological constraints imposed by flight, regardless of specific flight style, generally dictate wing bone structure. Previous studies on wing bone structure mainly focused on the diaphysis, and investigated a relatively narrow range of ecological categories (Habib and Ruff 2008) or taxonomic groups (e.g. Simons et al. 2011), or focused on individual ecological habits (e.g., wing-propelled diving, 38). These studies do not provide wide-scale comparative frameworks, yet our results align with the general macroevolutionary patterns in avian humeral external morphology highlighted by Serrano et al. (2020). Indeed, both wing bone internal structure (as shown by this work) and humeral external geometry (as shown by Serrano et al.) appear minimally disparate and relatively conserved, with a few distinct trends that may represent adaptive optima departing from a generally conserved configuration likely corresponding to secondary specializations towards highly distinctive ecologies (Serrano et al. 2020) (see below). Therefore, we can hypothesise that a conserved structural design is widespread across bird phylogeny and that this represents a functional optimum that limits further diversification of internal bone morphology. Several previous investigations on bird wing bones described them as hollow and lightweight on average, with a thin and dense layer of cortical bone, showing internal osseous struts/ridges, rounded transverse sections and structural specialisations to resist bending and torsion loadings. This general configuration has been derived from studying wing bone structure in few bird taxa (e.g. de Margerie et al. 2005; Pennycuick 2008; Dumont 2010; Novitskaya et al. 2017; Sullivan et al. 2017). Since our work is the first macroevolutionary analysis of bird wing bone structural traits over a diverse and large sample of species, we can assume that the limited disparity and overlapping structure of most of the studied taxa reflects constraints imposed by these general structural demands. Only highly distinctive functional regimes, such as loss of flight and/or adaptations to a marine habitat, appear to drive consistent, discernible deviations from these broadly conserved structural characteristics of the avian wing.

Wing bone structural variation is instead primarily influenced by allometry. Indeed, all structural levels yielded significant relationships with body mass proxy (BMp) and a strong BMp effect size (Table 3). To our knowledge, no allometric studies of bird wing bone trabecular parameters have previously been undertaken across a taxonomically broad sample, with only one study previously investigating humeral diaphyseal structural properties (Cubo and Casinos 1998). Determining specific allometric trends (e.g., positive/negative allometry) is beyond the scope of this work, but our dataset could be utilized and expanded in future investigations aiming to test whether allometric patterns previously proposed for birds (for diaphyseal, Cubo and Casinos 1998, and epiphyseal traits, the latter focusing on the femur, Doube et al. 2011), hold for the wing bone structural traits quantified here. Trabecular parameters from non-avian reptile humeral metaphyses suggest that general patterns may hold across amniotes (Plasse et al. 2019), but the specific flight-related adaptations of avian dinosaurs (aside from known allometric scaling variability among clades or skeletal elements, Kivell 2016) could be explored further in focused investigations. Our results align with those from a recent macroevolutionary analysis of amniote femoral head trabecular parameters, which highlighted strong effects of body mass (Gônet et al. 2023). It is becoming increasingly clear from macroevolutionary studies of bone structure (e.g. Amson and Bibi 2021; Alfieri et al. 2025) at wide taxonomic scales that the predominance of allometric effects should be expected. From this strong allometry-driven variation, secondary, yet interesting, ecological patterns can be identified.

While generally low disparity is shown by both external humeral anatomy (Serrano et al. 2020) and internal wing bone structure, the external humeral morphology shows distinctive trends related to aerial hovering and aquatic diving (Serrano et al. 2020), whereas we found distinct structural patterns potentially related to flightlessness and/or a marine lifestyle. Moreover, our results imply that humeral and ulnar proximal and distal epiphyses evolved following different evolutionary models relative to the diaphysis (i.e. Ornstein–Uhlenbeck, OU, for all epiphyses except that of the proximal humerus, which instead followed Early Burst, EB; contrasting with humeral and ulnar diaphyses, which are explained by λ and Brownian Motion, BM, respectively; Table 3). In addition, the humeral diaphysis was the level most strongly influenced by ecological effects as it was the only region with an eco-cluster effect size larger than 0.1, and it yielded a substantial number of significant pairwise comparisons (Table 3). Thus, morphological evolution of the avian humerus and ulna appear to exhibit heterogeneous evolutionary patterns at different anatomical levels (external vs. internal morphology as well as epiphysis vs. diaphysis). This finding evinces a degree of mosaicism in the evolution of avian wing bone structure, which is emerging as a critical pattern in investigations of vertebrate phenotypic diversification (Barton and Harvey 2000; Bastir and Rosas 2009; Navalón et al. 2022).

#### 4.1.2. Effects of flightlessness and marine habits on wing bone structure

Most pairwise comparisons among eco-clusters that yielded significance concern two groups: eco- cluster C, i.e. marine aquatic predator birds, showing wing-propelled swimming (e.g., penguins), and eco- cluster H, i.e. flightless/poorly flying birds (but not including burst-flying taxa) (Table 3, Tables S13-S17). The size-corrected distribution of traits strongly contributing to DFs are significantly different for these groups, revealing that a few deviant taxa are predominantly driving these patterns.

The two flightless palaeognaths in our sample (the kiwi *Apteryx* and the emu *Dromaius*) are frequently set apart from most other taxa. Specifically, in the humerus, they clearly show higher epiphyseal trabecular node density (NodDen) (Fig. S5F-G, Fig. 10D-E), a pattern mirrored only by *Dromaius* in ulnar epiphyses (Fig. 4A, Fig. S11C-D, Fig. S13F-I; *Apteryx*’s ulnar epiphyseal data were discarded due to a very low number of trabeculae). Also, *Dromaius* and *Apteryx* show lower values for a suite of size-corrected humeral/ulnar diaphyseal traits, i.e. cross-sectional area (SC-CSA), second moments of area (SC-Imax and SC-Imin), polar section modulus (SC-Zpol) and total cross-sectional area (SC-TotAr), expected to positively relate to biomechanical loadings (Crowder and Stout 2011; Parsi-Pour and Kilbourne 2020) (Fig. S8). Thus, *Dromaius* and *Apteryx* have an epiphyseal structure characterized by an increased NodDen (*Dromaius* in humerus and ulna; *Apteryx* in the humerus, pending data on ulnar epiphyses); yet, surprisingly, this arrangement is combined with humeral/ulnar diaphyses that are less structurally able to resist mechanical loadings. Moreover, *Dromaius* is characterized by a distinctively lower fractal dimension (i.e. FD_prox,_ representing a proxy for the structural complexity of the trabecular network) relative to other taxa in our dataset.

FD has been hypothesised to inversely relate to mechanical loadings (Alfieri et al. 2025). Meanwhile, NodDen has been positively correlated to epiphyseal mechanical loading (Veneziano et al. 2021; Alfieri et al. 2025; Nguyen et al. *under review*, and references). Hence, high NodDen and low FD suggest increased mechanical loadings in the humeral epiphysis of *Apteryx* and *Dromaius*. This seemingly contrasts with the potential functional interpretation for the diaphyseal trait values which, instead, suggest anomalously low loadings relative to other taxa in our dataset. However, it should be noted that different loading regimes are expected to be dominant in distinct regions of long bones, i.e. bending/torsion in the diaphysis *vs.* axial in the epiphyses (Biewener et al. 1996; Pontzer et al. 2006; Carter and Beaupré 2007; Barak et al. 2011), and the typical loading regime of the avian forelimb is dominated by bending and torsion due to the constraints imposed by flight (de Margerie et al. 2005; Pennycuick 2008; Dumont 2010; Novitskaya et al. 2017; Sullivan et al. 2017). Therefore, if bending/torsion loadings characterize flight and this biomechanical regime is expected to dominate in the diaphysis, the flightless taxa in our dataset illustrate a release from flight-related constraints on wing internal bone diaphyseal structure. Accordingly, we expect to observe patterns similar to those of *Apteryx* and *Dromaius* in other terrestrial flightless taxa should they be sampled in future studies of a similar nature. A higher density of trabecular connections in the epiphyses of *Apteryx* and *Dromaius* is likely explained by relaxed selection pressures for lightweight bones, perhaps driven by the alleviation of the need to invest energy in bone resorption and mass-saving strategies. Similar explanations have been proposed for other tetrapods, such as the denser, more compact humeri and femora of slow arboreal mammals, which may be explained by relaxed selection pressures to decrease bone mass in these organisms that show an extraordinarily inactive and cautious lifestyle (Alfieri et al. 2023). The exceptionally low structural complexity (i.e., low FD), as well as the particularly thin trabeculae (i.e., low SC-Tb.Th; Fig. S5E) in the *Dromaius*’ humeral epiphyses are more difficult to fit within this framework, and we hope that the functional and evolutionary factors underlying these trends are explored in future work.

The other flightless taxa in our sample include the marine taxa *Nannopterum harrisi* (flightless cormorant) and *Spheniscus humboldti* (Humboldt penguin) which both exhibit patterns of convergence with the terrestrial flightless taxa for the humeral and ulnar traits discussed above (Fig. 4, Fig. S5-S13). Our data strongly suggest that flightlessness releases the wing bones from structural constraints regardless of marine or terrestrial contexts, resulting in the evolutionary acquisition of similar structural traits. Although this conclusion is drawn from only a small sample of flightless birds, the fact that *N. harrisi* exhibits these patterns and the flying cormorant in our dataset (*Phalacrocorax carbo*) does not, shows that functionally constrained morphologies may be altered rapidly once constraints related to maintenance of a functional flight apparatus are relaxed, as evidenced by the recent divergence time (∼7 Mya) of *N. harrisi* from flying cormorants (Burga et al. 2017). *Nannopterum* and, to a lesser extent, *Spheniscus* also exhibit similarly low values for ulnar diaphysis traits, similar to those of *Apteryx* and *Dromaius* discussed above (Fig. S12), and all four flightless taxa in our sample show higher ulnar diaphyseal global compactness (Cg) compared to virtually all the flighted taxa in our dataset (Fig. S4F). Although this trait has often been related to hydrostatic adaptations (see discussion below), it may also be indicative of flightlessness more generally. However, we make these interpretations cautiously because other observations from our data suggest that this scenario may be overly simplistic. Some puzzling patterns may be ascribed to issues of preservation of bone internal structure; for instance, the flying *Columba* yielded a high ulnar Cg, mirroring flightless taxa, but this pattern is only shown at the mid-diaphyseal level (i.e. ulnar Cg_50_), and not along the entire diaphysis (i.e. ulnar Cg_Aver_). This suggests that the high Cg_50_ of *Columba* may derive from the medullary cavity filled with other material, e.g. sediments, a condition that may artefactually and locally (i.e. only at mid-diaphysis) increase Cg. Yet, other patterns do not fit our interpretation and are consistently observed; for instance, the flying ducks *Anas* and *Oxyura*, and the swift *Chaetura*, as well as reluctant flying (but not flightless) taxa such as the malleefowl *Leipoa*, converge upon flightless birds in some structural traits (Fig. 4, Fig. S5-S13). Further work, including a larger taxon sample with closely related flying and flightless species from marine and terrestrial environments (e.g., from within Rallidae, Anatidae and Phalacrocoracidae) is warranted to explore these patterns in detail.

*Spheniscus* shows a distinctively more compact humeral diaphysis than other taxa, although other marine species share this feature to a lesser degree (e.g., the shearwater *Puffinus* and the auk *Uria*; Fig. 4B). High humeral compactness is not shown by terrestrial flightless birds within our sample and, thus, it likely represents an adaptation to a marine habitat, for example to increase ballast and decrease buoyancy (Habib 2010). Highly dense osteosclerotic bones in penguins have previously been attributed to their aquatic ecology, although this pattern was mainly explained through their thicker diaphyseal cortex (Habib 2010; Ksepka et al. 2015). *Spheniscus*, *Uria* and *Puffinus* in our sample exhibit high cortical thickness (SC-Ct.Th_50_) values (Table S1), and this trait is usually ecologically diagnostic (e.g., it may discriminate between aquatic, terrestrial and aerial taxa, Currey and Alexander 1985; Swartz et al. 1992). Thicker bone cortex in wing-propelled divers including *Spheniscus*, *Uria* and *Puffinus* appears to be a common convergent trait in both flightless and flying taxa (Habib 2010; Smith and Clarke 2014), and the ecological pressure exerted by swimming mechanics on wing bone compactness may outweigh that of flight. Notably, the increase of bone compactness has been related to decreased pneumaticity (Currey and Alexander 1985; Gutzwiller et al. 2013; Burton et al. 2023, 2025; Field et al. 2025). These aspects are tightly interacting, whereby bones with low pneumaticity show thicker cortices and are more compact compared to pneumatized bones, and this condition is common in aquatic taxa (Gutzwiller et al. 2013; Burton et al. 2023), with functional implications related to buoyancy. Although pneumatization and ecology are related, the latter is not the only factor determining whether a bone is pneumatic or apneumatic (Burton et al. 2023). Beyond the tentative explanation related to weakened pressure for bone mass-saving in the case of flightlessness (see above), it is also possible that the relatively high compactness in some non-aquatic taxa may be related to relatively low degrees of pneumaticity. Future studies addressing ulnar pneumaticity in these taxa may clarify this question.

Another trait for which *Spheniscus* and *Uria* (see also others in Watanabe et al. 2020) and, to a lesser extent, *Puffinus* morphologically converge is a higher value for humeral cross-sectional shape (CSS), indicative of elliptical cross-sections, with *Spheniscus* showing an extraordinarily high value (Fig. S8M-N, S9L-M). This is not surprising—the flattened forelimb bones of penguins are well-known (e.g., 37) and cross-sections from such a peculiarly shaped humerus are highly elliptical. Auks and shearwaters have been reported as also exhibiting relatively flattened forelimb elements (Habib 2010), justifying their relatively higher humeral CSS converging with penguins (Fig. S8M-N, S9L-M). Structurally flattened wing bones are associated with forelimb-propelled marine diving (Habib 2010). This locomotor behavior involves the use of the forelimb as a hydrofoil and is optimized in penguins, which exclusively forage via forelimb-propelled marine diving, unlike other extant diving birds that still retain aerial flight, e.g., auks (although multiple extinct lineages of flightless auks are known from the fossil record, Watanabe et al. 2020), and shearwaters. Water is more dense than air, and the fact that wing-propelled locomotion produces thrust during both upstroke and downstroke (with substantially higher power stroke frequencies) subjects the wing to high forelimb loadings (Lovvorn and Liggins 2002; Habib 2010). In this regard, flattened shapes and elliptical diaphyseal cross-sections would cause a tapered leading edge of wing bones, which, in turn, is potentially linked to the generation of substantial thrust during both upstroke and downstroke (Habib 2010). In addition to *Spheniscus*, *Uria*, and *Puffinus*, six other taxa in our sample are occasionally capable of performing wing-propelled swimming, while being capable of flying like auks and shearwaters, but they do not show any pattern of higher CSS (S9L-M). Four of them, i.e. the wader *Tringa*, the oystercatcher *Haematopus* and the kingfishers *Alcedo* and *Choloroceryle*, live in wetland, coastal or riverine habitats (Table S2). Hence, the fact that none of them mirror penguins, auks and shearwaters for living in marine habitats seems to suggest that only extreme specialisations to forelimb- propelled swimming in marine environments are associated with elliptical forelimb bone cross-sections. However, the fact that the other two wing-propelled swimming taxa—the petrel *Pelecanoides* and the gannet *Morus*—do not exhibit elliptical forelimb bone sections despite inhabiting marine environments (with *Pelecanoides* exhibiting a similar aquatic lifestyle to that of auks), suggests that the functional interpretation behind higher humeral CSS is likely more complex and warrants further investigation.

A relatively high CSS is also found in some hummingbirds, but this trait appears to be attributable to their unusual overall humeral morphology, characterized by a dramatically shortened humeral shaft, which is rendered particularly robust by enlarged articular processes (Zusi 2013). CSS becomes higher as cross-sections deviate from circularity, and prominent articular processes along the shaft will alter the diaphyseal shape so that it strongly deviates from a tubular model, yielding highly elliptical sections. Although high CSS is often related to stronger directional loadings (Alfieri et al. 2022 and references), if it is measured on sections extracted from diaphyses with prominent articular processes (as in hummingbirds), it will not be interpretable in this way; that is, the sections in hummingbirds appear less circular as a result of autapomorphic external morphological features, not as a result of inner structural adaptations (Patel et al. 2013).

#### 4.1.2. Characterisation of ecological diversity

In this work we analyzed ecological diversity through discrete categories, dubbed eco-clusters, capturing various aspects of avian ecological variation: flight style, habitat, primary lifestyle, migration, trophic level and trophic niche. In doing so, we followed the rationale of previous studies of bird wing anatomy (e.g. Baumgart et al. 2021), in order to avoid restricting functional hypotheses to flight style alone, as has also been done (e.g. Baliga et al. 2019; Lowi-Merri et al. 2021). Interestingly, previous work, whether exclusively focusing on flight style or not, followed different approaches to describe ecological variability (e.g., using presence/absence behavioral variables in linear models, e.g. Lowi-Merri et al. 2021), or employing discrete categories, either literature-based (Habib and Ruff 2008; Baumgart et al. 2021) or derived from clustering techniques (Baliga et al. 2019). This methodological heterogeneity probably reflects limitations in all approaches when ecological diversity needs to be classified for functional and evolutionary morphological studies. The eco-clusters that we identified successfully summarized extensive information in one categorical variable. Moreover, this ecological classification enables studying eco-morphological convergence (Fig. 3C and above), and as such it may provide a promising framework for future studies. However, it should be noted that in our ecological classification system, a portion of ecological variance is unavoidably lost (i.e., not all the taxa assigned to an eco-cluster exhibit all of its defining features, Fig. S4). Hence, the ecological classification system that we used, based on both cluster analysis and adjustments that we performed (Section S8) can be improved (e.g., by including additional ecological data, or using alternative clustering techniques). Also, using presence/absence variables (e.g. Lowi-Merri et al. 2021) overcomes this limitation but necessitates the application of complex models, in which many factors and co-variates (potentially interacting with one another) are involved. Quantitative behavioral data (e.g., relative frequencies of particular behaviors) may represent a profitable way forward in order to attempt to mitigate these issues in the future, as has been done in mammals (Granatosky 2018).

## 5. Conclusions

Studying the inner structure of wing skeletal elements in a phylogenetically broad sample of 140 living bird taxa, we found that the main ecological variable contributing to wing internal bone structural variation is flight. Overall, we find that internal bone structure in the humerus and ulna of birds exhibits low disparity overall, and the variability that exists is predominantly driven by body mass. We propose that the functional and mechanical demands of flight are essentially universal across bird phylogeny regardless of specific flight behaviors (e.g., hovering, burst flight, soaring). We interpret this relatively homogeneous and conserved configuration as reflecting widely recognised aspects of the structural design of avian wing bones, i.e. elements that are hollow, with low mass, exhibiting a thin but compact cortex, struts/ridges internally, and circular cross-sections—an architectural arrangement that enhances resistance to bending and torsional loadings. Once this general configuration was established in the ancestry of crown group birds, most flying birds appear to have retained this structure irrespective of most other ecological factors, which collectively appear to have minimal effects on internal bone structure, with the exception of wing-propelled aquatic divers. Indeed, some wing-propelled diving birds exhibit distinctively higher global compactness and ellipticity of their humeral diaphyses, features plausibly reflecting hydrostatic and hydrodynamic adaptations. Also, we found that both terrestrial and aquatic flightless taxa converge in some bone structural traits — e.g., higher trabecular density of connections, lower diaphyseal polar section modulus — suggesting that as birds are released from the ecological and mechanical constraints imposed by flight, the internal structure of the wing bones undergo predictable forms of modification.

Although wing bone inner structure generally follows the same macroevolutionary patterns as identified in investigations of external wing bone morphology (Serrano et al. 2020), our findings open avenues for further work aiming to elucidate how wing bone structural traits adapt to loss of flight, as well as how ecological signal varies between wing bone inner structure *vs*. external shape, and between wing long bone epiphyses and diaphyses. Interestingly, the latter question may contribute to identifying differential evolutionary patterns for different traits within the same skeletal element, enabling novel explorations of evolutionary mosaicism. Finally, our work should improve estimates of the flying ability of fossil birds from preserved wing bones and thereby facilitate more accurate ecological reconstructions of fossil bird taxa.

## Supporting information

It contains all supplementary methods and results (excluding Figs S5-S13, which are shown separately in pdf, see below)

Results for proximal humeral structural parameters.

Traitgrams for proximal humeral parameters of interest (identified as detailed in the Main Text)

Results for humeral diaphyseal structural parameters.

Further boxplots of humeral diaphyseal parameters of interest (identified as detailed in the Main Text)

Traitgrams for humeral diaphyseal parameters of interest (identified as detailed in the Main Text)

Results for distal humeral structural parameters.

Results for proximal ulnar structural parameters.

Results for ulnar diaphyseal structural parameters.

Results for distal ulnar structural parameters.

Excel file with raw results for morphological data

Excel file with raw results for ecological data

## Acknowledgements

For allowing us to collect and/or CT-scan museum specimens (both through the MorphoSource ‘Bird Tempo’, https://www.morphosource.org/projects/00000C420, and the ‘Florida Museum of Natural History: Vertebrate Paleontology: Microfossils’, https://www.morphosource.org/projects/00000C580?locale=en, projects and newly acquired specimens) we thank: Roger Benson (American Museum of Natural History, New York, USA), Edward Stanley (Florida Museum of Natural History), Mathew Lowe, Mike Brooke and Keturah Smithson (University of Cambridge, UK), Judith White, Jo Cooper, Mike Day, Brett Clark, Vincent Fernandez and Agnese Lanzetti (Natural History Museum, London, UK), Stephanie Lechki (University of Oxford, UK), Mark Carnall and Eileen Westwig (Oxford University Museum of Natural History, Oxford, UK), Janet Hinshaw and Matt Friedman (University of Michigan, Museum of Zoology, Ann Arbour, Michigan, USA), Kristof Zyskowski (Yale Peabody Museum, New Haven, Connecticut, USA), Ben Marks and John Bates (Field Museum of Natural History, Chicago, USA), Tom Davies, Ben Moon and Liz Martin-Silverstone (University of Bristol), April Neander and Zhe-Xi Luo (University of Chicago PaleoCT). Also, we thank Alessio Veneziano (Archéozoologie, Archéobotanique: Sociétés, Pratiques et Environnements (AASPE), Muséum National d’Histoire Naturelle, CNRS, Paris, France) and Vikram Baliga (University of British Columbia) for helping on R functions for bone and cluster analysis. This work is part of the project BE- BOST, financed by the grant TMPFP3_217022 awarded to F.A. by the Swiss National Science Foundation (https://www.snf.ch; SNSF Swiss Postdoctoral Fellowships, SPF). Additional funding was provided by UKRI grant MR/X015130/1 to DJF. For the purpose of open access, the authors have applied a Creative Commons Attribution (CC BY) licence to any Author Accepted Manuscript version arising. EMS was supported by a Sarah Woodhead Research Fellowship (Girton College, Cambridge) and OED was supported by an EAVP Research Grant.

## Author contributions

F.A. collected the sample, acquired, processed, and analysed μCT data, performed statistical analyses, and drafted the manuscript. O.E.D. built the composite phylogeny and, together with E.M.S. and F.A. collected the ecological data. F.A., O.E.D., E.M.S., A.-C.F and D.J.F. conceived the study, interpreted the data, and contributed to the writing and editing of the manuscript.

## Data availability statement

Raw morphological and ecological datasets, additional methods and additional results are available as Supplementary Materials.

## Data sources

Data sources exploited to extract morphological (i.e. CT-scans downloaded from MorphoSource) and ecological data (i.e. databases and literature) are listed in Materials and Methods, related Supplementary Sections and Tables.

## References

1. Alfieri, F., L. Botton-Divet, J. A. Nyakatura, and E. Amson. 2022. Integrative approach uncovers new patterns of ecomorphological convergence in slow arboreal xenarthrans. J Mamm Evol, doi: 10.1007/s10914-021-09590-5.

2. Alfieri, F., L. Botton-Divet, J. Wölfer, J. A. Nyakatura, and E. Amson. 2023. A macroevolutionary common-garden experiment reveals differentially evolvable bone organization levels in slow arboreal mammals. Communications Biology 6(1):995.

3. Alfieri, F., J. A. Nyakatura, and E. Amson. 2021. Evolution of bone cortical compactness in slow arboreal mammals. Evolution 75:542–554.

4. Alfieri, F., A. Veneziano, D. Panetta, P. A. Salvadori, E. Amson, and D. Marchi. 2025. The relationship between primate distal fibula trabecular architecture and arboreality, phylogeny and size. Journal of Anatomy 00:1–29.

5. Amson, E., and F. Bibi. 2021. Differing effects of size and lifestyle on bone structure in mammals. BMC Biol 19:87.

6. Baliga, V. B., I. Szabo, and D. L. Altshuler. 2019. Range of motion in the avian wing is strongly associated with flight behavior and body mass. Science advances 5(10):eaaw6670.

7. Barak, M. M., D. E. Lieberman, and J.-J. Hublin. 2011. A Wolff in sheep’s clothing: trabecular bone adaptation in response to changes in joint loading orientation. Bone 49:1141–1151.

8. Barrowclough, G. F., J. Cracraft, J. Klicka, and R. M. Zink. 2016. How many kinds of birds are there and why does it matter? PLoS ONE 11:e0166307.

9. Barton, R. A., and P. H. Harvey. 2000. Mosaic evolution of brain structure in mammals. Nature 405:1055–1058.

10. Bastir, M., and A. Rosas. 2009. Mosaic evolution of the basicranium in *Homo* and its relation to modular development evolutionary biology. Evol. Biol. 36:57–70.

11. Basu, C., A. M. Wilson, and J. R. Hutchinson. 2019. The locomotor kinematics and ground reaction forces of walking giraffes. J Exp Biol 222:jeb159277.

12. Baumgart, S. L., P. C. Sereno, and M. W. Westneat. 2021. Wing shape in waterbirds: Morphometric patterns associated with behavior, habitat, migration, and phylogenetic convergence. Integr. Org Biol 3 (1).

13. Beauchamp, G. 2023. Is wing morphology across birds associated with life history and sociality? Frontiers in Bird Science 2:1305453.

14. Biewener, A. A. 2011. Muscle function in avian flight: achieving power and control. Philos. Trans. R. Soc. B: Biol. Sci. 366:1496–1506.

15. Biewener, A. A., N. L. Fazzalari, D. D. Konieczynski, and R. V. Baudinette. 1996. Adaptive changes in trabecular architecture in relation to functional strain patterns and disuse. Bone 19:1–8.

16. Billerman, S. M., B. K. Keeney, G. M. Kirwan, F. Medrano, N. D. Sly, and M. G. Smith, Editors (2025). Birds of the World. Cornell Laboratory of Ornithology, Ithaca, NY, USA. 10.2173/bow

17. Bjarnason, A., and R. . B. . J. Benson. 2021. A 3D geometric morphometric dataset quantifying skeletal variation in birds. MorphoMuseuM 7(1).

18. Burga, A., W. Wang, E. Ben-David, P. C. Wolf, A. M. Ramey, C. Verdugo, K. Lyons, P. G. Parker, and L. Kruglyak. 2017. A genetic signature of the evolution of loss of flight in the Galapagos cormorant. Science 356:eaal3345.

19. Burton, M. G. P., R. B. Benson, and D. J. Field. 2023. Direct quantification of skeletal pneumaticity illuminates ecological drivers of a key avian trait. Proceedings of the Royal Society B 290(1995):20230160.

20. Burton, M. G. P., K. Mellor, E. Smith, J. Benito, E. Martin-Silverstone, P. O’Connor, and D. J. Field. 2025. The influence of soft tissue volume on estimates of skeletal pneumaticity: implications for fossil archosaurs. Philosophical Transactions of the Royal Society 380:20230428.

21. Carter, D. R., and G. S. Beaupré. 2007. Skeletal Function and Form: Mechanobiology of Skeletal Development, Aging, and Regeneration. Cambridge: Cambridge University Press.

22. Chen, A., E. M. Steell, R. B. J. Benson, and D. J. Field. 2025. Towards a comprehensive anatomical matrix for crown birds: phylogenetic insights from the pectoral girdle and forelimb skeleton. bioRxiv.

23. Chinsamy, A. 2023. Palaeoecological deductions from osteohistology. Biology Letters 19(8):20230245.

24. Chinsamy, A., D. Angst, A. Canoville, and U. B. Göhlich. 2020. Bone histology yields insights into the biology of the extinct elephant birds (Aepyornithidae) from Madagascar. Biological Journal of the Linnean Society 130, Issue 2:268–295.

25. Chirchir, H., H. Dean, K. Carlson, and C. Ruff. 2017. Revisiting the Evolution of Low Trabecular Bone Density in Modern Humans. The FASEB Journal 31:251.3-251.3. Federation of American Societies for Experimental Biology.

26. Clavel, J., G. Escarguel, and G. Merceron. 2015. mvmorph: an R package for fitting multivariate evolutionary models to morphometric data. Methods Ecol Evol 6:1311–1319.

27. Cotter, M. M., S. W. Simpson, B. M. Latimer, and C. J. Hernandez. 2009. Trabecular microarchitecture of hominoid thoracic vertebrae. Anat Rec 292:1098–1106.

28. Crowder, C., and S. Stout. 2011. Bone Histology: An Anthropological Perspective. CRC Press, Boca Raton.

29. Cubo, J., and A. Casinos. 1998. Biomechanical significance of cross-sectional geometry of avian long bones. European Journal of Morphology 36(1).

30. Currey, J., and R. M. Alexander. 1985. The thickness of the walls of tubular bones. Journal of Zoology 206:453–568.

31. Currey, J. D. 2002. Bones: Structure and Mechanics. Princeton University Press.

32. De Margerie, E. 2002. Laminar bone as an adaptation to torsional loads in flapping flight. J Anat 201:521– 526.

33. de Margerie, E., S. Sanchez, J. Cubo, and J. Castanet. 2005. Torsional resistance as a principal component of the structural design of long bones: comparative multivariate evidence in birds. Anat Rec A Discov Mol Cell Evol Biol 282:49–66.

34. Dial, K. P. 2003. Evolution of avian locomotion: correlates of flight style, locomotor modules, nesting biology, body size, development, and the origin of flapping flight. Auk 120:941–952.

35. Domander, R., A. A. Felder, and M. Doube. 2021. BoneJ2 - refactoring established research software. Wellcome Open Research. 6:37.

36. Doube, M., M. M. Klosowski, A. M. Wiktorowicz-Conroy, J. R. Hutchinson, and S. J. Shefelbine. 2011. Trabecular bone scales allometrically in mammals and birds. Proc. Biol. Sci. 278:3067–3073.

37. Dumont, E. R. 2010. Bone density and the lightweight skeletons of birds. Proceedings of the Royal Society B: Biological Sciences 277:2193–2198. Royal Society.

38. Field, D. J., M. G. P. Burton, J. Benito, O. J. C. Plateau, and G. Navalón. 2025. Whence the birds: 200 years of dinosaurs, avian antecedents. Biology Letters 21.

39. Field, D. J., C. Lynner, C. Brown, and S. A. F. Darroch. 2013. Skeletal Correlates for Body Mass Estimation in Modern and Fossil Flying Birds. PLoS ONE 8:e82000.

40. Frongia, G. N., M. Muzzeddu, P. Mereu, G. Leoni, F. Berlinguer, M. Zedda, V. Farina, V. Satta, M. Di Stefano, and S. Naitana. 2018. Structural features of cross_sectional wing bones in the griffon vulture (*Gyps fulvus*) as a prediction of flight style. Journal of Morphology 279:1753–1763.

41. Gill, F. B. 2007. Ornithology, 3rd edition. W.H. Freeman and Company, London.

42. Gônet, J., M. Laurin, and J. R. Hutchinson. 2023. Evolution of posture in amniotes–Diving into the trabecular architecture of the femoral head. Journal of Evolutionary Biology 36(8):1150–1165.

43. Granatosky, M. C. 2018. A review of locomotor diversity in mammals with analyses exploring the influence of substrate use, body mass and intermembral index in primates. J. Zool 306:207–216.

44. Gutzwiller, S. C., A. Su, and P. M. O’Connor. 2013. Postcranial pneumaticity and bone structure in two clades of neognath birds. The Anatomical Record 296(6):867–876.

45. Habib, M. 2010. The structural mechanics and evolution of aquaflying birds. Biological Journal of the Linnean Society 99(4):687–698.

46. Habib, M. B., and C. B. Ruff. 2008. The effects of locomotion on the structural characteristics of avian limb bones. Zoological Journal of the Linnean Society 153:601–624.

47. Jablonski, D. 2022. Evolvability and macroevolution: overview and synthesis. Evol Biol 49:265–291.

48. Kish, T. M. 2011. Study of the microstructure and mechanical properties of hummingbird wing-bones. Diss. Massachusetts Institute of Technology.

49. Kitchell, J. A. 1985. Evolutionary paleoecology: recent contributions to evolutionary theory. Paleobiology 11:91–104.

50. Kivell, T. L. 2016. A review of trabecular bone functional adaptation: what have we learned from trabecular analyses in extant hominoids and what can we apply to fossils? J. Anat. 228:569–594.

51. Ksepka, D. T., S. Werning, M. Sclafani, and Z. M. Boles. 2015. Bone histology in extant and fossil penguins (Aves: Sphenisciformes). Journal of Anatomy 227(5):611–630.

52. Lieberman, D. E. 1997. Making behavioral and phylogenetic inferences from hominid fossils: considering the developmental influence of mechanical forces. Annual Review of Anthropology 26:185–210.

53. Louis, L. D., R. C. K. Bowie, and R. Dudley. 2022. Wing and leg bone microstructure reflects migratory demands in resident and migrant populations of the Dark_eyed Junco (*Junco hyemalis*). Ibis 164.1:132–150.

54. Lovvorn, J., and G. Liggins. 2002. Interactions of body shape, body size, and stroke-acceleration patterns in costs of underwater swimming by birds. Functional Ecology 16:106–112.

55. Lowi-Merri, T. M., R. B. J. Benson, S. Claramunt, and D. C. Evans. 2021. The relationship between sternum variation and mode of locomotion in birds. BMC Biol 19:165.

56. Maderspacher, F. 2022. Flightless birds. Current Biology R1155–R1162.

57. Maechler, M., P. Rousseeuw, A. Struyf, M. Hubert, and K. Hornik. 2022. cluster: Cluster Analysis Basics and Extensions. R package version 2.1.4.

58. Mitchell, J. 2016. Cortical Bone Remodeling in Amniota. A Functional, Evolutionary and Comparative Perspective of Secondary Osteons. Bonn, Germany.

59. Navalón, G., A. Bjarnason, E. Griffiths, and R. B. Benson. 2022. Environmental signal in the evolutionary diversification of bird skeletons. Nature 611(7935):306–311.

60. Nguyen, U., F. Alfieri, A. Licht, and J. A. Nyakatura. n.d. Trabecular structure correlates with leaping distance in tamarins. Under review.

61. Norberg, U. M. 1990. Vertebrate Flight. Springer-Verlag, Berlin.

62. Novitskaya, E., C. J. Ruestes, M. M. Porter, V. A. Lubarda, M. A. Meyers, and J. McKittrick. 2017. Reinforcements in avian wing bones: Experiments, analysis, and modeling. Journal of the Mechanical Behavior of Biomedical Materials 76:85–96.

63. Nudds, R. L., G. J. Dyke, and J. M. V. Rayner. 2007. Avian brachial index and wing kinematics: putting movement back into bones. Journal of Zoology 272:218–226.

64. Padian, K. 2011. Vertebrate palaeohistology then and now: A retrospective in the light of the contributions of Armand de Ricqlès. Comptes Rendus Palevol 10:303–309.

65. Parsi-Pour, P., and B. M. Kilbourne. 2020. Functional morphology and morphological diversification of hind limb cross-sectional traits in mustelid mammals. Integr. Org. Biol. 2:obz032.

66. Patel, B. A., C. B. Ruff, E. L. R. Simons, and J. M. Organ. 2013. Humeral cross-sectional shape in suspensory primates and sloths. Anat Rec (Hoboken) 296:545–556.

67. Pennycuick, C. J. 2008. Modelling the flying bird. Elsevier.

68. Pinheiro, J., D. Bates, S. DebRoy, and D. Sarkar. 2020. nlme: Linear and Nonlinear Mixed Effects Models. R package version 3.1–147.

69. Plasse, M., E. Amson, J. Bardin, Q. Grimal, and D. Germain. 2019. Trabecular architecture in the humeral metaphyses of non_avian reptiles (Crocodylia, Squamata and Testudines): lifestyle, allometry and phylogeny. Journal of Morphology 280:982–998.

70. Pontzer, H., D. E. Lieberman, E. Momin, M. J. Devlin, J. D. Polk, B. Hallgrímsson, and D. M. L. Cooper. 2006. Trabecular bone in the bird knee responds with high sensitivity to changes in load orientation. J Exp Biol 209:57–65. The Company of Biologists Ltd.

71. R Core Team. 2024. R: A language and environment for statistical computing. R Foundation for Statistical Computing, Vienna, Austria.

72. Rayner, J. M. V. 1988. Form and Function in Avian Flight. Pp. 1–66 *in* Current Ornithology. Johnston R. (ed), Boston, Springer.

73. Revell, L. J. 2012. phytools: an R package for phylogenetic comparative biology (and other things). Methods Ecol. Evol. 3:217–223.

74. Ruff, C., B. Holt, and E. Trinkaus. 2006. Who’s afraid of the big bad Wolff?: ‘Wolff’s law’ and bone functional adaptation. Am. J. Phys. Anthropol. 129:484–498.

75. Ryan, T. M., and C. N. Shaw. 2012. Unique suites of trabecular bone features characterize locomotor behavior in human and non-human anthropoid primates. PLoS ONE 7:e41037.

76. Schindelin, J., I. Arganda-Carreras, E. Frise, V. Kaynig, M. Longair, T. Pietzsch, S. Preibisch, C. Rueden, S. Saalfeld, B. Schmid, J.-Y. Tinevez, D. J. White, V. Hartenstein, K. Eliceiri, P. Tomancak, and A. Cardona. 2012. Fiji: an open-source platform for biological-image analysis. Nat. Methods 9:676–682.

77. Serrano, F. J., M. Costa-Pérez, G. Navalón, and A. Martín-Serra. 2020. Morphological Disparity of the Humerus in Modern Birds. Diversity 12:173.

78. Simons, E. L. R., T. L. Hieronymus, and P. M. O’Connor. 2011. Cross sectional geometry of the forelimb skeleton and flight mode in pelecaniform birds. J. Morphol. 272:958–971.

79. Smith, N. A., and J. A. Clarke. 2014. Osteological histology of the Pan_Alcidae (Aves, Charadriiformes): correlates of wing_propelled diving and flightlessness. The Anatomical Record 297(2):188–199.

80. Stiller, J., S. Feng, A.-A. Chowdhury, I. Rivas-González, D. A. Duchêne, Q. Fang, Y. Deng, A. Kozlov, A. Stamatakis, S. Claramunt, J. M. T. Nguyen, S. Y. W. Ho, B. C. Faircloth, J. Haag, P. Houde, J. Cracraft, M. Balaban, U. Mai, G. Chen, R. Gao, C. Zhou, Y. Xie, Z. Huang, Z. Cao, Z. Yan, H. A. Ogilvie, L. Nakhleh, B. Lindow, B. Morel, J. Fjeldså, P. A. Hosner, R. R. da Fonseca, B. Petersen, J. A. Tobias, T. Székely, J. D. Kennedy, A. H. Reeve, A. Liker, M. Stervander, A. Antunes, D. T. Tietze, M. F. Bertelsen, F. Lei, C. Rahbek, G. R. Graves, M. H. Schierup, T. Warnow, E. L. Braun, M. T. P. Gilbert, E. D. Jarvis, S. Mirarab, and G. Zhang. 2024. Complexity of avian evolution revealed by family-level genomes. Nature 629:851–860. Nature Publishing Group.

81. Sullivan, T. N., B. Wang, H. D. Espinosa, and M. A. Meyers. 2017. Extreme lightweight structures: avian feathers and bones. Materials Today 20:377–391.

82. Swartz, S. M., M. B. Bennet, and D. R. Carrier. 1992. Wing bone stresses in free flying bats and the evolution of skeletal design for flight. Nature 359:726–729.

83. Tibshirani, R., G. Walther, and T. Hastie. 2001. Estimating the number of clusters in a data set via the gap statistic. J R Stat Soc Series B Stat Methodol. 63:411–23.

84. Tobalske, B. W. 2016. Avian Flight. Pp. 149–166 *in* Handbook of Bird Biology. John Wiley & Sons, Lovette, I. J., Fitzpatrick, J. W. (Eds.).

85. Tobias, J. A., C. Sheard, A. L. Pigot, A. J. M. Devenish, J. Yang, F. Sayol, M. H. C. Neate-Clegg, N. Alioravainen, T. L. Weeks, R. A. Barber, P. A. Walkden, H. E. A. MacGregor, S. E. I. Jones, C. Vincent, A. G. Phillips, N. . M. Marples, F. A. Montaño-Centellas, V. Leandro-Silva, S. Claramunt, B. Darski, B. G. Freeman, T. P. Bregman, C. R. Cooney, E. C. Hughes, E. J. R. Capp, Z. K. Varley, N. R. Friedman, H. Korntheuer, A. Corrales-Vargas, C. H. Trisos, B. C. Weeks, D. M. Hanz, T. Töpfer, G. A. Bravo, V. Remeš, L. Nowak, L. S. Carneiro, A. J. Moncada R., B. Matysioková, D. T. Baldassarre, A. Martínez-Salinas, J. D. Wolfe, P. M. Chapman, B. G. Daly, M. C. Sorensen, A. Neu, M. A. Ford, R. J. Mayhew, L. Fabio Silveira, D. J. Kelly, N. N. D. Annorbah, H. S. Pollock, A. M. Grabowska-Zhang, J. P. McEntee, J. Carlos T. Gonzalez, C. G. Meneses, M. C. Muñoz, L. L. Powell, G. A. Jamie, T. J. Matthews, O. Johnson, G. R. R. Brito, K. Zyskowski, R. Crates, M. G. Harvey, M. Jurado Zevallos, P. A. Hosner, T. Bradfer-Lawrence, J. M. Maley, F. G. Stiles, H. S. Lima, K. L. Provost, M. Chibesa, M. Mashao, J. T. Howard, E. Mlamba, M. A. H. Chua, B. Li, M. I. Gómez, N. C. García, M. Päckert, J. Fuchs, J. R. Ali, E. P. Derryberry, M. L. Carlson, R. C. Urriza, K. E. Brzeski, D. M. Prawiradilaga, M. J. Rayner, E. T. Miller, R. C. K. Bowie, R.-M. Lafontaine, R. P. Scofield, Y. Lou, L. Somarathna, D. Lepage, M. Illif, E. L. Neuschulz, M. Templin, D. M. Dehling, J. C. Cooper, O. S. G. Pauwels, K. Analuddin, J. Fjeldså, N. Seddon, P. R. Sweet, F. A. J. DeClerck, L. N. Naka, J. D. Brawn, A. Aleixo, K. Böhning-Gaese, C. Rahbek, S. A. Fritz, G. H. Thomas, and M. Schleuning. 2022. AVONET: morphological, ecological and geographical data for all birds. Ecology Letters 25:581–597.

86. Töpfer, T. 2018. Morphological Variation in Birds: Plasticity, Adaptation, and Speciation. Pp. 63–74 *in* Bird Species, Tietze D.T. (Ed). Springer Open, Basel, Switzerland.

87. Veneziano, A., M. Cazenave, F. Alfieri, D. Panetta, and D. Marchi. 2021. Novel strategies for the characterization of cancellous bone morphology: Virtual isolation and analysis. Am J Phys Anthropol ajpa.24272.

88. Wang, X., and J. A. Clarke. 2015. The evolution of avian wing shape and previously unrecognized trends in covert feathering. Proc. R. Soc. B. 282:20151935.

89. Watanabe, J., D. J. Field, and H. Matsuoka. 2020. Wing musculature reconstruction in extinct flightless auks (*Pinguinus* and *Mancalla*) reveals incomplete convergence with penguins (Spheniscidae) due to differing ancestral states. Integr. Org. Biol. obaa040.

90. Wyles, J. S., J. G. Kunkel, and A. C. Wilson. 1983. Birds, behavior, and anatomical evolution. PNAS 80(14):4394–4397.

91. Zusi, R. L. 2013. Introduction to the skeleton of hummingbirds (Aves: Apodiformes, Trochilidae) in functional and phylogenetic contexts. Ornithological Monographs 77.1:1–94.

