## Supplementary material for "Constrained variation in the internal architecture of avian wing bones": It contains all supplementary methods and results (excluding Figs S5-S13, which are shown separately in pdf, see below)

### Section S1

The μCT scans studied in this work were collected from three sources:

1. ‘TEMPO birds’ MorphoSource project (https://www.morphosource.org/projects/00000C420; project ID: 00000C420), referred to as ‘TEMPO birds’ in Tables S1 and S2;
2. ‘Florida Museum of Natural History: Vertebrate Paleontology: Microfossils’ MorphoSource project (https://www.morphosource.org/projects/00000C580?locale=en; project ID: 00000C580), referred to as ‘FMNH-VP’ in Tables S1 and S2;
3. Specimens sampled from the bird osteological collection of the Natural History Museum, Tring (UK) and μCT scanned at the Cambridge Biotomography Centre (Cambridge, UK), referred to as ‘NHMUK (Tring)’ in Tables S1 and S2;

### Section S2: bone orientation steps

As further detailed below, we first constrained the humeri, ulnae and femora positions in the *xy* plane, by orienting: the humeri with the point of maximum curvature of the head tangent to the *x*-axis; the ulnae with the semi-lunar notch tangent to the *x*-axis, the femora with the two dorsalmost points of the distal epiphysis tangent to the *x*-axis. Then, the mid-points of the proximal and the distal metaphyses were aligned along the *z*-axis, achieving a standard orientation for the three long bones

The steps followed to orient long bones in standard anatomical position are here summarized on the humerus, the ulna and the femur of *Anser fabalis* NHMUK 1895-2-6-9

**Humerus**

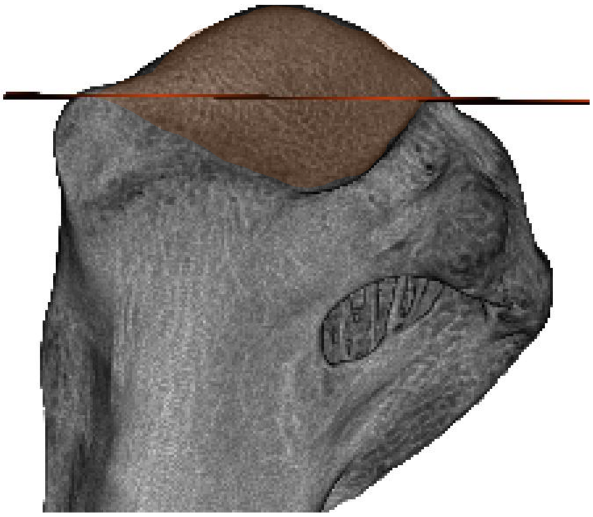

A. The humerus is observed in caudal view and the humeral head (highlighted in brown) proximodistal (PD) mid-level (around the 50% of the PD length of the head) is identified (here shown with a red plane)

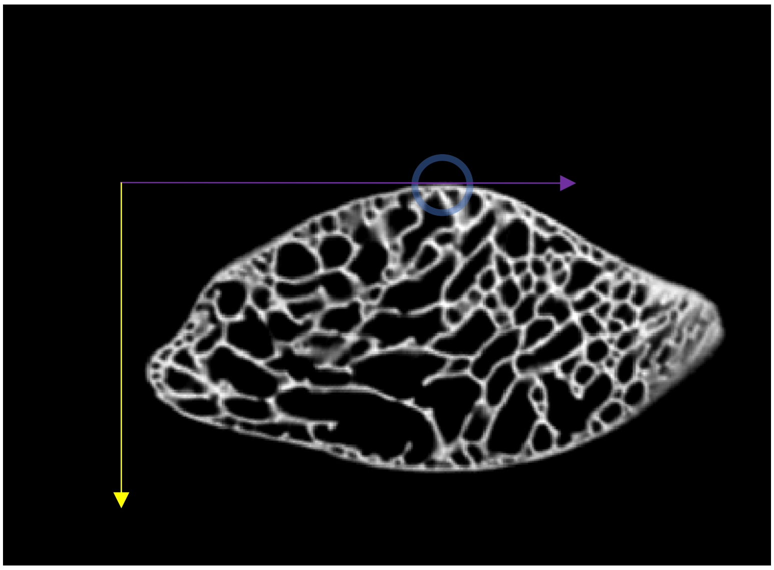

B. The cross-section corresponding to the humeral head PD mid-level (identified by the red plane in the step A) is here visualized. The humerus is rotated in the *xy* plane such that it has the point of maximum curvature of the head (on the caudal side, highlighted with a blue circle) tangent to the *x*-axis (shown in purple)

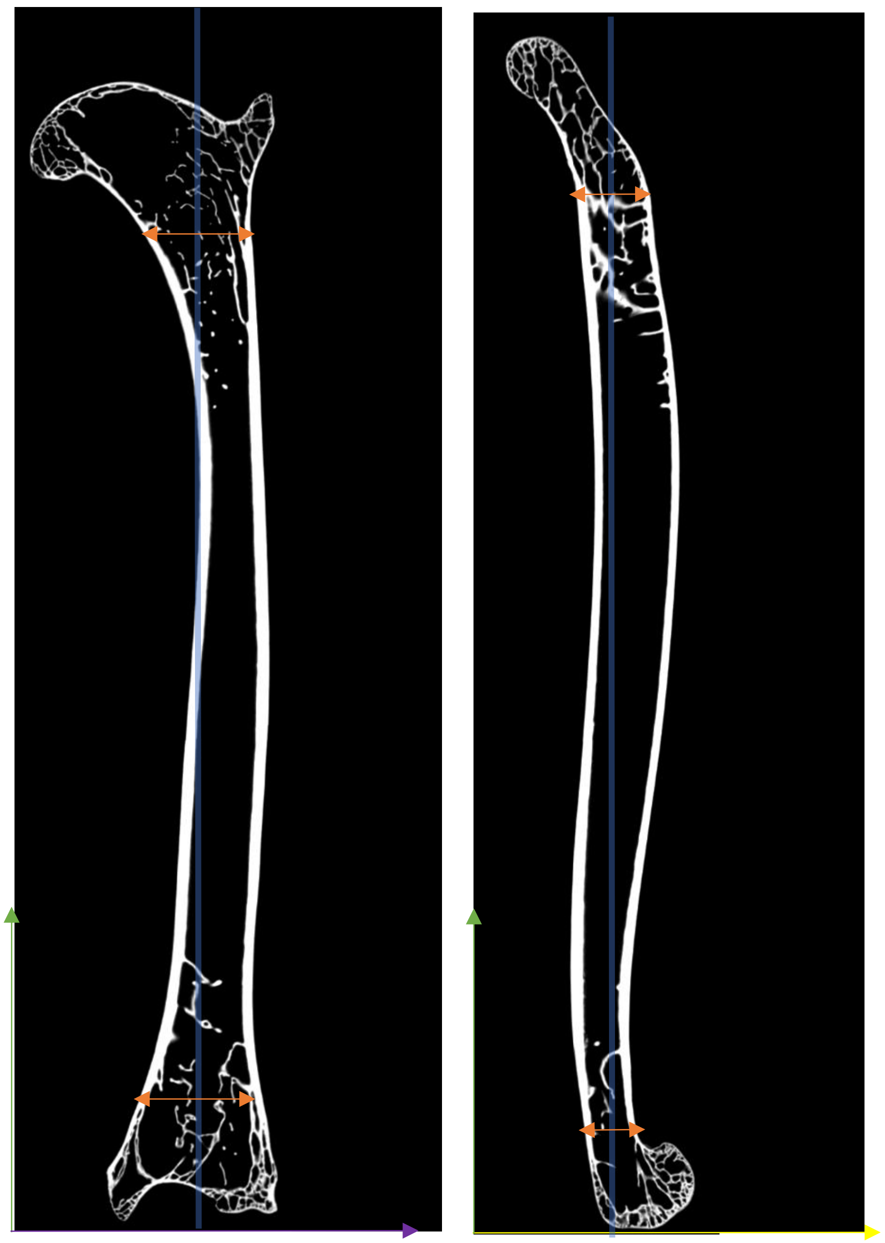

C. Humeral coronal (left) and sagittal (right) sections. Rotating the bone in the two views, the humerus is oriented in the *xz* and *yz* planes, respectively, by aligning the mid-points of the proximal and distal metaphyseal regions (i.e. transitional regions between epiphysis and diaphysis; shown with an orange double-headed arrow) such that they both lie on a line (shown in shaded blue) which is parallel to the *z*-axis (shown in green)

**Ulna**

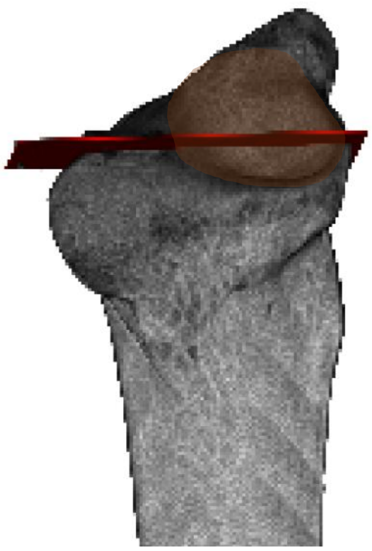

A. The semi-lunar notch (highlighted in brown) PD mid-level (around the 50% of the PD length of the notch) is identified (here shown with a red plane)

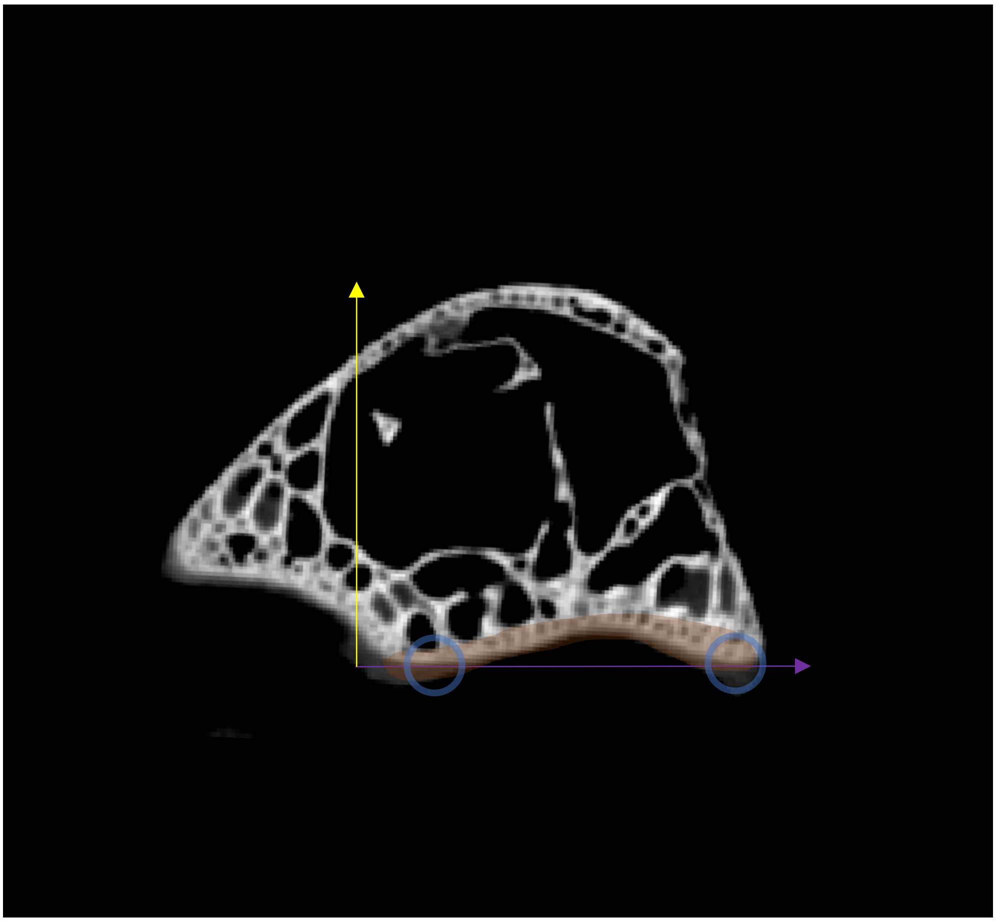

B. The cross-section corresponding to the semi-lunar notch PD mid-level (identified by the red plane in the step A) is here visualized. The ulna is rotated in the *xy* plane such that it has the *x*-axis contacting the two caudalmost points of the semi-lunar notch (points shown with blue circles, semi-lunar notch outlined in in orange).

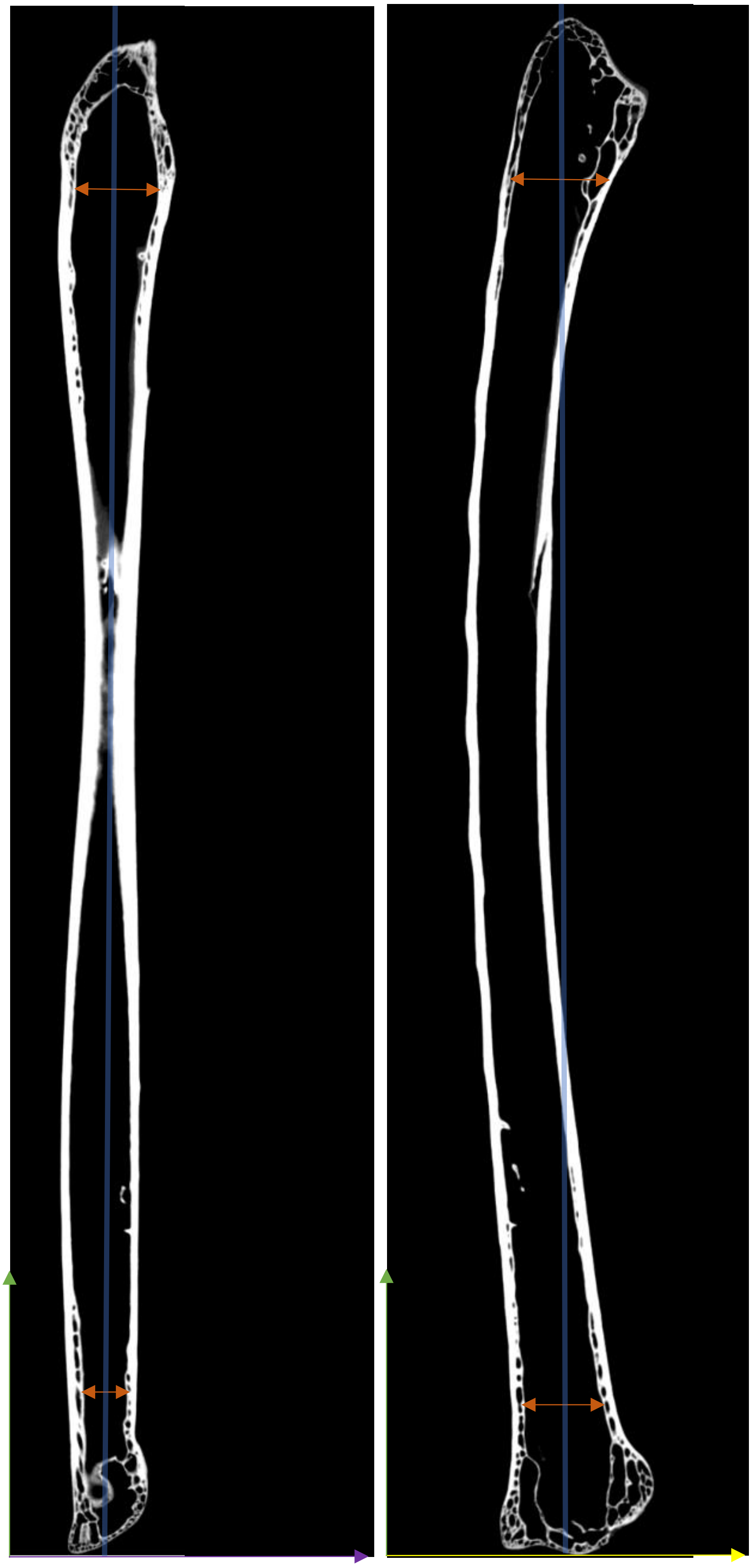

C. Ulnar coronal (left) and sagittal (right) sections. Rotating the bone in the two views, the ulna is oriented in the *xz* and *yz* planes, respectively, by aligning the mid-points of the proximal and distal metaphyseal regions (i.e. transitional regions between epiphysis and diaphysis; shown with an orange double-headed arrow) such that they both lie on a line (shown in shaded blue) which is parallel to the *z*-axis (shown in green)

**Femur**

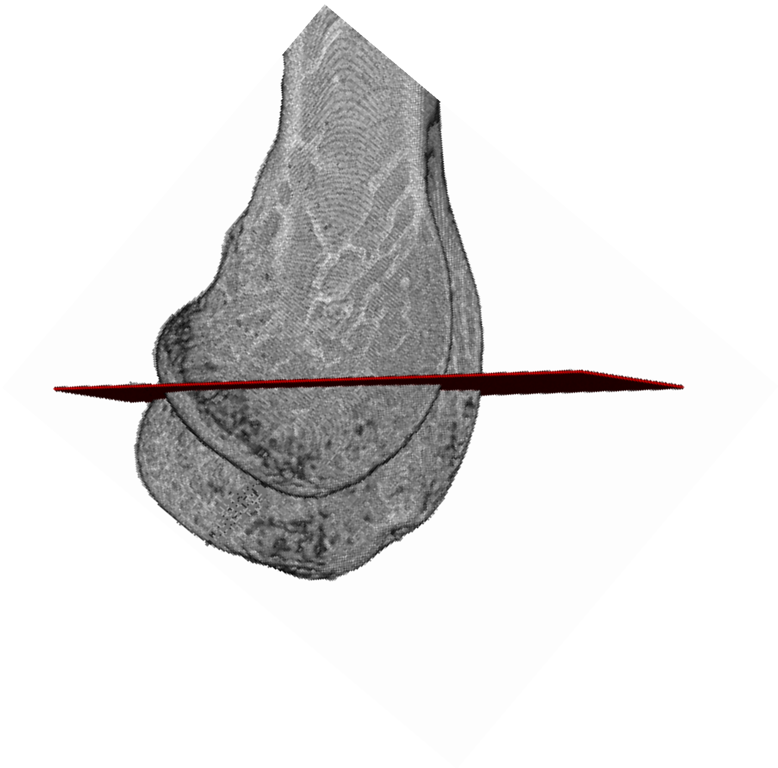

A. The PD mid-level (around the 50% of the PD length) of the femur medial condyle is identified (here shown with a red plane)

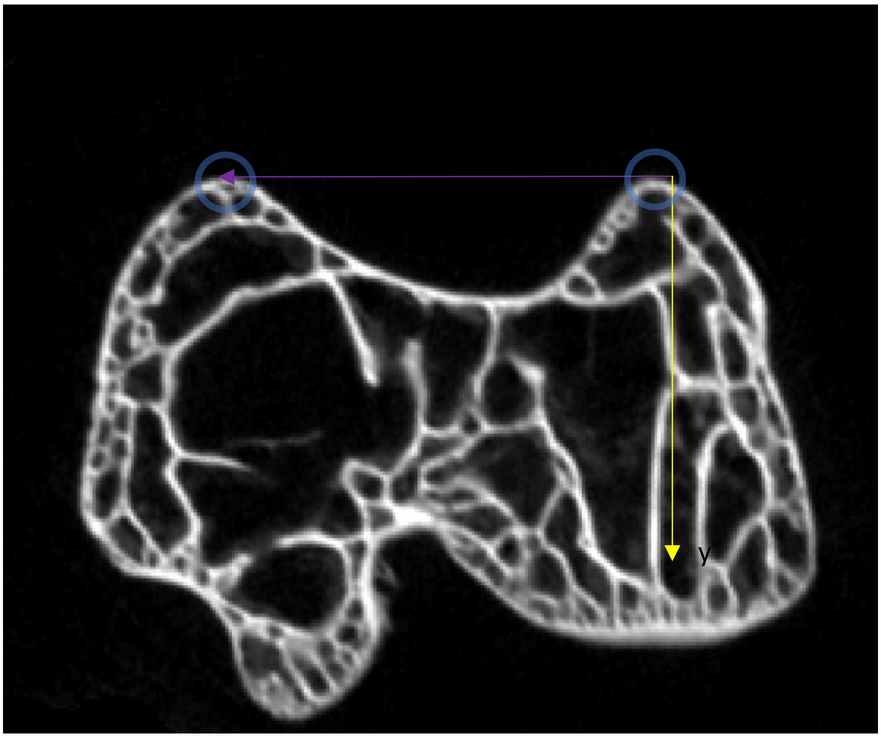


B. The cross-section corresponding to the medial condyle PD mid-level (identified by the red plane in the step A) is here visualized. The femur is rotated in the *xy* plane such that it has the *x*-axis contacting the two dorsalmost points of the distal epiphysis (points shown with blue circles, semi-lunar notch outlined in in orange).

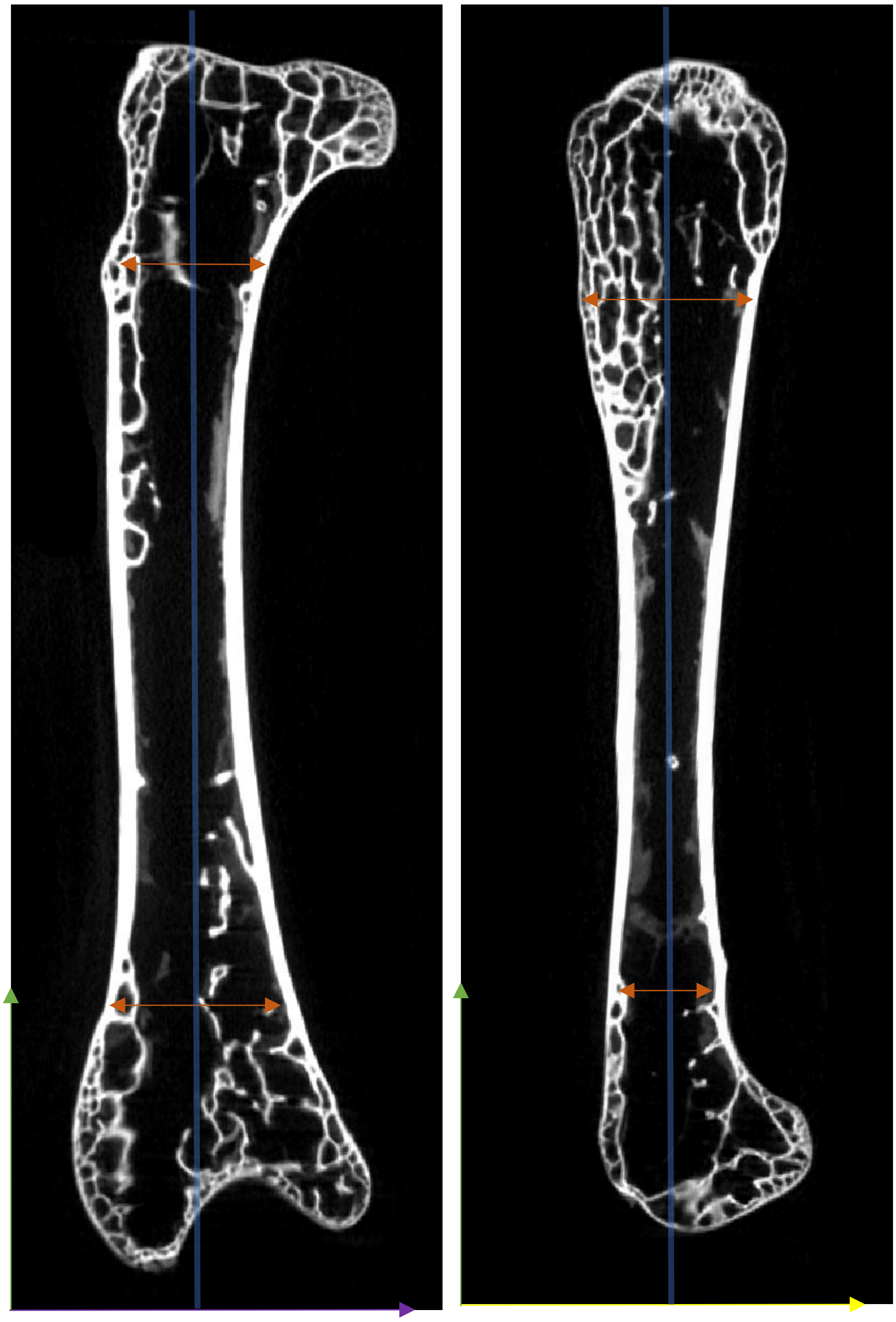

D. Femoral coronal (left) and sagittal (right) sections. Rotating the bone in the two views, the femur is oriented in the *xz* and *yz* planes, respectively, by aligning the mid-points of the proximal and distal metaphyseal regions (i.e. transitional regions between epiphysis and diaphysis; shown with an orange double-headed arrow) such that they both lie on a line (shown in shaded blue) which is parallel to the *z*-axis (shown in green)

### Section S3

The studied bones, once oriented as detailed Section S2 and imported in FIJI (Schindelin et al. 2012), were processed to isolate image sub-stacks representing the diaphysis, and the proximal and the distal epiphyses. Concerning the diaphysis, from stacks of the oriented bones we extracted a sub-stack including the central 35% of whole bone length. After a visual preliminary assessment, we considered this range as optimal to study diaphyseal structural variability, while excluding epiphyseal articular processes. As for the epiphyses, to analyse homologous regions we used standard anatomical markers, identified by moving proximodistally along the stack. The epiphyseal stacks were isolated for the humeri and ulnae, but not for the femora (for which no epiphyseal variables were analysed). As further detailed below, for the humerus, the distalmost boundary of the proximal epiphysis was defined as the beginning of the tricipital (or pneumotricipital) fossa, while the proximalmost boundary of the distal epiphysis was defined as the beginning of the dorsal condyle. For the ulna, the distalmost boundary of the proximal epiphysis was identified as the first level at which the 2D cross-sections of the bone approximate an annular shape; i.e. the end of the proximal epiphysis articulations. To define the proximalmost limit of the distal epiphysis, we identified the first 2D-cross sections that deviate from an approximately annular shape; i.e. the beginning of distal epiphyses articulations.

The steps followed to isolate proximal and distal epiphyses in a homologous way are here summarized on the humerus and the ulna of *Anser fabalis* NHMUK 1895-2-6-9

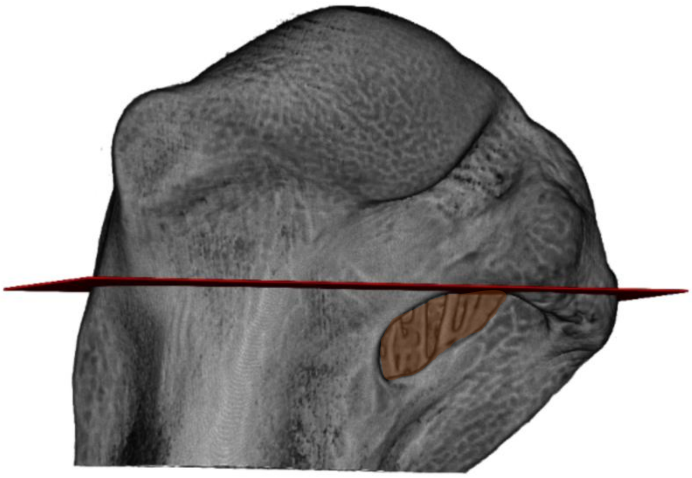

A. In this work, the humeral proximal epiphysis is considered the region proximal to proximalmost point of the tricipital (or pneumotricipital, in case of pneumatic taxa) fossa. The (pneumo)tricipital fossa is highlighted in brown, while the limit is highlighted with a red plane.

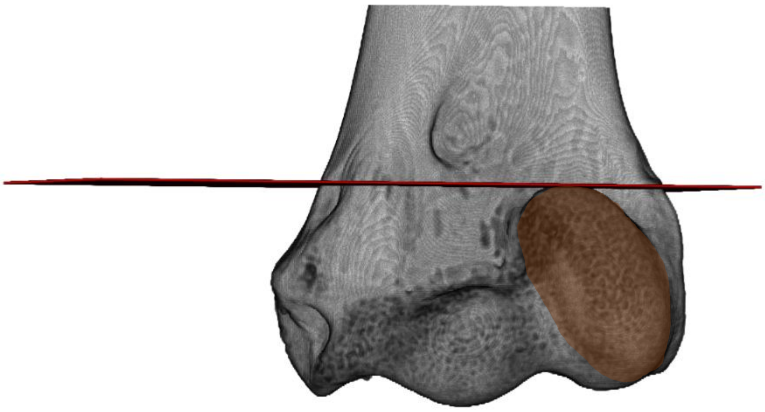

B. In this work, the humeral distal epiphysis is considered the region that is distal to the proximalmost point of the dorsal condyle. The dorsal condyle is highlighted in brown, while the limit is highlighted with a red plane.

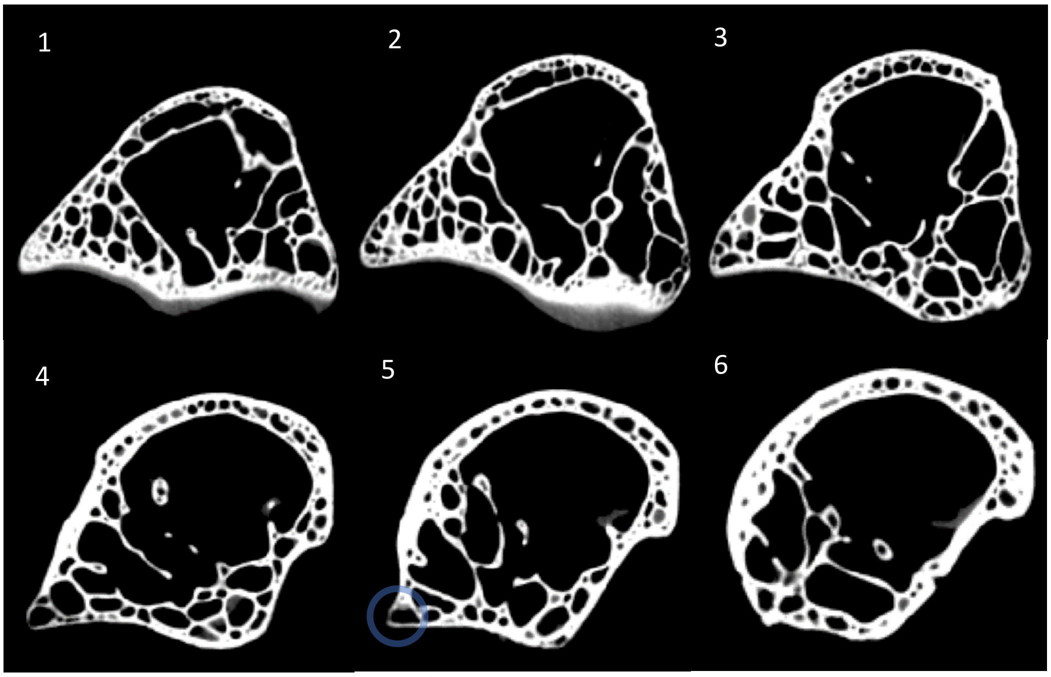


C. In this work, to homologously define the ulnar proximal epiphysis, we exploited the change of profile in cross-sections (deriving from moving proximodistally along the oriented bone) which is similar in all the studied birds. Specifically, figures 1-6 show a gradual movement from a section (1) cutting the proximal ulna across the semi-lunar notch (see comparison with Section S2, ulnar orienting, figure B, above), hence well within the proximal epiphysis, to other sections (2-5) which gradually show less features typical of the articular facets and instead tend to resemble more the typical annular shape of diaphyseal sections (e.g. figure 6). In this case, figure 5 represents the section defined as the distal limit of the proximal epiphysis (corresponding to the red plane, highlighted in figure 7), identified with the disappearing of last protuberance (i.e. the one highlighted with a blue circle, in figure 5) corresponding to

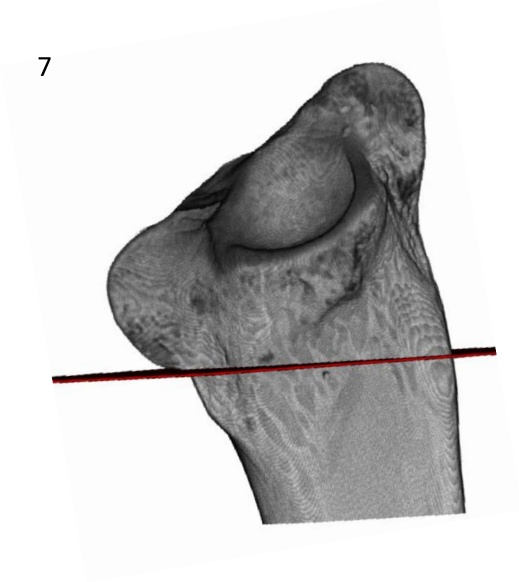

articular surfaces of the proximal epiphysis. This feature was used across all the sample to isolate the ulnar proximal epiphysis.

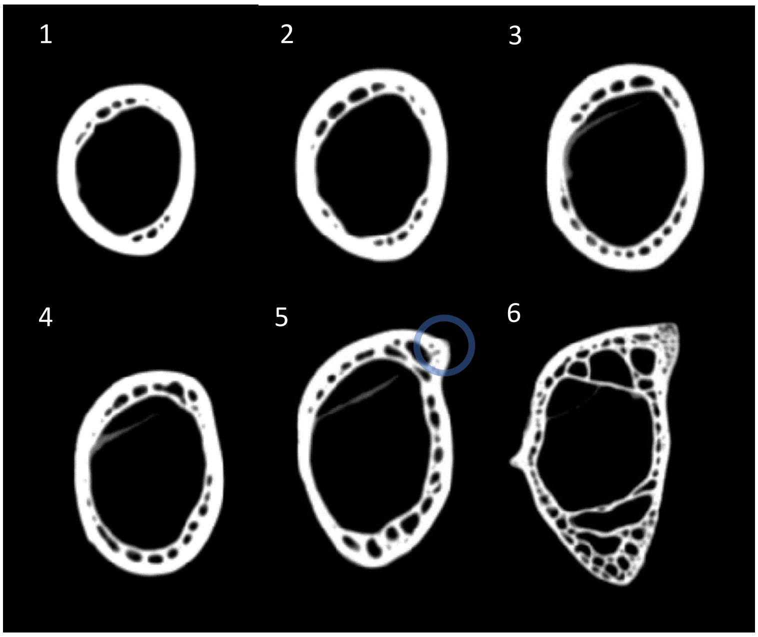


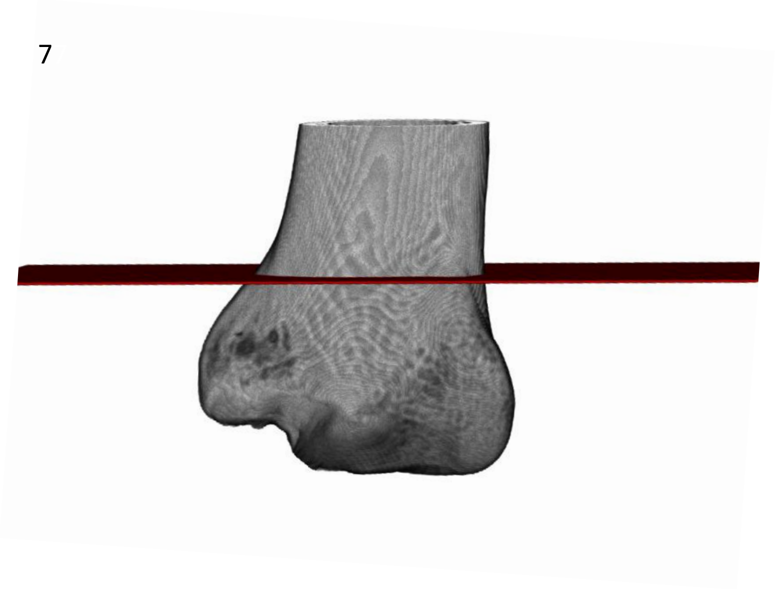


D. In this work, to homologously define the ulna distal epiphysis, we exploited the change of profile in cross-sections (deriving from moving proximodistally along the oriented bone) which is similar in all the studied birds. Specifically, figures 1-6 show a gradual movement from a section (1) cutting the ulna along the diaphysis (as one can see from the clearly annular shape of the section) to other sections (2-5) which gradually depart from the annular shape of figure 1 (e.g. they become increasingly elliptical). In this case, figure 5 represents the section defined as the proximal limit of the distal epiphysis (corresponding to the red plane, highlighted in figure 7), identified with the appearing of first protuberance (i.e. the one highlighted with a blue circle, in figure 5) corresponding to articular surfaces of the distal epiphysis. This feature was used across all the sample to isolate the ulnar distal epiphysis

**Section S4**

In FIJI, epiphyseal sub-stacks, isolated as detailed in Section S3, were pre-processed before extracting trabecular parameters (TP), the latter being potentially influenced by µCT resolution (Kivell et al. 2011; Lukova et al. 2024). As further explained and shown below, we used high-resolution scans, showing widely better quality of internal structure representation than e.g. medical CT scans. However, most of these µCT scans were not primarily acquired to quantify TPs, a condition not excluding sub-optimal resolution. To deal with it, we preliminarily and visually validated all the epiphyseal sub-stacks, ascertaining that individual trabeculae were sufficiently distinct from intertrabecular spaces and cortical bone. Furthermore, on all the epiphyseal stacks, to the exclusion of those deriving from scans acquired for this work (hence not affected by this issue), we performed an additional step, used by Bishop et al. to analyse dinosaur TPs extracted from scans with sub-optimal resolution (Bishop et al. 2018). Namely, we re-sampled the stacks by increasing the resolution (without affecting data structure), to enhance the performance of the binarizing (i.e. separating 'bone' from 'non-bone') algorithm that we used, i.e. local thresholding ('Auto Local Threshold' tool, 'Bernsen' algorithm (Bernsen 1986), radius=5, contrast threshold=25). Each result was visually validated, by comparing non-binarized and binarized stacks, and the latter were then 'cleaned', both automatically (purifying, i.e. removal of small, isolated particles, 'Purify' tool, BoneJ2 plugin (Domander et al. 2021)) and manually ('Paintbrush' tool) (see further details below).

To deal with the possible issue related to sub-optimal resolution for µCT scans collected from previous project, we followed some assessments and processing steps which are summarized as following, on the humeral proximal epiphysis of *Buteo rufofuscus* NHMUK 1862-1-18-1

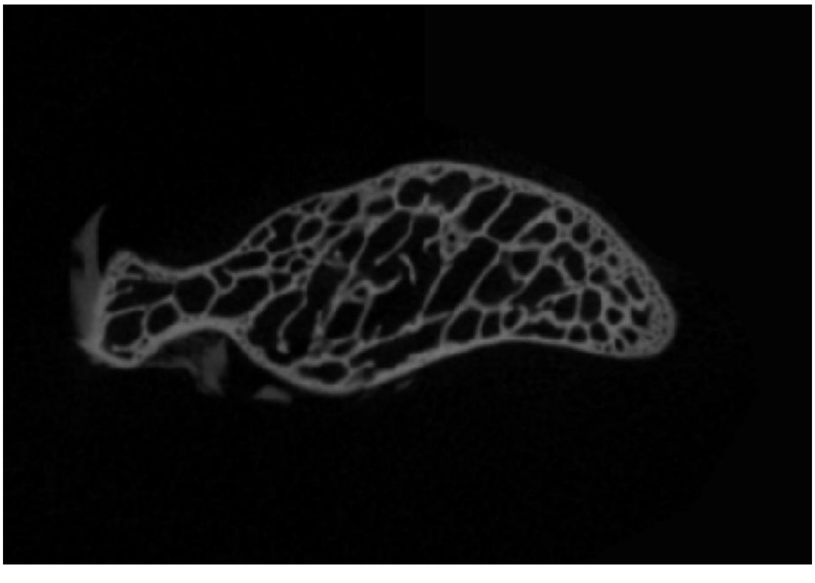

The epiphysis is assessed visually in its internal structure, by observing if individual trabeculae are clearly distinguished from surrounding cortical bone and intertrabecular spaces

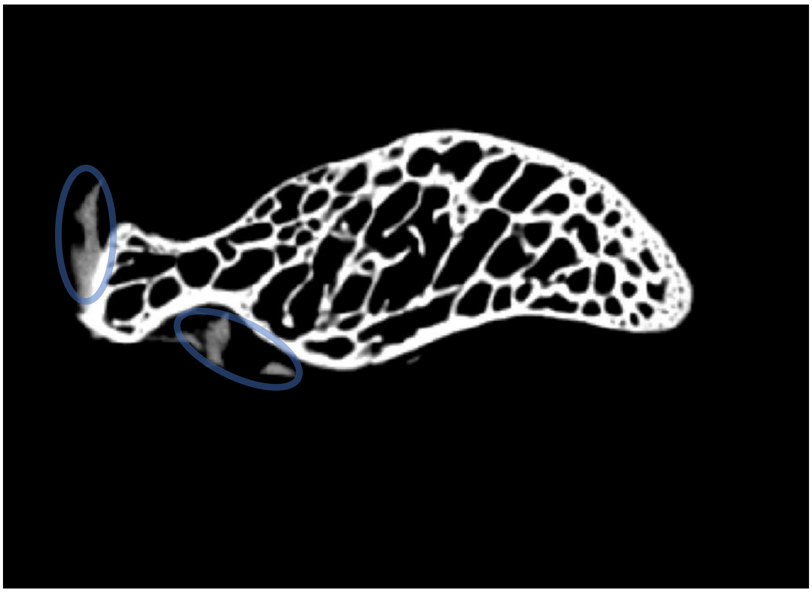

This assessment can be enhanced after tuning brightness and contrast (Adjust->Brightness and Contrast) in FIJI (Schindelin et al. 2012). It allows to distinguish what should be recognized as bone, clearly lighter and tending to be white, from other grey areas which should be deleted (see below), here highlighted with blue ellipses and representing scanning noise or soft tissues, as cartilages

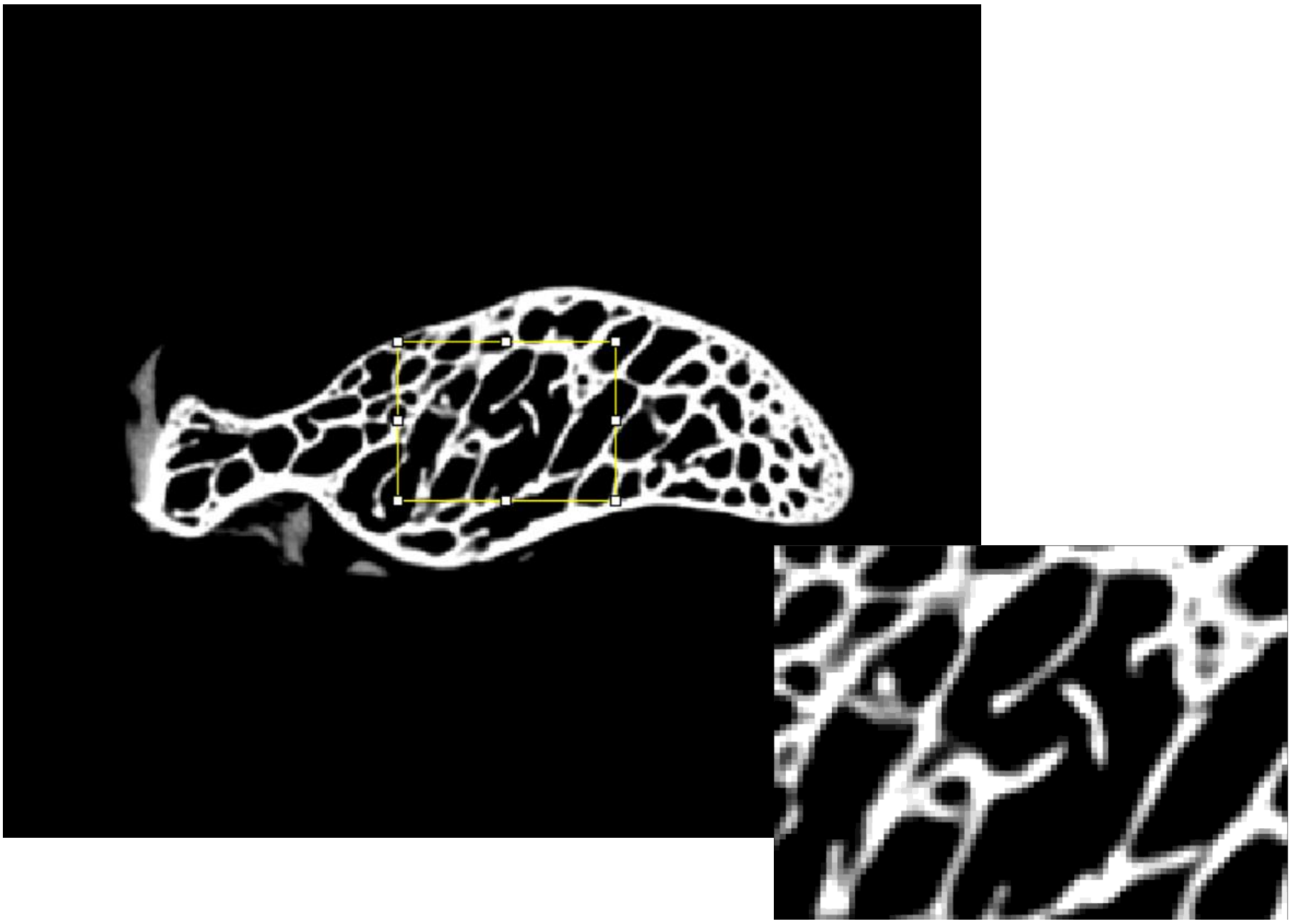

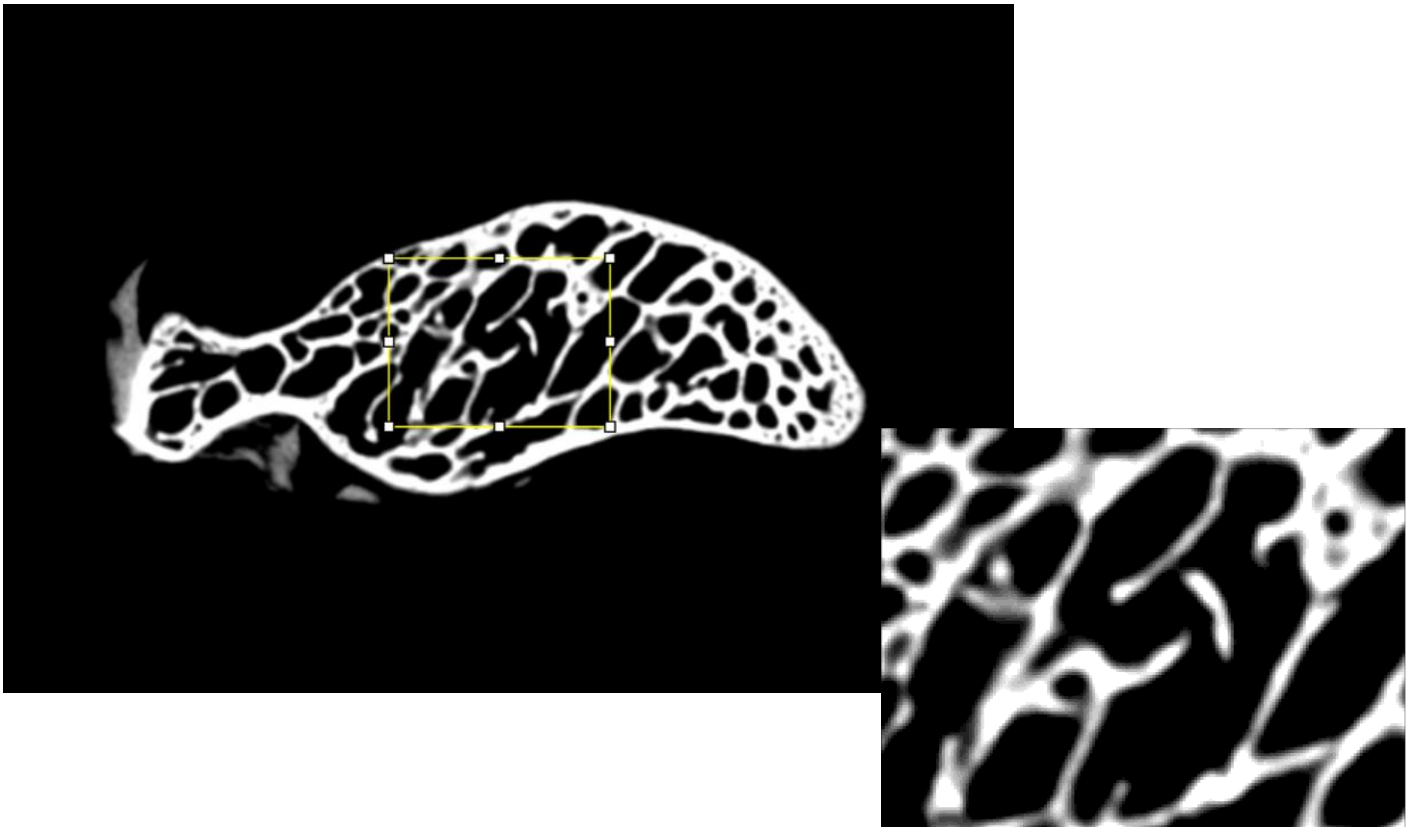

As suggested by Bishop et al. (2018), re-sampling the stacks to increase resolution (but without affecting the data structure) increases the number of voxels representing trabeculae and intertrabecular spaces and, ultimately, the local contrast, i.e. at the transition between ‘bone’ and ‘non-bone’. This can be observed confronting figure C and figure D above. In C, a rectangular sample of trabecular bone is extracted from the epiphysis of the stack before re-sampling and it is visualized in its magnified version on the bottom right. In D, the same epiphysis and sample are shown but after re-sampling to increase scan resolution. As one can observe, the transition area between trabeculae and intertrabecular spaces are better defined in D. Due to this feature, re-sampling is particularly efficient to study trabecular parameters from scans with potentially sub-optimal resolution, when it is combined with local thresholding segmentation algorithms, improving the performance of the latter (as done by Bishop et al. 2018). For scans potentially represented

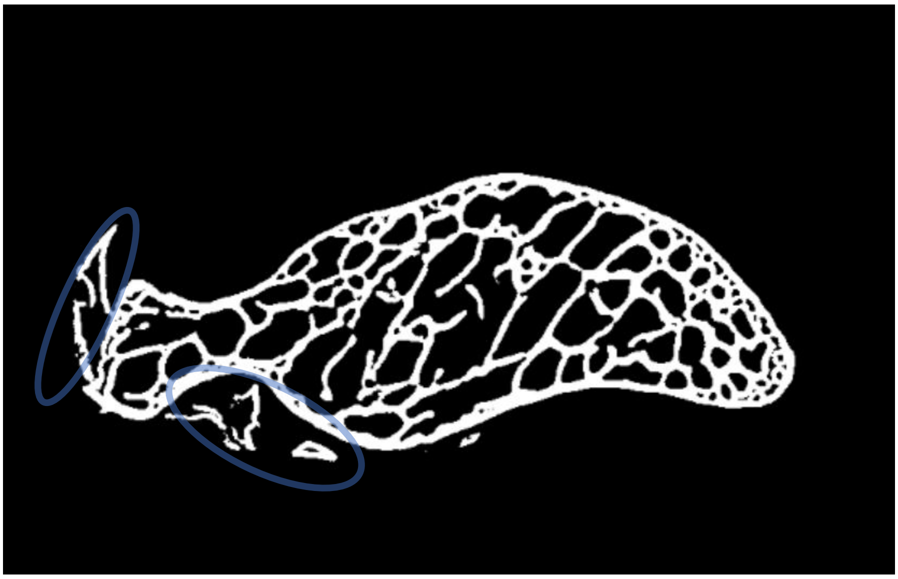

by sub-optimal resolution, we decided to follow this approach, re-sampling the epiphyseal stacks, and specifically doubling the 3D resolution (through the 'Scale' FIJI tool, and using a bicubic interpolation). Differently from Bishop et al. (2018), who chose a scaling factor of 3 (i.e. tripling the resolution), we considered a factor of 2 sufficient. It is justified by the fact that our resolution issues are not severe as those anticipated by Bishop et al. (who applied this protocol to medical CT-scans) and by the necessity to avoid the generation of too computationally demanding epiphyseal stacks as input for the automatic separation of trabecular bone from cortical bone in R (one of subsequent steps; see main text, Veneziano et al. 2021; and Alfieri et al. 2025). Thus, after re-sampling using a scaling factor=2, we segmented 'bone' from 'non-bone' (i.e. binarizing) in the epiphyseal stacks through local thresholding ('Auto Local Threshold' tool) using the 'Bernsen' algorithm (Bernsen 1986). The binarization result is shown in figure E. Through preliminary tests we found the likely optimal algorithm settings across our sample, i.e. window radius=5; contrast threshold=25, that noticeably align to those used by Bishop et al., with the contrast threshold lying in a range making the erroneous recognition as bone of high-density non-bone material less likely (Bishop et al. 2018). Each segmentation result was visually validated by comparing non-segmented (e.g. figure B) and segmented (e.g. figure E) stacks. It allowed us to establish how the binarized stacks needed to be 'cleaned'. The 'cleaning' procedure was done automatically ('Purify' tool, BoneJ2 plugin, Domander et al. 2021), deleting small particles (i.e. few voxels) assigned to the bone phase but being 'floating' (i.e. not connected) in 3D (due to nature of trabecular bone, unconnected particles necessarily represent imaging artifacts). Besides automatic cleaning, exploiting the comparison between non-binarized and binarized stacks, we could additionally perform manual cleaning (e.g. remaining noise around the bone from soft issues, e.g. cartilages). In this specific case, one can see that after binarization some non-bone grey values in the original non-binarized stack were erroneously recognized as bone (see blue ellipses in figure E). Hence, these regions, which should not

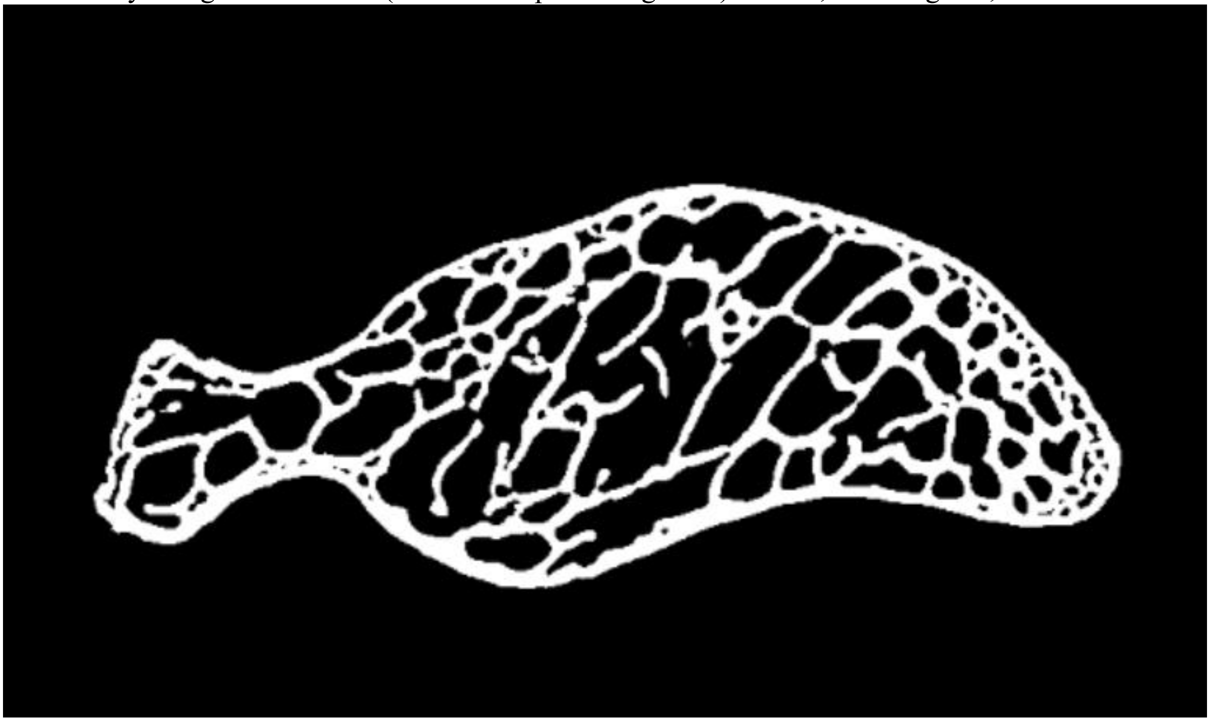

be part of the bone (‘foreground’, white) phase, were manually assigned to ‘non-bone’ (foreground, black) through the FIJI ‘Paintbrush’ tool. It results in the correctly binarized stack visualized in figure E.

**Section S5**

In R, trabecular bone was isolated from cortical bone through the ‘makeBrush’ function (‘EBimage’ package, Pau et al. 2010) and the ‘splitBone’ function (‘indianaBones’ package, Veneziano et al. 2021). We quantified trabecular properties from stacks including only trabecular bone (‘Regions of Interest’, ROIs), through both traditionally employed structural and recently developed topological variables. For the former, we imported ROIs in FIJI and used the respective BoneJ2 routines to compute: ‘degree of anisotropy’ (DA, unitless), ‘trabecular thickness’ (Tb.Th, mm), ‘bone volume fraction’ (BV/TV, unitless) and ‘connectivity density’ (Conn.D, $mm^{-3}$). To compute topological properties, ROIs were first imported in Avizo 2020.2 and transformed into topological skeletons (‘Auto Skeleton’ tool, where trabecular networks were reduced to lattices compounded by nodes (trabecular connections) and branches (trabeculae), which retains the topological properties of the structure (Veneziano et al. 2021; and Alfieri et al. 2025). Topological skeletons were imported in R and the packages ‘indianaBones’ (Veneziano et al. 2021) and ‘Rdimtools’ (You and Shung 2022) were used to extract ‘node density’ (NodDen), ‘trabecular tortuosity’ (TrabTort), ‘average trabecular length’ (TrabLen), and ‘fractal dimension’ (FD). For NodDen and TrabTort, which are represented by the range of values across the topological skeleton, the arithmetic mean and median were used to calculate NodDenAver, NodDenMed, TrabTortAver and TrabTortMed.

From the proximal/distal humeral and ulnar trabecular datasets, we discarded data obtained from ROIs including a too low number of trabeculae. Indeed, this condition would make the TP result meaningless. A lower limit of 50 trabeculae is often used in mammal studies (e.g., see Amson et al. 2017; Mielke et al. 2018; Alfieri et al. 2022, 2023), with the number of trabeculae approximated by Connectivity (a parameter preliminarily computed to extract Conn.D). We generally followed this recommendation but lowered it to 45 to not exclude some observations with Connectivity being only slightly lower than 50 (e.g., 49.5 in *Archilochus colubris* uf-o-26675 proximal humerus or 48 in *Piprites chloris* FMNH 290398, distal humerus).

**Section S6**

Along the diaphysis, we computed cross-sectional properties (CSP) through a slice-by-slice approach (Amson 2019; Alfieri et al. 2022, 2023) implemented in FIJI. Specifically, each diaphyseal slice was thresholded (‘Threshold’ tool), purified and used to compute ‘global compactness’ (Cg, %), ‘total cross-sectional area’ (TotAr, $mm^2$), (both through the ‘Measure’ tool) and, through the ‘SliceGeometry’ tool of BoneJ (Doube et al. 2010), we computed ‘cross-sectional area’ (CSA, $mm^2$), ‘seconds moments of area’ (around the minor, Imax, and major axes, Imin; $mm^4$), ‘polar section modulus' (Zpol, $mm^3$), 'cross-sectional shape' (Imax/Imin, unitless) and 'perimeter' (mm). The latter was not used in the statistical analysis but computed to estimate the body mass proxy (see main text). As it often occurs when the slice-by-slice protocol is employed (Amson 2019; Alfieri et al. 2022, 2023), during CSP computation, some slices arose as yielding biased data (e.g. due to bone damages, interrupting the cortical bone outline). We excluded data from these slices without data imputation, since we were not interested in the diaphyseal profile of CSP variation (differently from e.g. Amson 2019) and we never found substantially long flawed segments. CSP from all the diaphyseal slices served to compute mean and mid-diaphyseal values (the latter from the 50% of bone length, often considered the most informative diaphyseal level in tetrapods, Laurin 2004; de Margerie et al. 2005; Habib and Ruff 2008)). The isolated flawed diaphyseal segments mentioned above, never intersected the mid-diaphyseal level. To compute F/H_SS we quantified femoral $Zpol_{50}$ and length, in addition to humeral length. Bone lengths were measured multiplying the slice number comprised between the proximalmost and distalmost point of the oriented bone by the scan resolution. Femoral $Zpol_{50}$ and humeral/ulnar $Ct.Th_{50}$ were quantified through 'SliceGeometry' on the thresholded and purified mid-diaphyseal 2D slices.

### Section S7:

Additional literature used to score flight style categories for the studied taxa is Rand (1936) (for *Atelornis*), Tobalske and Dial (1994) (for parrots), Kemp and Kemp (1980) (for *Bucorvus*), Witherby et al. (1938) (for *Coracias*), Stiles and Whitney (1983) (for *Oxyruncus*), Tobalske (2001) for woodpeckers, Keith et al. (1970) for flufftails, Skaed 1949 (for *Upupa epops*)

**Figure S1.** The gap statistic metric is maximized in correspondence to 9 clusters, hence allowing us to determine that **9** is the optimal number of eco-clusters is the best performing scheme to classify the ecological diversity included in the 140 x 15 matrix (built as detailed in the main text)

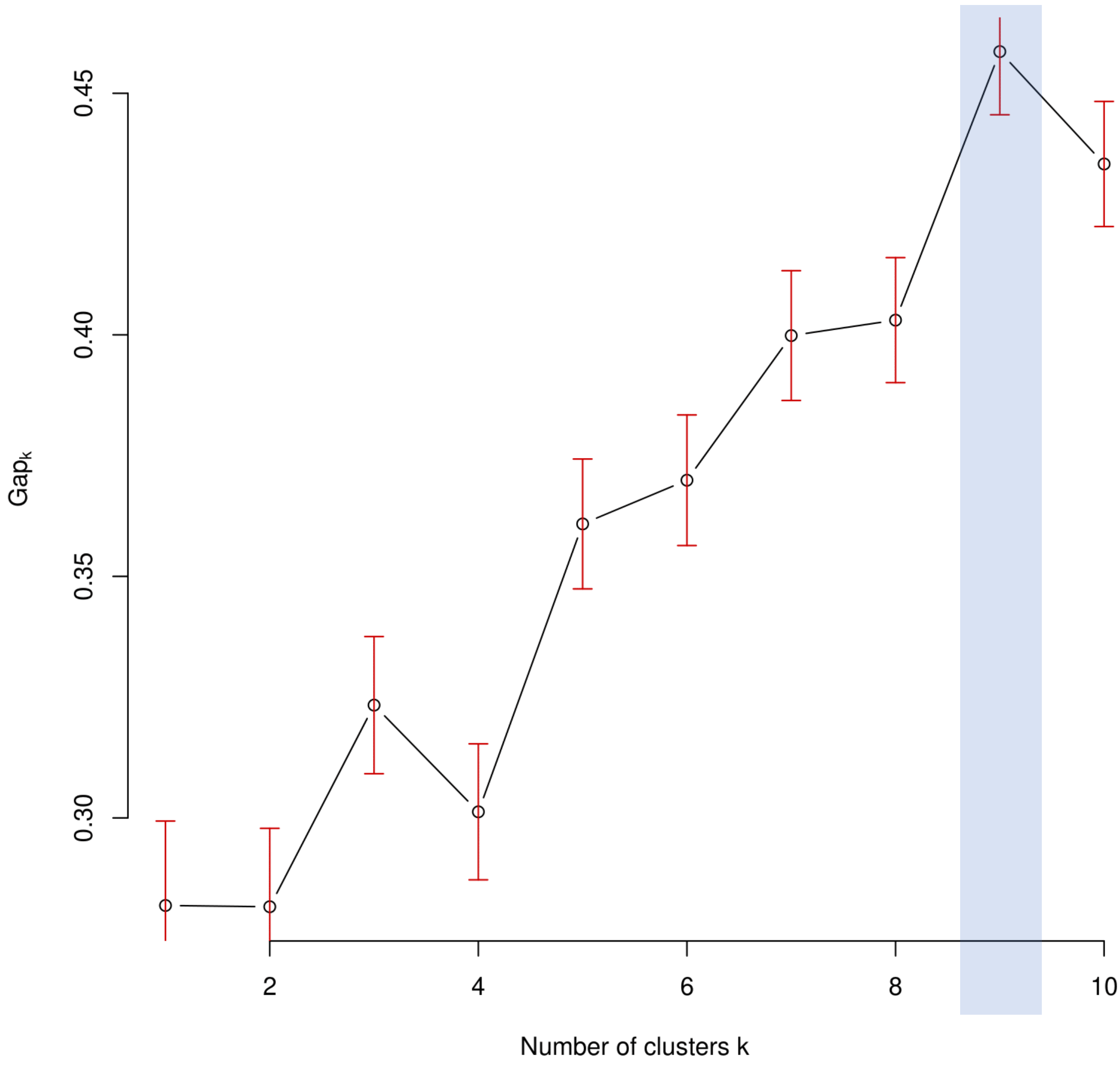

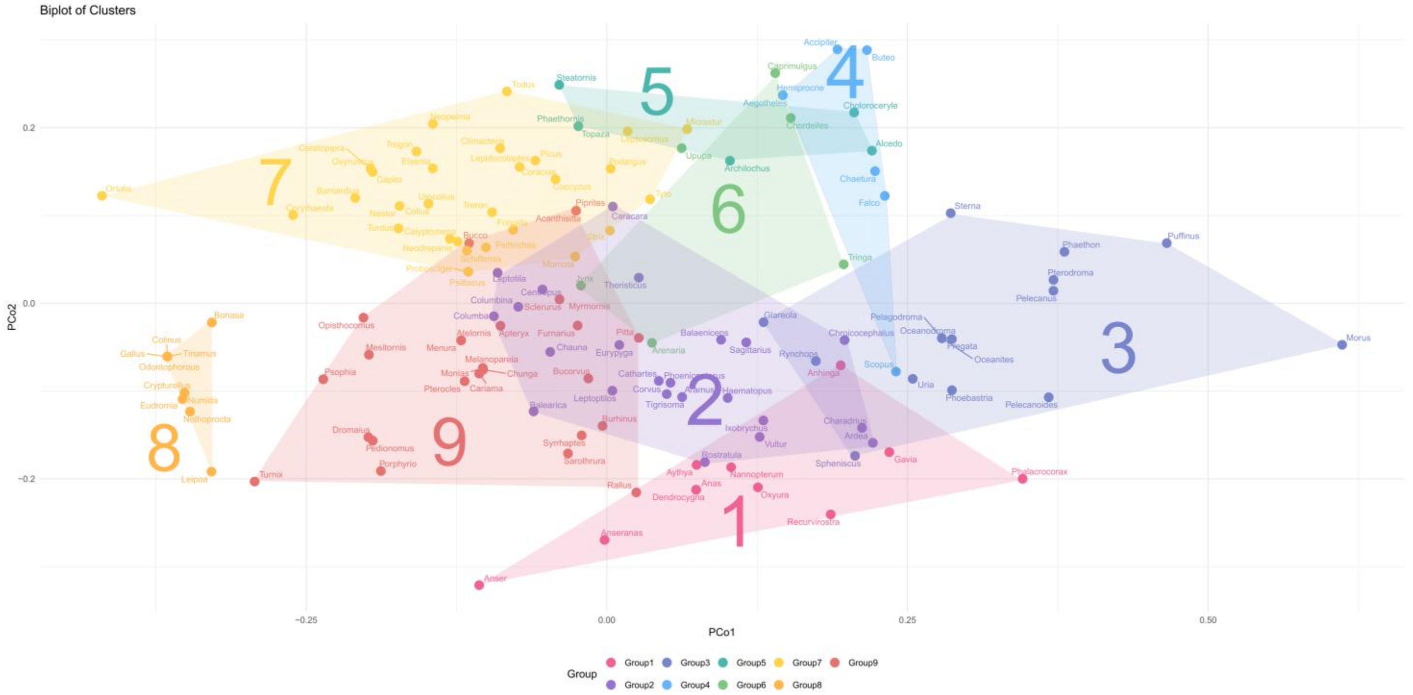


**Figure S2.** Biplot built with PCo1 and PCo2 from the PCoA run on the dissimilarity matrix, in turn deriving from the ecological dataset, showing the distribution of the 140 taxa studied in this work. The taxa are grouped according to their belonging to eco-clusters (as shown through convex hulls) which are 9, since it is the optimal number deriving from gap statistics (see Fig. S1). The belonging of each taxon to eco-clusters 1-9 is here exclusively based on hierarchical clustering.

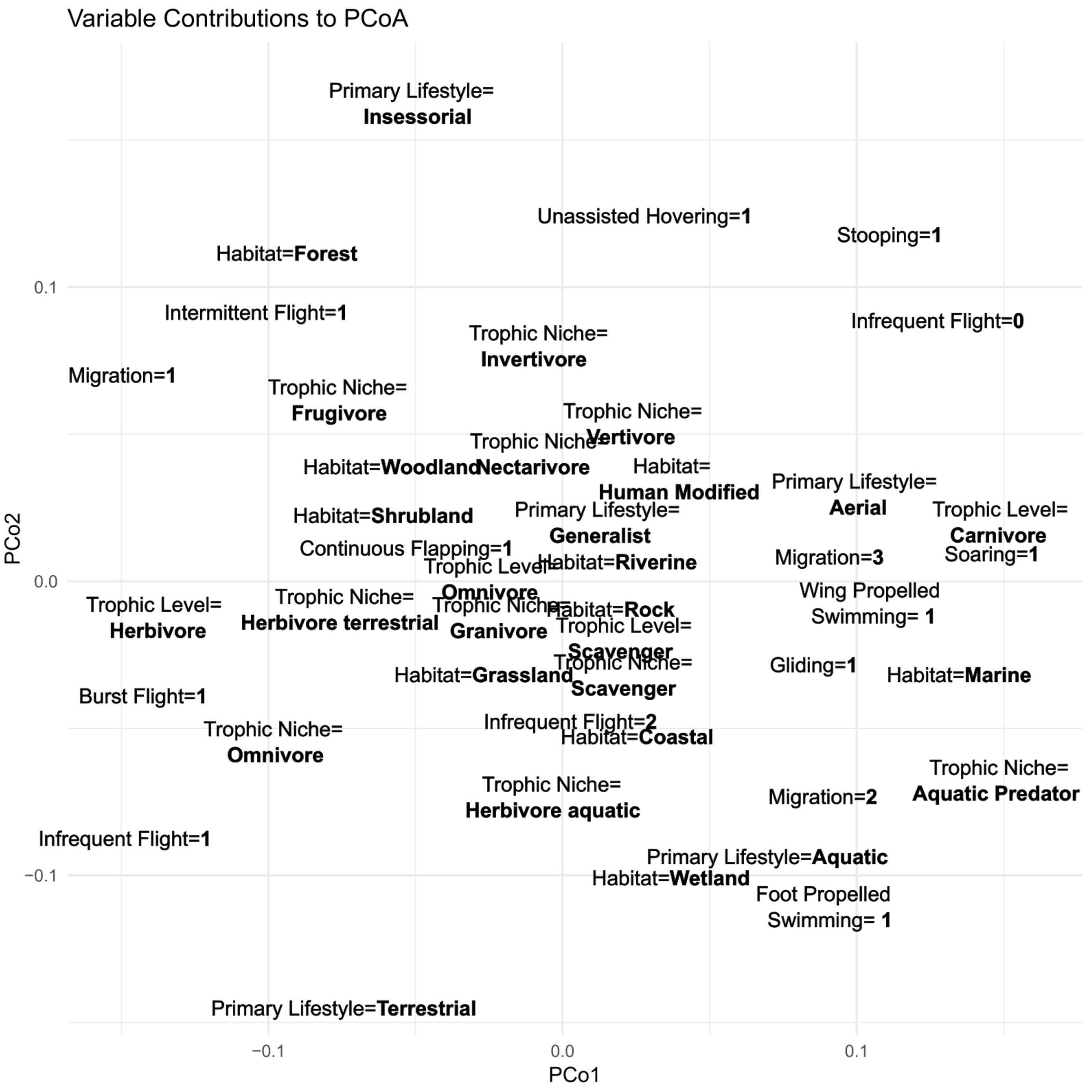


**Figure S3.** Plot showing how the levels of the discrete variables (compounding the 140 x 15 ecological dataset) contribute to the taxa distribution shown on the PCo1-PCo2 biplot (Fig. S2 and Fig. S4).

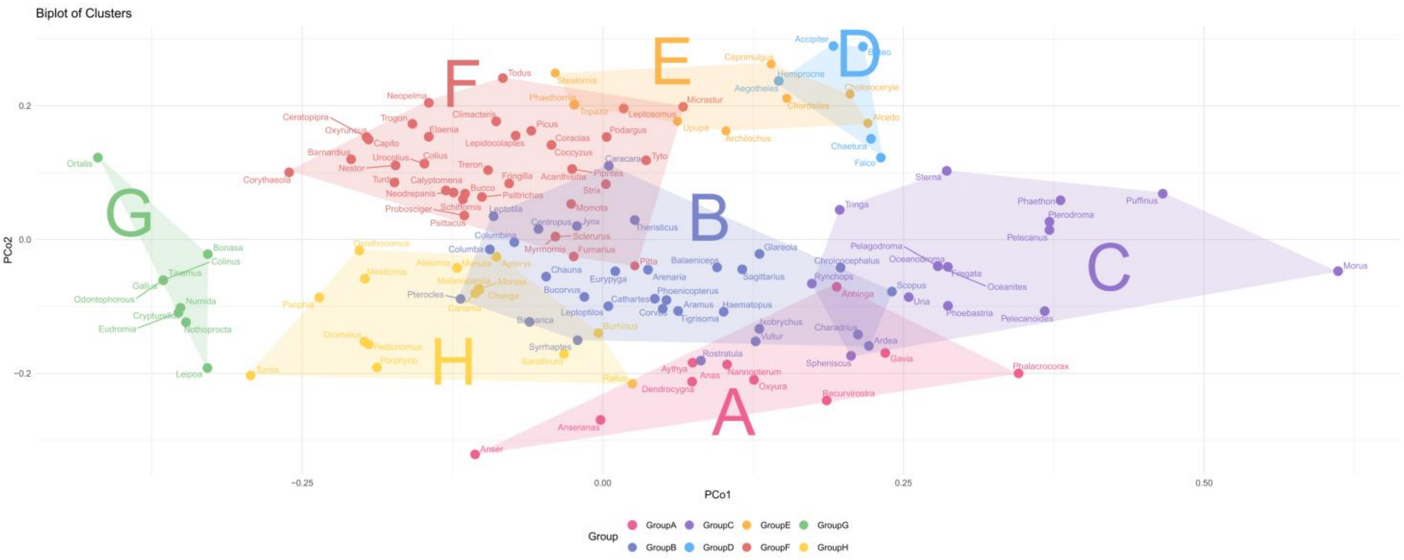


**Figure S4.** Biplot built with PCo1 and PCo2 from the PCoA run on the dissimilarity matrix, in turn deriving from the ecological dataset, showing the distribution of the 140 taxa studied in this work. The taxa are grouped according to their belonging to eco-clusters (as shown through convex hulls) that are 8 (we edited some of the assignments deriving from hierarchical clustering, as detailed in Section S8, see below) and that, to avoid confusion with the eco-clusters in Fig. S2 (only defined by hierarchical clustering, see above), are here termed with letters, i.e. A-H.

**Section S8:**

EcoClusters 1-9 are those defined by hierarchical clustering (and which number, i.e. 9, derives from the gap statistics) (see Figs. S1-S2, above). Their ecological features were identified through PCos-ecological variables correlation (Fig. S3). It allowed us to assess whether all taxa assignments to eco-clusters 1-9 were correct. As detailed below, in cases of misassignment, we manually edited some eco-clusters. This resulted in a final eco-cluster system based on both hierarchical clustering and our assessments, comprising eight groups, Eco-Clusters A-H.

Eco-Clusters 1-9 (only defined by hierarchical clustering, see Fig. S2)

EcoCluster 1. Members identified by hierarchical clustering are:

- *Anseranas*
- *Anser*
- *Anas*
- *Aythya*
- *Oxyura*
- *Dendrocygna*
- *Recurvirostra*
- *Gavia*
- *Anhinga*
- *Phalacrocorax*
- *Nannopterum*

They are characterized by:

- **Foot propelled swimming=1** (all the taxa)
- **Habitat=Wetland** (all the taxa but *Anser,* showing **Habitat=Coastal**, and *Nannopterum*, showing **Habitat=Marine**)
- **Primary Lifestyle=Aquatic** (72% of the taxa, with the exclusion of *Anseranas*, *Anser* and *Recurvirostra*, which show **Primary Lifestyle=Terrestrial)**

Comments:

- Three taxa do not show **Primary Lifestyle=Aquatic**. They are *Anseranas*, *Anser* and *Recurvirostra*, which show, instead, **Primary Lifestyle=Terrestrial.** The latter feature is one of the defining features of EcoCluster 2 (see below) but since these three taxa show **Foot propelled swimming=1**, which is totally absent from EcoCluster 2, we preferred to have them in EcoCluster 1

------------------------------------------------------------------------------------------------------------

EcoCluster 2, Members identified by hierarchical clustering are:

- *Chauna*
- *Phoenicopterus*
- *Columbina*
- *Columba*
- *Leptotila*
- *Centropus*
- *Aramus*
- *Balearica*
- *Chroicocephalus*
- *Rostratula*
- *Charadrius*
- *Haematopus*

- *Eurypyga*
- *Leptoptilos*
- *Theristicus*
- *Ardea*
- *Ixobrychus*
- *Tigrisoma*
- *Balaeniceps*
- *Cathartes*
- *Vultur*
- *Sagittarius*
- *Caracara*
- *Corvus*

They are characterized by:

- No exclusive ecological traits
- Since it is a numerous EcoCluster (i.e. 24 taxa) and it occupies the central region of Fig. S2, it is here considered the EcoCluster of ecologically non-differentiated birds

Comments:

- **Primary Lifestyle = Terrestrial** is very common (all the taxa but *Chroicocephalus*), but it is shared with other EcoClusters (e.g. EcoCluster8, EcoCluster9), hence it is not exclusive
- **Continuous Flapping=1** in the 75% of the taxa, but it is not exclusive (e.g. 82 % in EcoCluster 1, 100% in EcoCluster 5)
- **Gliding=1** in all the taxa but for this trait they resemble EcoCluster3 (which has **Gliding=1** for all the taxa but one taxon), Eco-Cluster4 (all the taxa). Thus, **Gliding=1** is not exclusive
- They have a quite common **Soaring=1** (62.5%) but it is shared with EcoCluster 3 (which has 69% of taxa with **Soaring=1**) and EcoCluster4 (all the taxa), hence it is not exclusive

------------------------------------------------------------------------------------------------------------

EcoCluster3, Members identified by hierarchical clustering are:

- *Uria*
- *Sterna*
- *Rynchops*
- *Glareola*
- *Phaethon*
- *Spheniscus*
- *Phoebastria*
- *Oceanites*
- *Pelagodroma*
- *Oceanodroma*
- *Puffinus*
- *Pelecanoides*
- *Pterodroma*
- *Pelecanus*
- *Fregata*
- *Morus*

They are characterized by:

- **Habitat=Marine** (all the taxa but *Glareola*)
- **Trophic Niche= Aquatic predator** (all the taxa but *Glareola*)

- **Wing Propelled Swimming=1** (although not present in most of the taxa, Fig. S3 and the position of **Wing Propelled Swimming=1** clearly suggest that this trait is distinctive and strongly contributes to set apart this EcoCluster)

Comments:

- They have **Gliding=1** in all the taxa but one taxon. However, for this they resemble EcoCluster2 (which have **Gliding=1** in all the taxa). Thus, it is not exclusive
- They have a quite common **Soaring=1** (ca. 69%) but it is shared with EcoCluster 2 (which has 62.5% of taxa with **Soaring=1**). Thus, it is not exclusive
- Two taxa show **Foot Propelled Swimming=1** (*Pelecanus* and *Morus*) but they do not have the other characteristics of EcoCluster 1(i.e. **Habitat=Wetland**, **Primary Lifestyle=Aquatic**). Thus, we preferred to leave them in EcoCluster3
- The pratincole *Glareola*, assigned to this EcoCluster by hierarchical clustering, does not show **Habitat=Marine, Trophic Niche= Aquatic predator** and not even **Wing Propelled Swimming=1.** Since it does not show any distinctive ecological trait (allowing it to be assigned to other EcoClusters), we assign it to EcoCluster2 (ecologically non-differentiated birds).

------------------------------------------------------------------------------------------------------------

EcoCluster 4, Members identified by hierarchical clustering are:

- *Chaetura*
- *Hemiprocne*
- *Aegotheles*
- *Scopus*
- *Buteo*
- *Accipiter*
- *Falco*

They are characterized by:

- **Stooping=1** (all the taxa), although it is not exclusively shown by EcoCluster 4 (i.e. shown by all representative of EcoCluster5, see below).

Comments:

- **Gliding=1** in all the taxa but for this trait they resemble EcoCluster3 (which have **Gliding=1** for all the taxa but one taxon), Eco-Cluster2 (all the taxa). Thus, **Gliding=1** is not exclusive
- All of them show **Soaring=1,** but it is not exclusive (e.g. 62.5% of taxa of EcoCluster2, or 69% in EcoCluster3)
- **Stooping=1** is very common in EcoCluster5 too (see below) but the variable loading plot (Fig. S3, see **Stooping=1**'s position in the plot) suggests that this behavior was especially responsible to set apart EcoCluster4. Also, EcoCluster5 is already characterized by a distinctive behavior (i.e. **Unassisted Hovering=1**, see below). Thus, we can consider **Stooping=1** as the distinctive behavior for eco-cluster 4.
- **Infrequent Flight=0** in all the taxa and position in Fig. S3 suggests that it contributed to set apart EcoCluster4. However, this condition is shared with many EcoClusters (e.g. is present in all the taxa in EcoClusters 2, 3; and in most of the taxa in EcoCluster1) and not in only another EcoCluster (as Stooping=1, see above). Hence, it is not exclusive.
- The hamerkop *Scopus* is assigned to this EcoCluster (and indeed it shows **Stooping=1**). However, its position in Fig. S2, widely outlying the other members of EcoCluster4, led us to assign it to other EcoClusters. Its **Habitat=Wetland** is likely responsible of its position approaching EcoCluster1. However, *Scopus* does not show the other defining features of EcoCluster1, i.e. **Foot propelled swimming=1** and **Primary Lifestyle=Aquatic**. Hence, we preferred to assign *Scopus* to EcoCluster2 (ecologically non-differentiated birds).

------------------------------------------------------------------------------------------------------------

EcoCluster 5, Members identified by hierarchical clustering are:

- *Topaza*
- *Phaethornis*
- *Archilochus*
- *Steatornis*
- *Alcedo*
- *Choloroceryle*

They are characterized by:

- **Unassisted hovering=1** (all the taxa)

Comments:

- All of them show **Stooping=1.** For this, they resemble EcoCluster4 (see above). However, we considered **Stooping=1** as a defining trait for EcoCluster4 (as justified above)

------------------------------------------------------------------------------------------------------------

EcoCluster 6, Members identified by hierarchical clustering are:

- *Tringa*
- *Arenaria*
- *Chordeiles*
- *Caprimulgus*
- *Upupa*
- *Jynx*

They are characterized by:

- No exclusive ecological traits
- Differently from EcoCluster4, EcoCluster6 does not allow us to identify an ecological trait mainly responsible for it to be set apart in Fig. S2 (i.e. it strongly overlaps with other EcoClusters). This, besides its small size (i.e. only 6 taxa) led us to split EcoCluster6 members across other EcoClusters (as justified in the comments below)

Comments:

- We recognized that three of the taxa of EcoCluster6 (i.e. *Chordeiles*, *Caprimulgus* and *Upupa*) show **Unassisted Hovering=1,** defining trait of EcoCluster5. Hence, it allows us to move these three taxa in EcoCluster5.
- The other three taxa, i.e. *Tringa*, *Arenaria* and *Jynx*, are moved to other EcoClusters. *Tringa*, showing **Wing Propelled Swimming=1** and **Trophic Niche= Aquatic predator**, is moved to EcoCluster3. *Arenaria* and *Jynx*, not showing any trait suggesting their possible belonging to any of the other EcoClusters, are assigned to EcoCluster2 (ecologically non-differentiated birds; see also *Arenaria* and *Jynx*'s positions on Fig. S2, clearly suggesting their belonging to EcoCluster2).

------------------------------------------------------------------------------------------------------------

EcoCluster 7, Members identified by hierarchical clustering are:

- *Ortalis*
- *Treron*
- *Corythaeola*
- *Coccyzus*
- *Podargus*
- *Tyto*
- *Strix*
- *Colius*

- *Urocolius*
- *Leptosomus*
- *Trogon*
- *Coracias*
- *Todus*
- *Momota*
- *Picus*
- *Capito*
- *Micrastur*
- *Barnardius*
- *Psittrichas*
- *Psittacus*
- *Probosciger*
- *Nestor*
- *Neodrepanis*
- *Calyptomena*
- *Lepidocolaptes*
- *Ceratopipra*
- *Neopelma*
- *Schiffornis*
- *Oxyruncus*
- *Elaenia*
- *Climacteris*
- *Turdus*
- *Fringilla*

They are characterized by:

- **Primary Lifestyle=Insessorial** (94% of the taxa), **Habitat=Forest** (72% of the taxa), **Intermittent Flight=1** (66% of the taxa)

Comments:

- The galliform Chachalaca**,** *Ortalis*, is assigned to this EcoCluster. However, *Ortalis* is a burst flier (**Burst Flight =1**) and shows **Trophic Level=Herbivore**, as all the other galliforms that we studied, and this trait defines EcoCluster8 (see below). Hence, we preferred to assign *Ortalis* to EcoCluster8, grouping it with other burst fliers. See also in Fig. S2 the position of *Ortalis*, being intermediate between EcoCluster7 and 8.

-----------------------------------------------------------------------------------------------------------

EcoCluster 8, Members identified by hierarchical clustering are:

- *Eudromia*
- *Nothoprocta*
- *Tinamus*
- *Crypturellus*
- *Leipoa*
- *Numida*
- *Odontophorous*
- *Colinus*
- *Gallus*
- *Bonasa*

They are characterized by:

- **Burst Flight=1** (all the taxa), **Trophic Level=Herbivore** (all the taxa)

-------------------------------------------------------------------------------------------------------------

EcoCluster 9, Members identified by hierarchical clustering are:

- *Dromaius*
- *Apteryx*
- *Mesitornis*
- *Monias*
- *Pterocles*
- *Syrrhaptes*
- *Opisthocomus*
- *Sarothrura*
- *Porphyrio*
- *Rallus*
- *Psophia*
- *Turnix*
- *Pedionomus*
- *Burhinus*
- *Bucorvus*
- *Atelornis*
- *Bucco*
- *Cariama*
- *Chunga*
- *Acanthisitta*
- *Pitta*
- *Myrmornis*
- *Melanopareia*
- *Sclerurus*
- *Furnarius*
- *Piprites*
- *Menura*

They are characterized by:

- **Infrequent Flight=1** or **Infrequent Flight=2**, i.e. they are either reluctant flying or flightless, but without distinctive ecological traits which are shown by other reluctant flying/flightless taxa (e.g. Wing Propelled Swimming, Burst Flight), assigned to other EcoClusters

Comments:

- Nine taxa, i.e. *Pterocles, Syrrhaptes, Bucorvus, Acanthisitta, Pitta, Myrmornis, Sclerurus, Furnarius* and *Piprites,* are assigned to this EcoCluster but they do not show **Infrequent Flight=1** or **Infrequent Flight=2**. *Pterocles, Syrrhaptes* and *Bucorvus* do not show any distinctive trait of the other EcoClusters, and thus are assigned to EcoCluster2 (ecologically non-differentiated birds). *Acanthisitta, Pitta, Myrmornis, Sclerurus, Furnarius* and *Piprites* show **Habitat=Forest** (five of them), a feature that in two taxa (i.e. *Acanthisitta* and *Piprites*) is combined with **Primary Lifestyle=Insessorial.** These features remind more of the characteristics of EcoCluster=7, hence these taxa are moved there. The same has been done for *Bucco* that, while showing **Infrequent Flight=1,** it shows **Habitat=Forest** and **Primary Lifestyle=Insessorial**

After the adjustments justified as detailed above, we have the following **8 final EcoClusters** (that we will identify using letters instead of numbers, to avoid confusion with the EcoClusters originally identified only through cluster analysis [see above])

**Eco-Clusters A-H** (defined by hierarchical clustering results, edited by us, as justified above)

**EcoClusterA** (Foot propelled swimming=1, Habitat=Wetland, Primary Lifestyle=Aquatic)

- *Anseranas*
- *Anser*
- *Anas*
- *Aythya*
- *Oxyura*
- *Dendrocygna*
- *Recurvirostra*
- *Gavia*
- *Anhinga*
- *Phalacrocorax*
- *Nannopterum*

------------------------------------------------------------------------------------------------------------

**EcoClusterB** (no distinctive ecological trait)

- *Chauna*
- *Phoenicopterus*
- *Columbina*
- *Columba*
- *Leptotila*
- *Centropus*
- *Aramus*
- *Balearica*
- *Chroicocephalus*
- *Rostratula*
- *Charadrius*
- *Haematopus*
- *Eurypyga*
- *Leptoptilos*
- *Theristicus*
- *Ardea*
- *Ixobrychus*
- *Tigrisoma*
- *Balaeniceps*
- *Cathartes*
- *Vultur*
- *Sagittarius*
- *Caracara*
- *Corvus*
- *Arenaria*
- *Jynx*
- *Glareola*
- *Scopus*

- *Pterocles*
- *Syrrhaptes*
- *Bucorvus*

--------------------------------------------------------------------------------------------------------------

**EcoClusterC** (Habitat=Marine, Trophic Niche= Aquatic predator, Wing Propelled Swimming=1)

- *Uria*
- *Sterna*
- *Rynchops*
- *Phaethon*
- *Spheniscus*
- *Phoebastria*
- *Oceanites*
- *Pelagodroma*
- *Oceanodroma*
- *Puffinus*
- *Pelecanoides*
- *Pterodroma*
- *Pelecanus*
- *Fregata*
- *Morus*
- *Tringa*

--------------------------------------------------------------------------------------------------------------

**EcoClusterD** (Stooping=1)

- *Chaetura*
- *Hemiprocne*
- *Aegotheles*
- *Buteo*
- *Accipiter*
- *Falco*

--------------------------------------------------------------------------------------------------------------

**EcoClusterE** (Unassisted Hovering=1):

- *Topaza*
- *Phaethornis*
- *Archilochus*
- *Steatornis*
- *Alcedo*
- *Choloroceryle*
- *Chordeiles*
- *Caprimulgus*
- *Upupa*

--------------------------------------------------------------------------------------------------------------

**EcoClusterF** (Intermittent Flight=1, Primary Lifestyle=Insessorial, Habitat=Forest)

- *Treron*
- *Corythaeola*

- *Coccyzus*
- *Podargus*
- *Tyto*
- *Strix*
- *Colius*
- *Urocolius*
- *Leptosomus*
- *Trogon*
- *Coracias*
- *Todus*
- *Momota*
- *Picus*
- *Capito*
- *Micrastur*
- *Barnardius*
- *Psittrichas*
- *Psittacus*
- *Probosciger*
- *Nestor*
- *Neodrepanis*
- *Calyptomena*
- *Lepidocolaptes*
- *Ceratopipra*
- *Neopelma*
- *Schiffornis*
- *Oxyruncus*
- *Elaenia*
- *Climacteris*
- *Turdus*
- *Fringilla*
- *Acanthisitta*
- *Pitta*
- *Myrmornis*
- *Sclerurus*
- *Furnarius*
- *Piprites*
- *Bucco*

----------------------------------------------------------------------------------------------------------

**EcoClusterG** (Burst Flight=1, Trophic Level=Herbivore)

- *Eudromia*
- *Nothoprocta*
- *Tinamus*
- *Crypturellus*
- *Leipoa*
- *Numida*
- *Odontophorous*
- *Colinus*
- *Gallus*

- *Bonasa*
- *Ortalis*

-----------------------------------------------------------------------------------------------------------

**EcoClusterH** (Infrequent Flight=1 or Infrequent Flight=2)

- *Dromaius*
- *Apteryx*
- *Mesitornis*
- *Monias*
- *Opisthocomus*
- *Sarothrura*
- *Porphyrio*
- *Rallus*
- *Psophia*
- *Turnix*
- *Pedionomus*
- *Burhinus*
- *Atelornis*
- *Cariama*
- *Chunga*
- *Melanopareia*
- *Menura*

**Section S9:** Additional Galliformes (e.g., *Ortalis ruficauda*), Anseriformes and Gruiformes were grafted to the tree based on Chen et al. (Chen et al. 2021), Prum et al. (Prum et al. 2015), and Garcia-R et al. (Garcia-R et al. 2020), respectively. The clade Charadriiformes of Stiller et al. (Stiller et al. 2024) was replaced with taxa following Černý and Natale (Černý and Natale 2022). Additional Accipitriformes and Psittaciformes were grafted to the tree based on Catanach et al. (Catanach et al. 2025) and Smith et al. (Smith et al. 2024), respectively, and the divergence time between cormorants (i.e. *Nannopterum – Phalacrocorax*) was taken from Burga et al. (Burga et al. 2017). For suboscine passerine birds, additional Furnariides were added based on Oliveros et al. (Oliveros et al. 2019) and Tyrannides based on Prum et al. (Prum et al. 2015). Family and order data for each taxon were taken from AVONET2_eBird dataset (Tobias et al. 2022) and checked with and/or adapted to Stiller et al. (Stiller et al. 2024).

| **Humerus TP$_{prox}$** | | | **Ulna TP$_{prox}$** | | |
|---|---|---|---|---|---|
| Model | EIC | ΔEIC | Model | EIC | ΔEIC |
| BM | 1413.38 | 256.24 | BM | 1274.781692 | 272.8209261 |
| OU | 1166.84 | 9.7 | **OU** | **1001.960766** | **0** |
| **EB** | **1157.14** | **0** | EB | 1008.574717 | 6.613950798 |
| λ | 1183.80 | 26.66 | λ | 1043.958356 | 41.99758962 |
| **Humerus CSP** | | | **Ulna CSP** | | |
| Model | EIC | ΔEIC | Model | EIC | ΔEIC |
| BM | -7678.338397 | 1229.509706 | **BM** | **-9377.317785** | **0** |
| OU | -8556.130453 | 351.7176508 | OU | -8349.258528 | 1028.059256 |
| EB | -8548.838673 | 359.0094309 | EB | -8007.6336 | 1369.684184 |
| **λ** | **-8907.848104** | **0** | λ | -8782.304357 | 595.0134271 |
| **Humerus TP$_{dist}$** | | | **Ulna TP$_{dist}$** | | |
| Model | EIC | ΔEIC | Model | EIC | ΔEIC |
| BM | 1537.651365 | 379.5558604 | BM | 1597.931591 | 407.9941003 |
| **OU** | **1158.095504** | **0** | **OU** | **1189.937491** | **0** |
| EB | 1168.506346 | 10.41084118 | EB | 1219.893126 | 29.9556349 |
| λ | 1198.215874 | 40.12036963 | λ | 1249.473727 | 59.53623587 |

**Table S12.** Across the six studied anatomical regions, the best fitting evolutionary model was identified before running multivariate PGLSs. It was done by fitting Brownian Motion (BM), Ornstein-Uhlenbeck (OU), Early Burst (EB) and Pagel's lambda (λ) and measuring the Extended Information Criterion (EIC) scores. Models with the lowest EIC score (i.e. ΔEIC=0) are the best fitting, at each anatomical region.

| Comparison | Test stat | *p*-value | adjusted *p*-value |
|---|---|---|---|
| Eco-clusterA - Eco-clusterB | 0.064782 | 0.567433 | 1 |
| Eco-clusterA - Eco-clusterC | 0.156762 | 0.033966 | 0.951049 |
| Eco-clusterA - Eco-clusterD | 0.120593 | 0.096903 | 1 |
| Eco-clusterA - Eco-clusterE | 0.046786 | 0.782218 | 1 |
| Eco-clusterA - Eco-clusterF | 0.107246 | 0.171828 | 1 |
| Eco-clusterA - Eco-clusterG | 0.108388 | 0.166833 | 1 |
| Eco-clusterA - Eco-clusterH | 0.135516 | 0.061938 | 1 |
| Eco-clusterB - Eco-clusterC | 0.119331 | 0.112887 | 1 |
| Eco-clusterB - Eco-clusterD | 0.148168 | 0.031968 | 0.895105 |
| Eco-clusterB - Eco-clusterE | 0.070519 | 0.498501 | 1 |
| Eco-clusterB - Eco-clusterF | 0.132524 | 0.062937 | 1 |
| Eco-clusterB - Eco-clusterG | 0.151384 | 0.037962 | 1 |
| Eco-clusterB - Eco-clusterH | 0.178452 | 0.004995 | 0.13986 |
| Eco-clusterC - Eco-clusterD | 0.160512 | 0.018981 | 0.531469 |
| Eco-clusterC - Eco-clusterE | 0.115544 | 0.137862 | 1 |
| Eco-clusterC - Eco-clusterF | 0.238425 | 0.003996 | 0.111888 |
| Eco-clusterC - Eco-clusterG | 0.236193 | 0.003996 | 0.111888 |
| **Eco-clusterC - Eco-clusterH** | **0.221025** | **0.000999** | **0.027972** |
| Eco-clusterD - Eco-clusterE | 0.073771 | 0.470529 | 1 |
| Eco-clusterD - Eco-clusterF | 0.10398 | 0.185814 | 1 |
| Eco-clusterD - Eco-clusterG | 0.138522 | 0.043956 | 1 |
| Eco-clusterD - Eco-clusterH | 0.093139 | 0.275724 | 1 |
| Eco-clusterE - Eco-clusterF | 0.078217 | 0.413586 | 1 |
| Eco-clusterE - Eco-clusterG | 0.128513 | 0.077922 | 1 |
| Eco-clusterE - Eco-clusterH | 0.106686 | 0.156843 | 1 |
| Eco-clusterF - Eco-clusterG | 0.098041 | 0.242757 | 1 |
| Eco-clusterF - Eco-clusterH | 0.13729 | 0.04995 | 1 |
| Eco-clusterG - Eco-clusterH | 0.106565 | 0.18981 | 1 |

**Table S13.** pMANCOVA pairwise comparisons ran on proximal humeral data. For each comparison the test statistics, the p-value and the p-value corrected for multiple testing (Bonferroni correction) are shown. Pairwise comparisons yielding significance after Bonferroni correction are in bold.

| Comparison | Test stat | *p*-value | adjusted *p*-value |
|---|---|---|---|
| Eco-clusterA - Eco-clusterB | 0.170912063 | 0.12987013 | 1 |
| Eco-clusterA - Eco-clusterC | 0.210395644 | 0.048951049 | 1 |
| Eco-clusterA - Eco-clusterD | 0.203678449 | 0.04995005 | 1 |
| **Eco-clusterA - Eco-clusterE** | **0.357353468** | **0.000999001** | **0.027972028** |
| Eco-clusterA - Eco-clusterF | 0.198624317 | 0.071928072 | 1 |
| Eco-clusterA - Eco-clusterG | 0.168800173 | 0.127872128 | 1 |
| Eco-clusterA - Eco-clusterH | 0.194000172 | 0.065934066 | 1 |
| **Eco-clusterB - Eco-clusterC** | **0.361644159** | **0.000999001** | **0.027972028** |
| Eco-clusterB - Eco-clusterD | 0.183379965 | 0.082917083 | 1 |
| Eco-clusterB - Eco-clusterE | 0.344267979 | 0.007992008 | 0.223776224 |
| Eco-clusterB - Eco-clusterF | 0.067683089 | 0.818181818 | 1 |
| Eco-clusterB - Eco-clusterG | 0.150313418 | 0.184815185 | 1 |
| Eco-clusterB - Eco-clusterH | 0.256484492 | 0.007992008 | 0.223776224 |
| Eco-clusterC - Eco-clusterD | 0.314512595 | 0.007992008 | 0.223776224 |
| Eco-clusterC - Eco-clusterE | 0.437546352 | 0.001998002 | 0.055944056 |
| Eco-clusterC - Eco-clusterF | 0.423347218 | 0.001998002 | 0.055944056 |
| Eco-clusterC - Eco-clusterG | 0.334314594 | 0.008991009 | 0.251748252 |
| **Eco-clusterC - Eco-clusterH** | **0.362632411** | **0.000999001** | **0.027972028** |
| Eco-clusterD - Eco-clusterE | 0.183494598 | 0.084915085 | 1 |
| Eco-clusterD - Eco-clusterF | 0.168471487 | 0.114885115 | 1 |
| Eco-clusterD - Eco-clusterG | 0.179825126 | 0.123876124 | 1 |
| Eco-clusterD - Eco-clusterH | 0.203035447 | 0.052947053 | 1 |
| Eco-clusterE - Eco-clusterF | 0.368300257 | 0.001998002 | 0.055944056 |
| Eco-clusterE - Eco-clusterG | 0.279318737 | 0.015984016 | 0.447552448 |
| **Eco-clusterE - Eco-clusterH** | **0.374613855** | **0.000999001** | **0.027972028** |
| Eco-clusterF - Eco-clusterG | 0.124528986 | 0.305694306 | 1 |
| Eco-clusterF - Eco-clusterH | 0.244746114 | 0.011988012 | 0.335664336 |
| Eco-clusterG - Eco-clusterH | 0.222976408 | 0.06993007 | 1 |

**Table S14.** pMANCOVA pairwise comparisons ran on humeral diaphyseal data. For each comparison the test statistics, the p-value and the p-value corrected for multiple testing (Bonferroni correction) are shown. Pairwise comparisons yielding significance after Bonferroni correction are in bold.

| Comparison | Test stat | *p*-value | adjusted *p*-value |
|---|---|---|---|
| Eco-clusterA - Eco-clusterB | 0.027819 | 0.959041 | 1 |
| Eco-clusterA - Eco-clusterC | 0.099196 | 0.218781 | 1 |
| Eco-clusterA - Eco-clusterD | 0.120013 | 0.098901 | 1 |
| Eco-clusterA - Eco-clusterE | 0.064287 | 0.588412 | 1 |
| Eco-clusterA - Eco-clusterF | 0.094146 | 0.261738 | 1 |
| Eco-clusterA - Eco-clusterG | 0.077872 | 0.375624 | 1 |
| Eco-clusterA - Eco-clusterH | 0.137026 | 0.055944 | 1 |
| Eco-clusterB - Eco-clusterC | 0.146188 | 0.031968 | 0.895105 |
| Eco-clusterB - Eco-clusterD | 0.166209 | 0.016983 | 0.475524 |
| Eco-clusterB - Eco-clusterE | 0.088925 | 0.334665 | 1 |
| Eco-clusterB - Eco-clusterF | 0.136644 | 0.058941 | 1 |
| Eco-clusterB - Eco-clusterG | 0.124837 | 0.0999 | 1 |
| **Eco-clusterB - Eco-clusterH** | **0.215882** | **0.000999** | **0.027972** |
| Eco-clusterC - Eco-clusterD | 0.137953 | 0.04995 | 1 |
| Eco-clusterC - Eco-clusterE | 0.123375 | 0.108891 | 1 |
| Eco-clusterC - Eco-clusterF | 0.140838 | 0.056943 | 1 |
| Eco-clusterC - Eco-clusterG | 0.096154 | 0.251748 | 1 |
| Eco-clusterC - Eco-clusterH | 0.186604 | 0.010989 | 0.307692 |
| Eco-clusterD - Eco-clusterE | 0.061157 | 0.605395 | 1 |
| Eco-clusterD - Eco-clusterF | 0.111898 | 0.130869 | 1 |
| Eco-clusterD - Eco-clusterG | 0.084983 | 0.344655 | 1 |
| Eco-clusterD - Eco-clusterH | 0.0798 | 0.401598 | 1 |
| Eco-clusterE - Eco-clusterF | 0.084459 | 0.353646 | 1 |
| Eco-clusterE - Eco-clusterG | 0.083304 | 0.366633 | 1 |
| Eco-clusterE - Eco-clusterH | 0.11263 | 0.138861 | 1 |
| Eco-clusterF - Eco-clusterG | 0.069795 | 0.524476 | 1 |
| Eco-clusterF - Eco-clusterH | 0.185976 | 0.005994 | 0.167832 |
| Eco-clusterG - Eco-clusterH | 0.058169 | 0.659341 | 1 |

**Table S15.** pMANCOVA pairwise comparisons ran on distal humeral data. For each comparison the test statistics, the p-value and the p-value corrected for multiple testing (Bonferroni correction) are shown. Pairwise comparisons yielding significance after Bonferroni correction are in bold.

| **Comparison** | **Test stat** | ***p*-value** | **adjusted *p*-value** |
|---|---|---|---|
| Eco-clusterA - Eco-clusterB | 0.053569 | 0.753247 | 1 |
| Eco-clusterA - Eco-clusterC | 0.083698 | 0.362637 | 1 |
| Eco-clusterA - Eco-clusterD | 0.143809 | 0.047952 | 1 |
| Eco-clusterA - Eco-clusterE | 0.063715 | 0.621379 | 1 |
| Eco-clusterA - Eco-clusterF | 0.071349 | 0.517483 | 1 |
| Eco-clusterA - Eco-clusterG | 0.031262 | 0.92008 | 1 |
| Eco-clusterA - Eco-clusterH | 0.062203 | 0.619381 | 1 |
| Eco-clusterB - Eco-clusterC | 0.06528 | 0.602398 | 1 |
| Eco-clusterB - Eco-clusterD | 0.208981 | 0.001998 | 0.055944 |
| Eco-clusterB - Eco-clusterE | 0.112057 | 0.160839 | 1 |
| Eco-clusterB - Eco-clusterF | 0.114984 | 0.118881 | 1 |
| Eco-clusterB - Eco-clusterG | 0.159357 | 0.044955 | 1 |
| Eco-clusterB - Eco-clusterH | 0.153046 | 0.034965 | 0.979021 |
| Eco-clusterC - Eco-clusterD | 0.188059 | 0.00999 | 0.27972 |
| Eco-clusterC - Eco-clusterE | 0.119757 | 0.128871 | 1 |
| Eco-clusterC - Eco-clusterF | 0.121501 | 0.111888 | 1 |
| Eco-clusterC - Eco-clusterG | 0.166925 | 0.020979 | 0.587413 |
| Eco-clusterC - Eco-clusterH | 0.136285 | 0.070929 | 1 |
| Eco-clusterD - Eco-clusterE | 0.105408 | 0.205794 | 1 |
| Eco-clusterD - Eco-clusterF | 0.139922 | 0.053946 | 1 |
| Eco-clusterD - Eco-clusterG | 0.120321 | 0.114885 | 1 |
| Eco-clusterD - Eco-clusterH | 0.091236 | 0.297702 | 1 |
| Eco-clusterE - Eco-clusterF | 0.087435 | 0.356643 | 1 |
| Eco-clusterE - Eco-clusterG | 0.068041 | 0.562438 | 1 |
| Eco-clusterE - Eco-clusterH | 0.075425 | 0.456543 | 1 |
| Eco-clusterF - Eco-clusterG | 0.116334 | 0.13986 | 1 |
| Eco-clusterF - Eco-clusterH | 0.094661 | 0.270729 | 1 |
| Eco-clusterG - Eco-clusterH | 0.08372 | 0.362637 | 1 |

**Table S16.** pMANCOVA pairwise comparisons ran on proximal ulnar data. For each comparison the test statistics, the p-value and the p-value corrected for multiple testing (Bonferroni correction) are shown. Pairwise comparisons yielding significance after Bonferroni correction are in bold.

| Comparison | Test stat | *p*-value | adjusted *p*-value |
|---|---|---|---|
| Eco-clusterA - Eco-clusterB | 0.083576 | 0.471528 | 1 |
| Eco-clusterA - Eco-clusterC | 0.166023 | 0.031968 | 0.895105 |
| Eco-clusterA - Eco-clusterD | 0.122914 | 0.132867 | 1 |
| Eco-clusterA - Eco-clusterE | 0.166593 | 0.01998 | 0.559441 |
| Eco-clusterA - Eco-clusterF | 0.062508 | 0.701299 | 1 |
| Eco-clusterA - Eco-clusterG | 0.117377 | 0.192807 | 1 |
| Eco-clusterA - Eco-clusterH | 0.190687 | 0.018981 | 0.531469 |
| Eco-clusterB - Eco-clusterC | 0.149532 | 0.053946 | 1 |
| Eco-clusterB - Eco-clusterD | 0.21796 | 0.003996 | 0.111888 |
| Eco-clusterB - Eco-clusterE | 0.156386 | 0.035964 | 1 |
| Eco-clusterB - Eco-clusterF | 0.077461 | 0.54046 | 1 |
| Eco-clusterB - Eco-clusterG | 0.131194 | 0.126873 | 1 |
| Eco-clusterB - Eco-clusterH | 0.210908 | 0.002997 | 0.083916 |
| **Eco-clusterC - Eco-clusterD** | **0.258919** | **0.000999** | **0.027972** |
| Eco-clusterC - Eco-clusterE | 0.17069 | 0.031968 | 0.895105 |
| Eco-clusterC - Eco-clusterF | 0.097164 | 0.32967 | 1 |
| Eco-clusterC - Eco-clusterG | 0.151655 | 0.057942 | 1 |
| Eco-clusterC - Eco-clusterH | 0.159576 | 0.032967 | 0.923077 |
| Eco-clusterD - Eco-clusterE | 0.161424 | 0.040959 | 1 |
| Eco-clusterD - Eco-clusterF | 0.206371 | 0.004995 | 0.13986 |
| **Eco-clusterD - Eco-clusterG** | **0.272819** | **0.000999** | **0.027972** |
| **Eco-clusterD - Eco-clusterH** | **0.274742** | **0.000999** | **0.027972** |
| Eco-clusterE - Eco-clusterF | 0.163059 | 0.032967 | 0.923077 |
| Eco-clusterE - Eco-clusterG | 0.199006 | 0.006993 | 0.195804 |
| Eco-clusterE - Eco-clusterH | 0.17911 | 0.015984 | 0.447552 |
| Eco-clusterF - Eco-clusterG | 0.114152 | 0.213786 | 1 |
| Eco-clusterF - Eco-clusterH | 0.191792 | 0.011988 | 0.335664 |
| Eco-clusterG - Eco-clusterH | 0.08594 | 0.44955 | 1 |

**Table S17.** pMANCOVA pairwise comparisons ran on distal ulnar data. For each comparison the test statistics, the p-value and the p-value corrected for multiple testing (Bonferroni correction) are shown. Pairwise comparisons yielding significance after Bonferroni correction are in bold.

| Humerus $TP_{prox}$ | | | | | |
|---|---|---|---|---|---|
| DF1 | | DF2 | | DF3 | |
| $R^2$ | *p* | $R^2$ | *p* | $R^2$ | *p* |
| 0.12 | <0.001 | 0.87 | <0.001 | 0.29 | <0.001 |
| **Humerus CSP** | | | | | |
| DF1 | | DF2 | | DF3 | |
| $R^2$ | *p* | $R^2$ | *p* | $R^2$ | *p* |
| 0.35 | <0.001 | 0.20 | <0.001 | 0.91 | 0.7745 |
| **Humerus $TP_{dist}$** | | | | | |
| DF1 | | DF2 | | DF3 | |
| $R^2$ | *p* | $R^2$ | *p* | $R^2$ | *p* |
| 0.78 | <0.001 | 0.374 | <0.001 | 0.06 | 0.004 |
| **Ulna $TP_{prox}$** | | | | | |
| DF1 | | DF2 | | DF3 | |
| $R^2$ | *p* | $R^2$ | *p* | $R^2$ | *p* |
| 0.86 | <0.001 | 0.57 | <0.001 | / | 0.1519 |
| **Ulna CSP** | | | | | |
| DF1 | | DF2 | | DF3 | |
| $R^2$ | *p* | $R^2$ | *p* | $R^2$ | *p* |
| 0.80 | <0.001 | 0.77 | <0.001 | 0.11 | <0.001 |
| **Ulna $TP_{dist}$** | | | | | |
| DF1 | | DF2 | | DF3 | |
| $R^2$ | p | $R^2$ | p | $R^2$ | *p* |
| 0.35 | <0.001 | 0.81 | <0.001 | 0.015 | 0.1737 |

**Table S18.** Across the six studied anatomical regions, the first three DFs deriving from DFA were tested for allometric effects through a correlation analysis with BMp.

| **Humeral TP$_{prox}$** | | | | | | |
|---|---|---|---|---|---|---|
| **Eco-clusterC - Eco-clusterH** | | | | | | |
| DF1 | DF2 | DF3 | DF4 | DF5 | DF6 | DF7 |
| **0.003** | 0.748 | 0.798 | 0.315 | 0.395 | 0.788 | 0.896 |
| **Humeral CSP** | | | | | | |
| **Eco-clusterA - Eco-clusterE** | | | | | | |
| DF1 | DF2 | DF3 | DF4 | DF5 | DF6 | DF7 |
| **0.007** | 0.226 | **0.015** | 0.707 | **0.006** | 0.225 | 0.544 |
| **Eco-clusterB - Eco-clusterC** | | | | | | |
| DF1 | DF2 | DF3 | DF4 | DF5 | DF6 | DF7 |
| **0.001** | **0.022** | 0.463 | 0.733 | 0.57 | 0.206 | 0.721 |
| **Eco-clusterC - Eco-clusterH** | | | | | | |
| DF1 | DF2 | DF3 | DF4 | DF5 | DF6 | DF7 |
| **0.014** | **0.044** | 0.32 | 0.803 | 0.814 | 0.653 | 0.793 |
| **Eco-clusterE - Eco-clusterH** | | | | | | |
| DF1 | DF2 | DF3 | DF4 | DF5 | DF6 | DF7 |
| **0.005** | **0.015** | 0.18 | 0.781 | 0.082 | 0.41 | 0.608 |
| **Humeral TP$_{dist}$** | | | | | | |
| **Eco-clusterB - Eco-clusterH** | | | | | | |
| DF1 | DF2 | DF3 | DF4 | DF5 | DF6 | DF7 |
| **0.006** | 0.742 | 0.315 | 0.985 | 0.473 | 0.702 | 0.356 |
| **Ulnar TP$_{dist}$** | | | | | | |
| **Eco-clusterC - Eco-clusterD** | | | | | | |
| DF1 | DF2 | DF3 | DF4 | DF5 | DF6 | DF7 |
| **0.002** | 0.313 | 0.297 | 0.966 | 0.681 | 0.952 | 0.9 |
| **Eco-clusterD - Eco-clusterG** | | | | | | |
| DF1 | DF2 | DF3 | DF4 | DF5 | DF6 | DF7 |
| **0.022** | 0.246 | 0.548 | 0.916 | 0.969 | 0.644 | 0.827 |
| **Eco-clusterD - Eco-clusterH** | | | | | | |
| DF1 | DF2 | DF3 | DF4 | DF5 | DF6 | DF7 |
| **0.002** | 0.483 | 0.682 | 0.751 | 0.705 | 0.959 | 0.595 |

**Table S19.** For the studied anatomical regions, we identified which DFs mainly contribute to set apart the eco-clusters arising as significantly different from pairwise comparisons (see Tables S13-S19). It was done by DFs *vs.* eco-clusters pANOVAs.

### Humerus $TP_{prox}$

| Variable | DF1 | DF2 | DF3 | DF4 | DF5 | DF6 | DF7 |
|---|---|---|---|---|---|---|---|
| $DA_{prox}$ | 0.039 | -0.166 | -0.129 | -0.026 | -0.315 | -0.274 | 0.728 |
| log10-$TbTh_{prox}$ | 1.307 | 0.401 | -1.644 | -0.448 | 0.61 | -0.54 | -0.131 |
| log10-$BVTV_{prox}$ | -1.827 | -0.73 | 0.257 | -0.051 | -0.957 | 0.747 | -0.007 |
| log10-$ConnD_{prox}$ | -0.247 | 0.309 | -0.325 | -0.007 | 1.402 | -0.545 | 0.329 |
| log10-$NodDenAver_{prox}$ | -0.982 | 1.913 | 1.098 | -3.119 | -5.903 | 7.099 | -4.896 |
| log10-$NodDenMed_{prox}$ | 2.13 | -2.401 | -1.73 | 3.555 | 6.088 | -6.909 | 4.191 |
| $TrabLenMean_{prox}$ | -0.714 | 0.134 | 0.565 | 0.268 | 0.658 | 0.333 | -0.423 |
| $TrabTortAver_{prox}$ | 0.273 | 0.236 | 0.314 | 0.014 | 0.402 | 1.377 | 0.961 |
| $TrabTortMed_{prox}$ | 0.32 | 0.177 | 0.109 | 0.323 | -0.305 | -1.016 | -0.754 |
| $FD_{prox}$ | 1.595 | 0.979 | -0.12 | 1.041 | 0.05 | 0.414 | -0.179 |

### Humerus CSP

| Variable | DF1 | DF2 | DF3 | DF4 | DF5 | DF6 | DF7 |
|---|---|---|---|---|---|---|---|
| log10-$TotAr_{50}$ | 1.949 | -1.772 | -0.5 | 5.893 | -6.549 | -13.128 | 1.949 |
| $Cg_{50}$ | 3.917 | -3.215 | -1.283 | 7.645 | -3.133 | 10.195 | 3.917 |
| log10-$TotAr_{Aver}$ | 2.557 | 0.281 | -5.018 | -6.071 | -6.458 | 14.043 | 2.557 |
| $Cg_{Aver}$ | -1.627 | 2.587 | 3.115 | -2.332 | 4.616 | -10.462 | -1.627 |
| log10-$CSA_{50}$ | -5.675 | 8.414 | 2.838 | -10.155 | 0.504 | -28.65 | -5.675 |
| log10-$CSA_{Aver}$ | 3.867 | -6.954 | -10.913 | -5.7 | -11.616 | 33.282 | 3.867 |
| log10-$Imin_{50}$ | 6 | 0.619 | -6.571 | 0.908 | 6.961 | 17.399 | 6 |
| log10-$Imin_{Aver}$ | -7.858 | -3.326 | 11.297 | 6.501 | 6.289 | -14.533 | -7.858 |
| log10-$Imax_{50}$ | 5.691 | 0.602 | -6.239 | 0.844 | 6.559 | 16.498 | 5.691 |
| log10-$Imax_{Aver}$ | -7.427 | -3.167 | 10.673 | 6.154 | 5.996 | -13.714 | -7.427 |
| log10-$Zpol_{50}$ | -10.171 | -14.352 | 11.674 | 6.331 | -6.381 | 3.267 | -10.171 |
| log10-$Zpol_{Aver}$ | 11.292 | 20.285 | -8.559 | -5.404 | 3.964 | -14.359 | 11.292 |
| log10-$Ct.Th_{50}$ | 0.163 | -1.137 | 0.272 | 0.437 | 0.5 | -0.375 | 0.163 |
| $FToHStrStrength_{50}$ | -0.02 | 0.009 | -0.15 | 0.128 | -0.167 | -0.036 | -0.02 |
| log10-$CSS_{Aver}$ | 0.155 | -1.007 | -0.516 | 0.302 | 2.3 | 1.411 | 0.155 |
| log10-$CSS_{50}$ | -0.408 | 0.704 | 0.131 | -0.931 | -2.607 | -1.431 | -0.408 |

### Humerus $TP_{dist}$

| Variable | DF1 | DF2 | DF3 | DF4 | DF5 | DF6 | DF7 |
|---|---|---|---|---|---|---|---|
| $DA_{prox}$ | -0.262 | -0.484 | -0.159 | 0.014 | 0.185 | -0.334 | 0.846 |
| log10-$TbTh_{prox}$ | -0.218 | -0.057 | -0.648 | -0.774 | 0.05 | 1.275 | 0.152 |
| log10-$BVTV_{prox}$ | -0.112 | 1.273 | 0.041 | 0.416 | 0.183 | -1.043 | 0.335 |
| log10-$ConnD_{prox}$ | 0.126 | -0.118 | -0.861 | 0.976 | -0.399 | 0.746 | -0.029 |
| log10-$NodDenAver_{prox}$ | -7.025 | 9.676 | 7.422 | -9.748 | -6.951 | -2.866 | 6.44 |
| log10-$NodDenMed_{prox}$ | 6.324 | -10.372 | -7.943 | 8.426 | 6.201 | 3.306 | -6.995 |
| $TrabLenMean_{prox}$ | 0.421 | -0.296 | -0.471 | 0.002 | -1.206 | 0.206 | -0.164 |
| $TrabTortAver_{prox}$ | 0.416 | -0.054 | -0.505 | -0.04 | -0.068 | -0.853 | -0.216 |
| $TrabTortMed_{prox}$ | -0.368 | -0.498 | 0.146 | 0.151 | 0.565 | 1.07 | -0.032 |
| $FD_{prox}$ | 0.448 | -1.201 | -0.488 | -1.61 | -0.009 | 0.283 | -0.95 |

**Table S20.** For each of the DFs from DFA on humeral structural levels, the relative contribution of original structural variables is shown through standardized coefficients.

| Ulnar $TP_{prox}$ | | | | | | | |
|---|---|---|---|---|---|---|---|
| **Variable** | **DF1** | **DF2** | **DF3** | **DF4** | **DF5** | **DF6** | **DF7** |
| $DA_{prox}$ | -0.242 | 0.496 | -0.29 | 0.595 | -0.66 | -0.076 | -0.175 |
| log10-$TbTh_{prox}$ | -0.561 | -0.751 | 0.825 | 1.082 | -0.219 | -0.03 | 0.965 |
| log10-$BVTV_{prox}$ | 0.861 | 1.536 | -0.226 | -1.543 | -0.269 | 0.806 | -0.044 |
| log10-$ConnD_{prox}$ | -0.077 | -1.239 | -0.28 | 1.136 | 0.841 | -0.876 | 1.345 |
| log10-$NodDenAver_{prox}$ | -7.81 | -2.203 | 5.561 | -8.2 | -2.576 | -10.365 | -0.878 |
| log10-$NodDenMed_{prox}$ | 7.842 | 1.36 | -4.953 | 8.306 | 1.632 | 10.201 | 0.602 |
| $TrabLenMean_{prox}$ | -0.52 | -0.831 | -0.488 | -0.449 | 0.088 | -0.364 | 0.193 |
| $TrabTortAver_{prox}$ | -0.15 | 0.099 | 0.474 | 0.306 | 0.244 | -1.166 | 0.567 |
| $TrabTortMed_{prox}$ | -0.179 | -0.633 | -1.057 | -0.123 | -0.328 | 0.181 | -0.199 |
| $FD_{prox}$ | -0.874 | -0.939 | 0.318 | 0.114 | -0.212 | -1.055 | -0.059 |
| **Ulnar CSP** | | | | | | | |
| **Variable** | **DF1** | **DF2** | **DF3** | **DF4** | **DF5** | **DF6** | **DF7** |
| log10-$TotAr_{50}$ | 26.731 | 4.997 | -38.678 | -34.135 | -8.025 | -5.468 | 16.937 |
| $Cg_{50}$ | 9.824 | 1.733 | -13.694 | -12.375 | -3.182 | -2.149 | 8.112 |
| log10-$TotAr_{Aver}$ | -25.9 | 32.719 | 33.108 | -16.118 | -5.124 | -10.498 | 0.278 |
| $Cg_{Aver}$ | -2.497 | -12.585 | 10.313 | 19.31 | 5.504 | 7.017 | -8.361 |
| log10-$CSA_{50}$ | -9.18 | 1.769 | 24.194 | 20.251 | 3.242 | 13.143 | -19.852 |
| log10-$CSA_{Aver}$ | -11.684 | 46.104 | -17.965 | -62.209 | -11.205 | -38.009 | 24.036 |
| log10-$Imin_{50}$ | -11.716 | -0.13 | 5.758 | 3.362 | -1.276 | -13.427 | 5.269 |
| log10-$Imin_{Aver}$ | 15.07 | -42.752 | -1.113 | 37.094 | 12.331 | 32.565 | -21.176 |
| log10-$Imax_{50}$ | -10.962 | -0.118 | 5.367 | 3.116 | -1.122 | -12.592 | 4.877 |
| log10-$Imax_{Aver}$ | 14.119 | -40.187 | -1.02 | 34.891 | 11.521 | 30.591 | -19.8 |
| log10-$Zpol_{50}$ | 10.353 | -5.06 | 5.185 | -2.312 | 0.589 | 6.515 | -7.823 |
| log10-$Zpol_{Aver}$ | 5.514 | 0.063 | -15.058 | 17.874 | 1.496 | -2.875 | 16.344 |
| log10-$Ct.Th_{50}$ | 0.414 | -0.272 | -1.077 | -2.145 | -2.888 | 0.828 | 1.697 |
| log10-$CSS_{Aver}$ | -1.185 | -0.816 | 0.822 | 1.39 | -1.982 | -0.031 | 2.946 |
| log10-$CSS_{50}$ | 1.225 | 0.127 | -1.231 | -1.285 | 2.384 | 0.489 | -2.198 |
| **Ulnar $TP_{dist}$** | | | | | | | |
| **Variable** | **DF1** | **DF2** | **DF3** | **DF4** | **DF5** | **DF6** | **DF7** |
| $DA_{prox}$ | 0.007 | 0.172 | -0.05 | 0.478 | 0.492 | -0.278 | 0.007 |
| log10-$TbTh_{prox}$ | 0.734 | 0.128 | -0.413 | 0.798 | -0.017 | -0.526 | 0.734 |
| log10-$BVTV_{prox}$ | -0.456 | -0.317 | 0.386 | -0.059 | -1.17 | 0.461 | -0.456 |
| log10-$ConnD_{prox}$ | -0.919 | -0.452 | -0.668 | -0.52 | 0.34 | -0.527 | -0.919 |
| log10-$NodDenAver_{prox}$ | 7.775 | -6.923 | -15.859 | 1.454 | -5.801 | -6.793 | 7.775 |
| log10-$NodDenMed_{prox}$ | -6.404 | 6.62 | 15.615 | -1.532 | 6.068 | 6.894 | -6.404 |
| $TrabLenMean_{prox}$ | 0.118 | 0.174 | -0.541 | -1.317 | 0.015 | 0.146 | 0.118 |
| $TrabTortAver_{prox}$ | 0.703 | -0.276 | 0.262 | 0.069 | 0.316 | 0.421 | 0.703 |
| $TrabTortMed_{prox}$ | -0.564 | 0.121 | 0.288 | -0.057 | -0.079 | -0.919 | -0.564 |
| $FD_{prox}$ | 2.106 | 0.18 | 0.48 | -0.435 | 0.463 | -0.248 | 2.106 |

**Table S21.** For each of the DFs from DFA on ulnar structural levels, the relative contribution of original structural variables is shown through standardized coefficients.

**References:**


Alfieri, F., L. Botton-Divet, J. A. Nyakatura, and E. Amson. 2022. Integrative approach uncovers new patterns of ecomorphological convergence in slow arboreal xenarthrans. J Mamm Evol, doi: 10.1007/s10914-021-09590-5.

Alfieri, F., L. Botton-Divet, J. Wölfer, J. A. Nyakatura, and E. Amson. 2023. A macroevolutionary common-garden experiment reveals differentially evolvable bone organization levels in slow arboreal mammals. Communications Biology 6(1):995.

Alfieri, F., A. Veneziano, D. Panetta, P. A. Salvadori, E. Amson, and D. Marchi. 2025. The relationship between primate distal fibula trabecular architecture and arboreality, phylogeny and size. Journal of Anatomy 00:1–29.

Amson, E. 2019. Overall bone structure as assessed by slice-by-slice profile. Evol Biol 46:343–348.

Amson, E., P. Arnold, A. H. van Heteren, A. Canoville, and J. A. Nyakatura. 2017. Trabecular architecture in the forelimb epiphyses of extant xenarthrans (Mammalia). Front Zool 14:52.

Bernsen, J. 1986. Dynamic thresholding of grey-level images. Proceedings of the 8th International conference on pattern recognition, Paris, France 1251-1255.

Bishop, P. J., S. A. Hocknull, C. J. Clemente, J. R. Hutchinson, A. A. Farke, B. R. Beck, R. S. Barret, and D. G. Lloyd. 2018. Cancellous bone and theropod dinosaur locomotion. Part I—an examination of cancellous bone architecture in the hindlimb bones of theropods. PeerJ, 6:e5778.

Burga, A., W. Wang, E. Ben-David, P. C. Wolf, A. M. Ramey, C. Verdugo, K. Lyons, P. G. Parker, and L. Kruglyak. 2017. A genetic signature of the evolution of loss of flight in the Galapagos cormorant. Science 356:eaal3345.

Catanach, T. A., M. R. Halley, and S. Pirro. 2025. Enigmas no longer: using ultraconserved elements to place several unusual hawk taxa and address the non-monophyly of the genus Accipiter (Accipitriformes: Accipitridae). Biological journal of the Linnean Society (2024) 144.

Černý, D., and R. Natale. 2022. Comprehensive taxon sampling and vetted fossils help clarify the time tree of shorebirds (Aves, Charadriiformes). Molecular Phylogenetics and Evolution 177:107620.

Chen, D. H., P. A. Hosner, D. L. Dittmann, J. P. O'Neill, E. L. Braun, and R. T. Kimball. 2021. Divergence time estimation of Galliformes based on the best gene shopping scheme of ultraconserved elements. BMC Ecology and Evolution 21:1–15.

de Margerie, E., S. Sanchez, J. Cubo, and J. Castanet. 2005. Torsional resistance as a principal component of the structural design of long bones: comparative multivariate evidence in birds. Anat Rec A Discov Mol Cell Evol Biol 282:49–66.

Domander, R., A. A. Felder, and M. Doube. 2021. BoneJ2 - refactoring established research software. Wellcome Open Research 6:37.

Doube, M., M. M. Kłosowski, I. Arganda-Carreras, F. P. Cordelières, R. P. Dougherty, J. S. Jackson, B. Schmid, J. R. Hutchinson, and S. J. Shefelbine. 2010. BoneJ: Free and extensible bone image analysis in ImageJ. Bone 47:1076–1079.

Garcia-R, J. C., E. M. Lemmon, A. R. Lemmon, and N. French. 2020. Phylogenomic reconstruction sheds light on new relationships and timescale of rails (Aves: Rallidae) evolution. Diversity 12(2), 70.

Habib, M. B., and C. B. Ruff. 2008. The effects of locomotion on the structural characteristics of avian limb bones. Zoological Journal of the Linnean Society 153:601–624.

Keith, S., C. W. Benson, and M. P. Stuart. 1970. The genus *Sarothrura* (Aves, Rallidae). Bulletin of the AMNH 143, article 1.

Kemp, A. C., and M. I. Kemp. 1980. The biology of the southern ground hornbill <i>Bucorvus leadbeateri</I> (Vigors)(Aves: Bucerotidae). Annals of the Transvaal Museum 32(4):65–100.

Kivell, T. L., M. M. Skinner, R. Lazenby, and J.-J. Hublin. 2011. Methodological considerations for analyzing trabecular architecture: an example from the primate hand. J. Anat. 218:209–225.

Laurin, M. 2004. The evolution of body size, Cope's rule and the origin of amniotes. Syst Biol 53:594–622.

Lukova, A., C. J. Dunmore, Z. J. Tsegai, S. Bachmann, A. Synek, and M. M. Skinner. 2024. Technical note: Does scan resolution or downsampling impact the analysis of trabecular bone architecture? American Journal of Biological Anthropology 185:e25023.

Mielke, M., J. Wölfer, P. Arnold, A. H. van Heteren, E. Amson, and J. A. Nyakatura. 2018. Trabecular architecture in the sciuromorph femoral head: allometry and functional adaptation. Zool Lett 4:10.

Oliveros, C. H., D. J. Field, D. T. Ksepka, F. K. Barker, A. Aleixo, M. J. Andersen, P. Alström, B. W. Benz, E. L. Braun, M. J. Braun, G. A. Bravo, R. T. Brumfield, R. T. Chesser, Claramunt S., Cracraft J., Cuervo A. M., Derryberry E. P., T. C. Glenn, M. G. Harvey, P. A. Hosner, L. Joseph, R. T. Kimbal, A. L. Mack, C. M. Miskelly, A. T. Peterson, M. B. Robbins, F. H. Sheldon, L. F. Silveira, B. T. Smith, N. D. White, R. G. Moylegg, and B. C. Faircloth. 2019. Earth history and the passerine superradiation. Proc. Natl. Acad. Sci. U.S.A 116 (16):7916-7925,.

Pau, G., F. Fuchs, O. Sklyar, M. Boutros, and W. Huber. 2010. EBImage - an R package for image processing with applications to cellular phenotypes. Bioinformatics 26(7):979–981.

Prum, R. O., J. S. Berv, A. Dornburg, D. J. Field, J. P. Townsend, E. M. Lemmon, and A. R. Lemmon. 2015. A comprehensive phylogeny of birds (Aves) using targeted next-generation DNA sequencing. Nature 526(7574):569–573.

Rand, A. L. 1936. The distribution and habits of Madagascar birds. Bull. Amer. Mus. Nat. Hist. 72:143–499.

Schindelin, J., I. Arganda-Carreras, E. Frise, V. Kaynig, M. Longair, T. Pietzsch, S. Preibisch, C. Rueden, S. Saalfeld, B. Schmid, J.-Y. Tinevez, D. J. White, V. Hartenstein, K. Eliceiri, P. Tomancak, and A. Cardona. 2012. Fiji: an open-source platform for biological-image analysis. Nat. Methods 9:676–682.

Smith, B. T., G. Thom, and L. Joseph. 2024. Revised evolutionary and taxonomic synthesis for parrots (Order: Psittaciformes) guided by phylogenomic analysis. Bulletin of the American Museum of Natural History 2024(468):1-87.

Stiles, F. G., and B. Whitney. 1983. Notes on the behavior of the Costa Rican Sharpbill (*Oxyruncus cristatus frater*). The Auk 100(1):117–125.

Stiller, J., S. Feng, A.-A. Chowdhury, I. Rivas-González, D. A. Duchêne, Q. Fang, Y. Deng, A. Kozlov, A. Stamatakis, S. Claramunt, J. M. T. Nguyen, S. Y. W. Ho, B. C. Faircloth, J. Haag, P. Houde, J. Cracraft, M. Balaban, U. Mai, G. Chen, R. Gao, C. Zhou, Y. Xie, Z. Huang, Z. Cao, Z. Yan, H. A. Ogilvie, L. Nakhleh, B. Lindow, B. Morel, J. Fjeldså, P. A. Hosner, R. R. da Fonseca, B. Petersen, J. A. Tobias, T. Székely, J. D. Kennedy, A. H. Reeve, A. Liker, M. Stervander, A. Antunes, D. T. Tietze, M. F. Bertelsen, F. Lei, C. Rahbek, G. R. Graves, M. H. Schierup, T. Warnow, E. L. Braun, M. T. P. Gilbert, E. D. Jarvis, S. Mirarab, and G. Zhang. 2024. Complexity of avian evolution revealed by family-level genomes. Nature 629:851–860. Nature Publishing Group.

Tobalske, B. W. 2001. Morphology, velocity, and intermittent flight in birds. American Zoologist 41.2:177–187.

Tobalske, B. W., and K. P. Dial. 1994. Neuromuscular control and kinematics of intermittent flight in budgerigars (*Melopsittacus undulatus*). Journal of experimental biology 187(1):1–18.

Tobias, J. A., C. Sheard, A. L. Pigot, A. J. M. Devenish, J. Yang, F. Sayol, M. H. C. Neate-Clegg, N. Alioravainen, T. L. Weeks, R. A. Barber, P. A. Walkden, H. E. A. MacGregor, S. E. I. Jones, C. Vincent, A. G. Phillips, N. . M. Marples, F. A. Montaño-Centellas, V. Leandro-Silva, S. Claramunt, B. Darski, B. G. Freeman, T. P. Bregman, C. R. Cooney, E. C. Hughes, E. J. R. Capp, Z. K. Varley, N. R. Friedman, H. Korntheuer, A. Corrales-Vargas, C. H. Trisos, B. C. Weeks, D. M. Hanz, T. Töpfer, G. A. Bravo, V. Remeš, L. Nowak, L. S. Carneiro, A. J. Moncada R., B. Matysioková, D. T. Baldassarre, A. Martínez-Salinas, J. D. Wolfe, P. M. Chapman, B. G.

Daly, M. C. Sorensen, A. Neu, M. A. Ford, R. J. Mayhew, L. Fabio Silveira, D. J. Kelly, N. N. D. Annorbah, H. S. Pollock, A. M. Grabowska-Zhang, J. P. McEntee, J. Carlos T. Gonzalez, C. G. Meneses, M. C. Muñoz, L. L. Powell, G. A. Jamie, T. J. Matthews, O. Johnson, G. R. R. Brito, K. Zyskowski, R. Crates, M. G. Harvey, M. Jurado Zevallos, P. A. Hosner, T. Bradfer-Lawrence, J. M. Maley, F. G. Stiles, H. S. Lima, K. L. Provost, M. Chibesa, M. Mashao, J. T. Howard, E. Mlamba, M. A. H. Chua, B. Li, M. I. Gómez, N. C. García, M. Päckert, J. Fuchs, J. R. Ali, E. P. Derryberry, M. L. Carlson, R. C. Urriza, K. E. Brzeski, D. M. Prawiradilaga, M. J. Rayner, E. T. Miller, R. C. K. Bowie, R.-M. Lafontaine, R. P. Scofield, Y. Lou, L. Somarathna, D. Lepage, M. Illif, E. L. Neuschulz, M. Templin, D. M. Dehling, J. C. Cooper, O. S. G. Pauwels, K. Analuddin, J. Fjeldså, N. Seddon, P. R. Sweet, F. A. J. DeClerck, L. N. Naka, J. D. Brawn, A. Aleixo, K. Böhning-Gaese, C. Rahbek, S. A. Fritz, G. H. Thomas, and M. Schleuning. 2022. AVONET: morphological, ecological and geographical data for all birds. Ecology Letters 25:581–597.

Veneziano, A., M. Cazenave, F. Alfieri, D. Panetta, and D. Marchi. 2021. Novel strategies for the characterization of cancellous bone morphology: Virtual isolation and analysis. Am J Phys Anthropol 175(4):920-930.

Witherby, H. F., F. C. R. Jourdain, N. F. Ticehurst, and B. W. Tucker. 1938. P. *in* The Handbook of British Birds. Witherby H.F and Witherby G (eds). London (Ltd).

You, K., and D. Shung. 2022. Rdimtools: An R package for dimension reduction and intrinsic dimension estimation. Software Impacts 14(1):100414.
