## Supplementary figures and images for "Constrained variation in the internal architecture of avian wing bones"

### Further boxplots of humeral diaphyseal parameters of interest (identified as detailed in the Main Text)

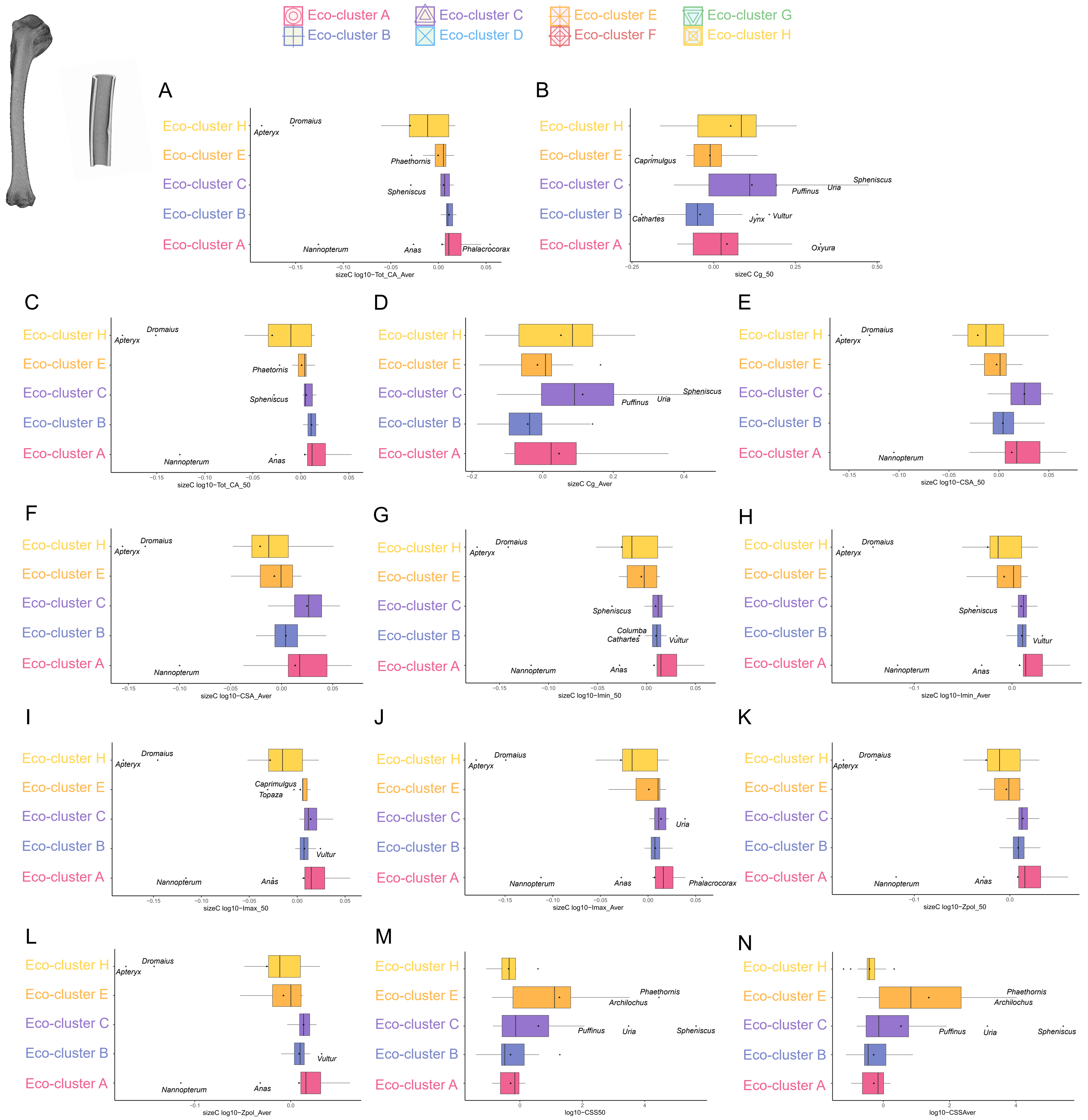

### Results for distal humeral structural parameters.

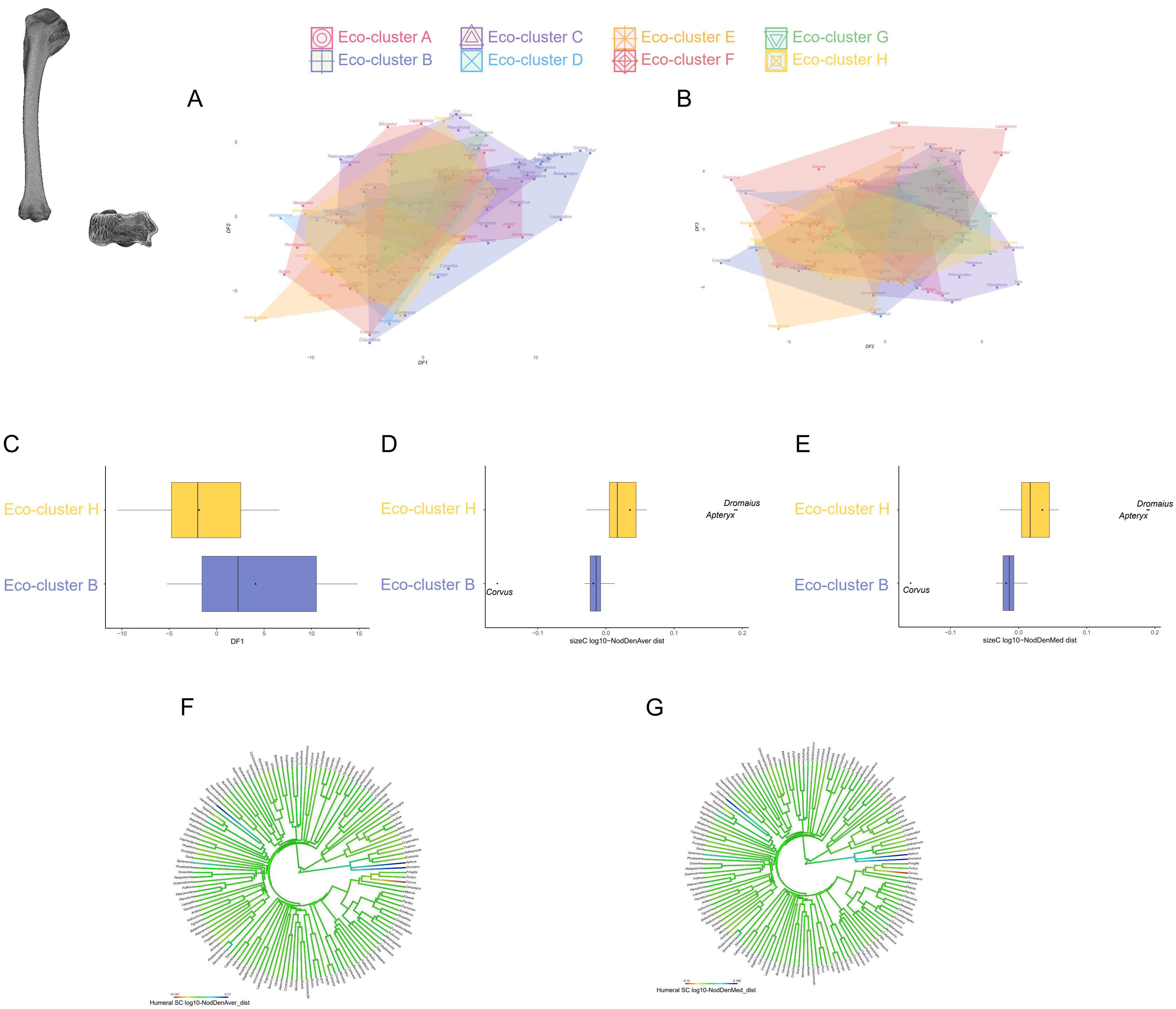

### Results for distal ulnar structural parameters.

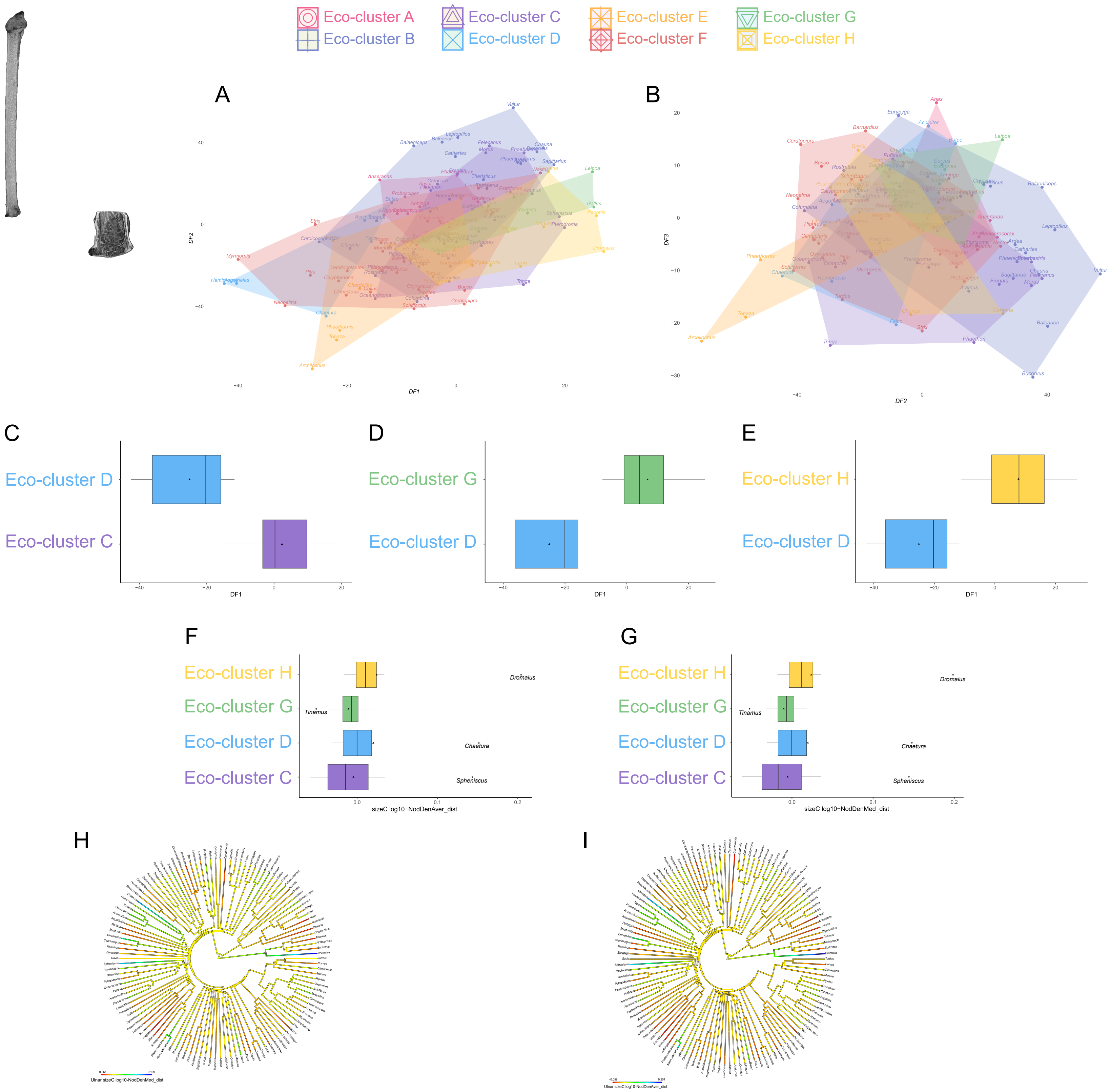

### Results for humeral diaphyseal structural parameters.

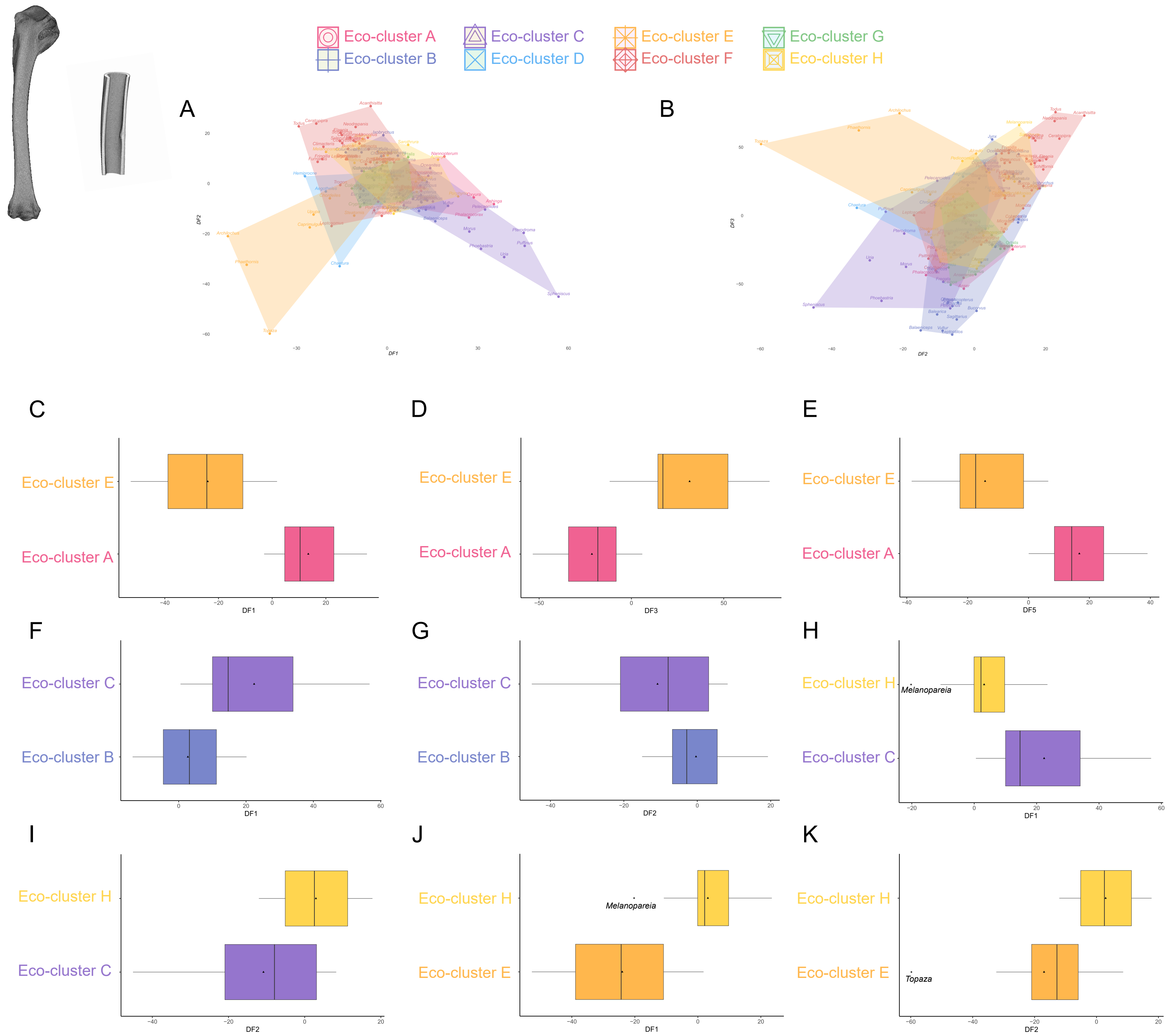

### Results for proximal humeral structural parameters.

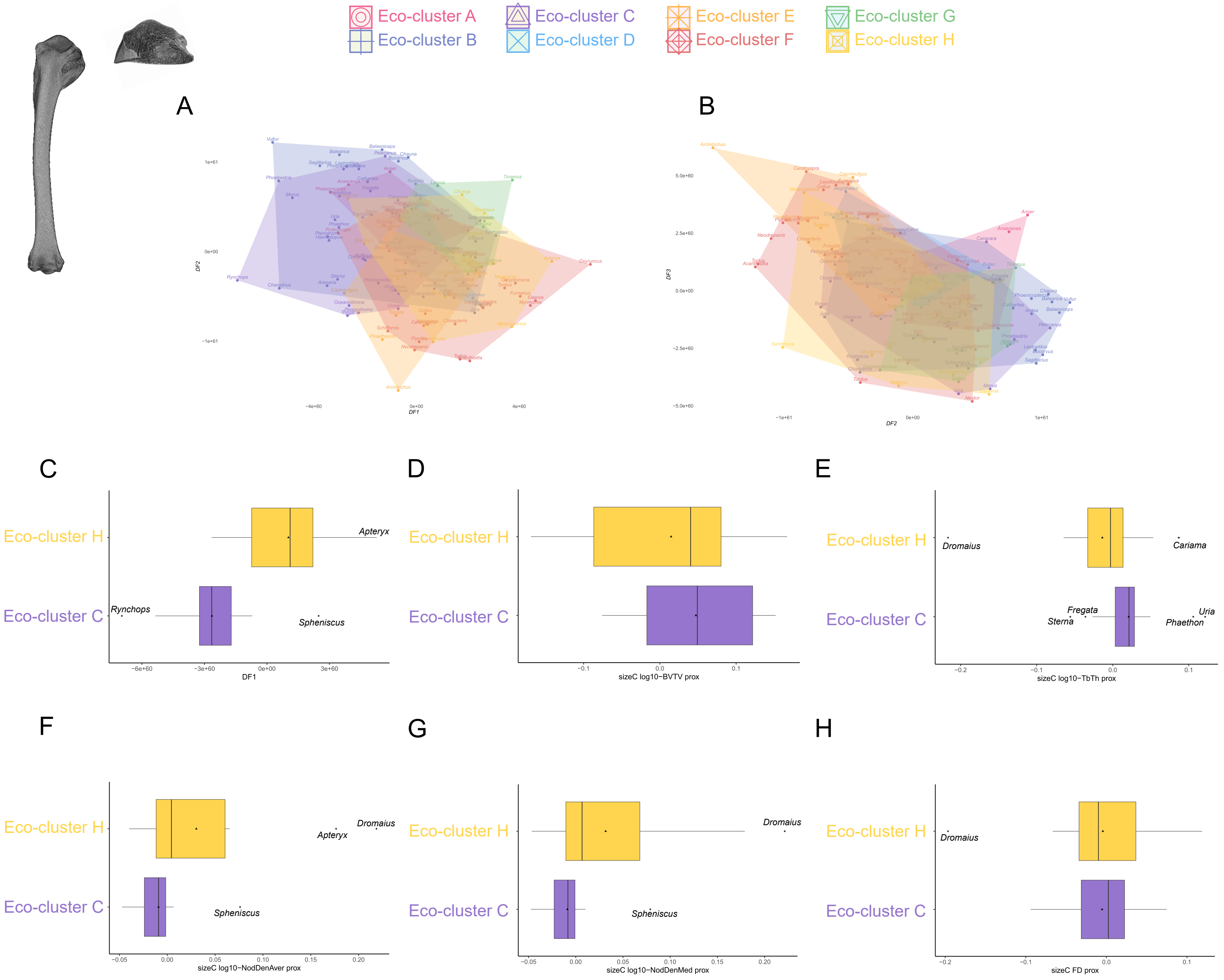

### Results for proximal ulnar structural parameters.

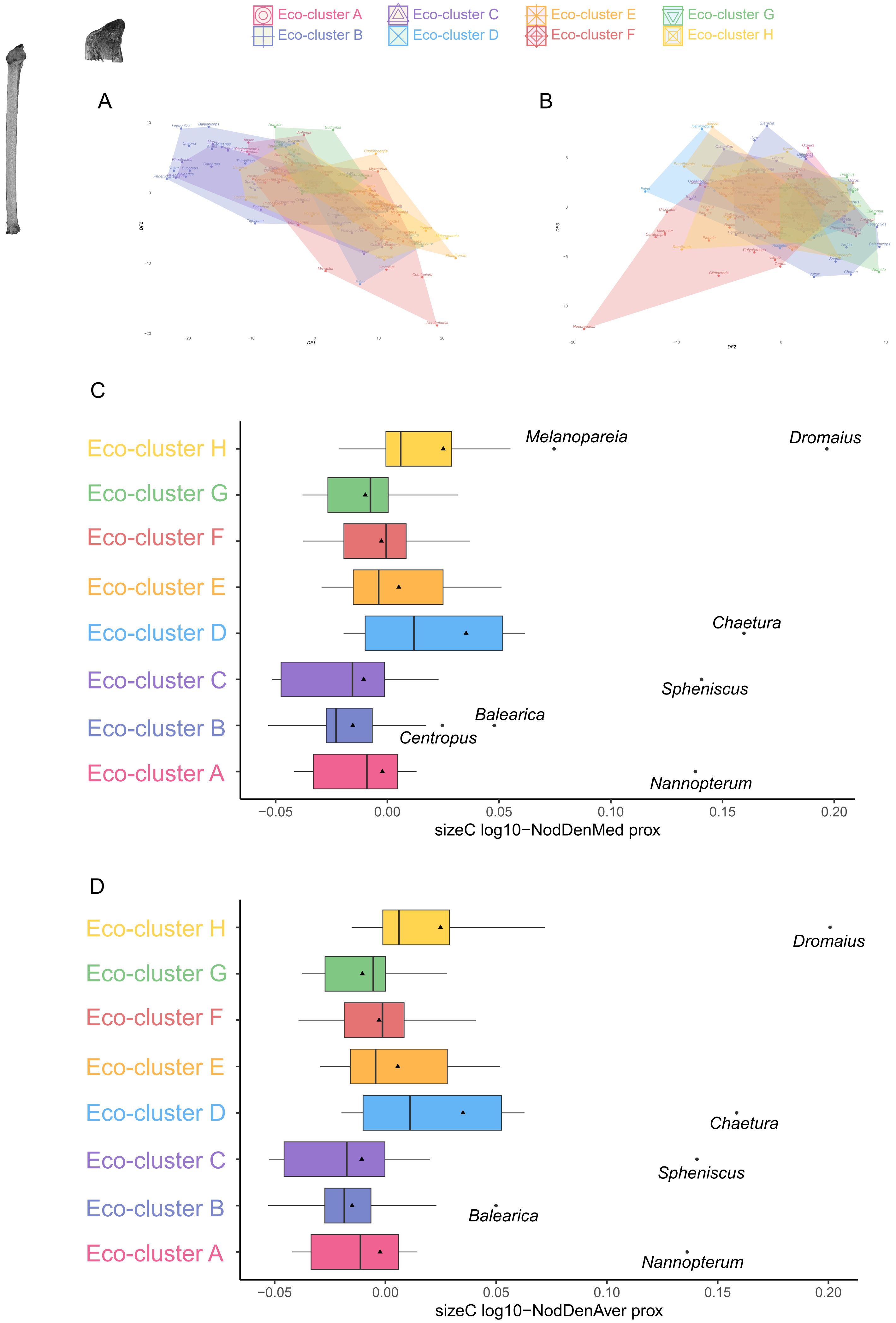

### Results for ulnar diaphyseal structural parameters.

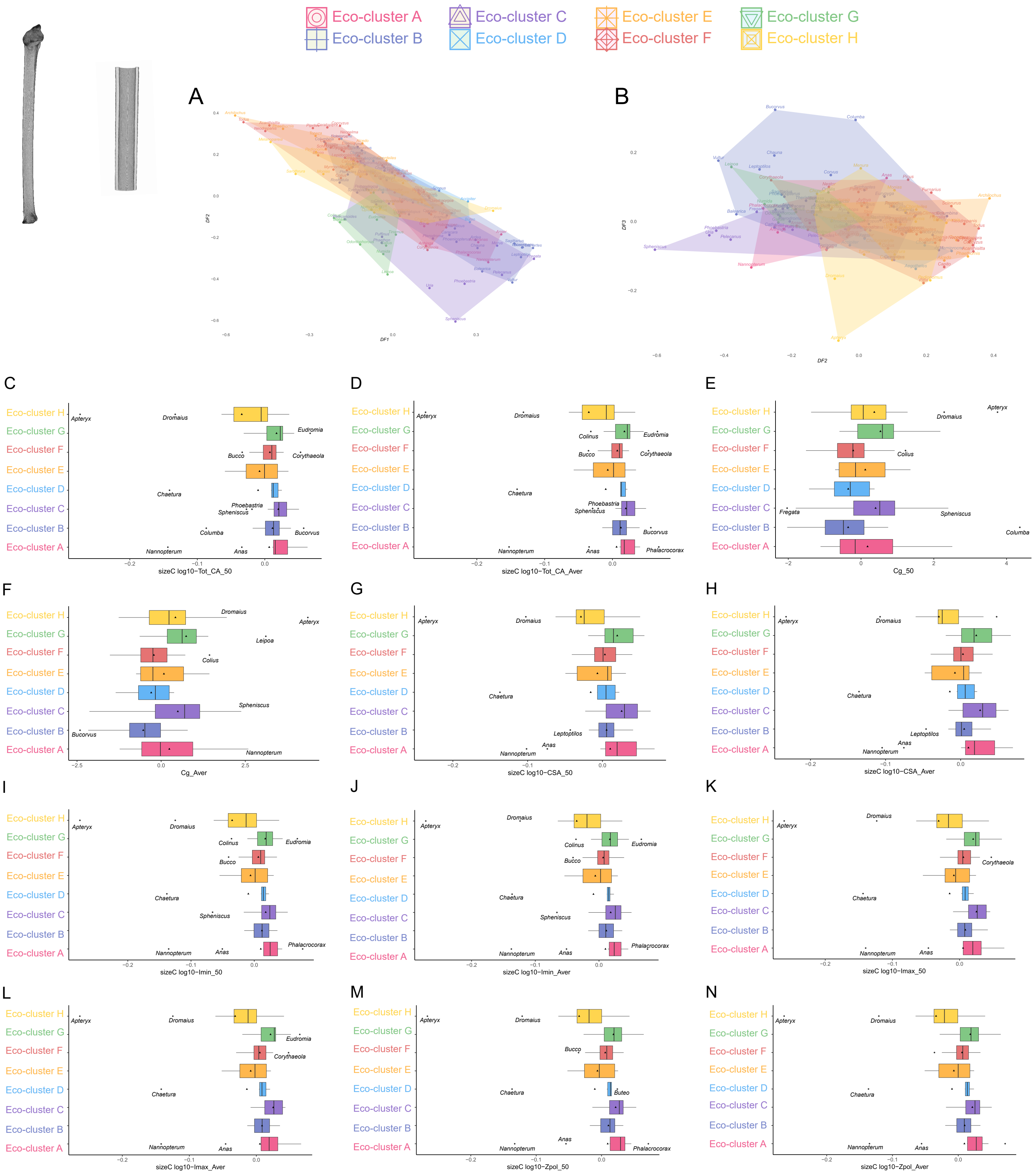

### Traitgrams for humeral diaphyseal parameters of interest (identified as detailed in the Main Text)

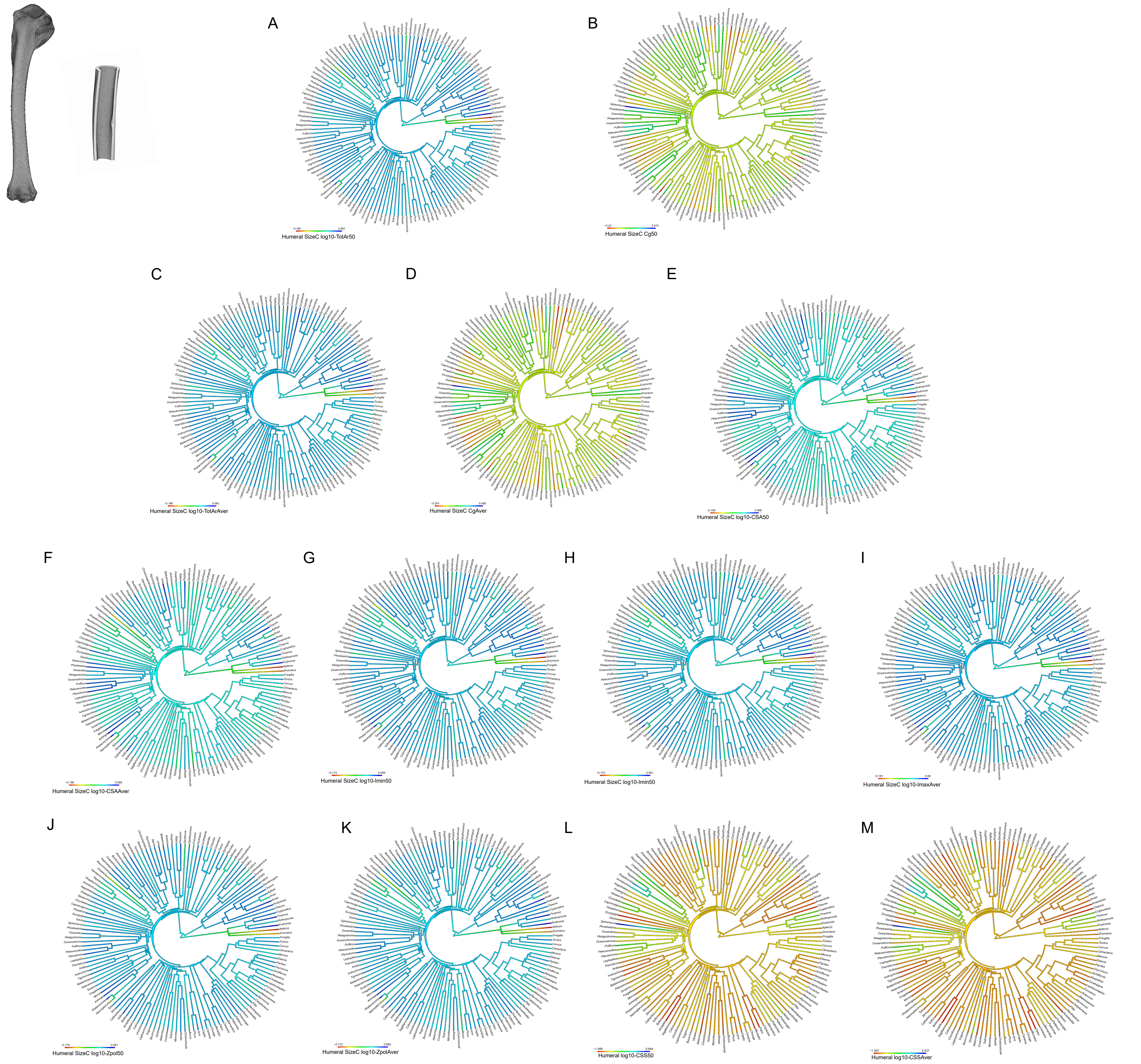

### Traitgrams for proximal humeral parameters of interest (identified as detailed in the Main Text)

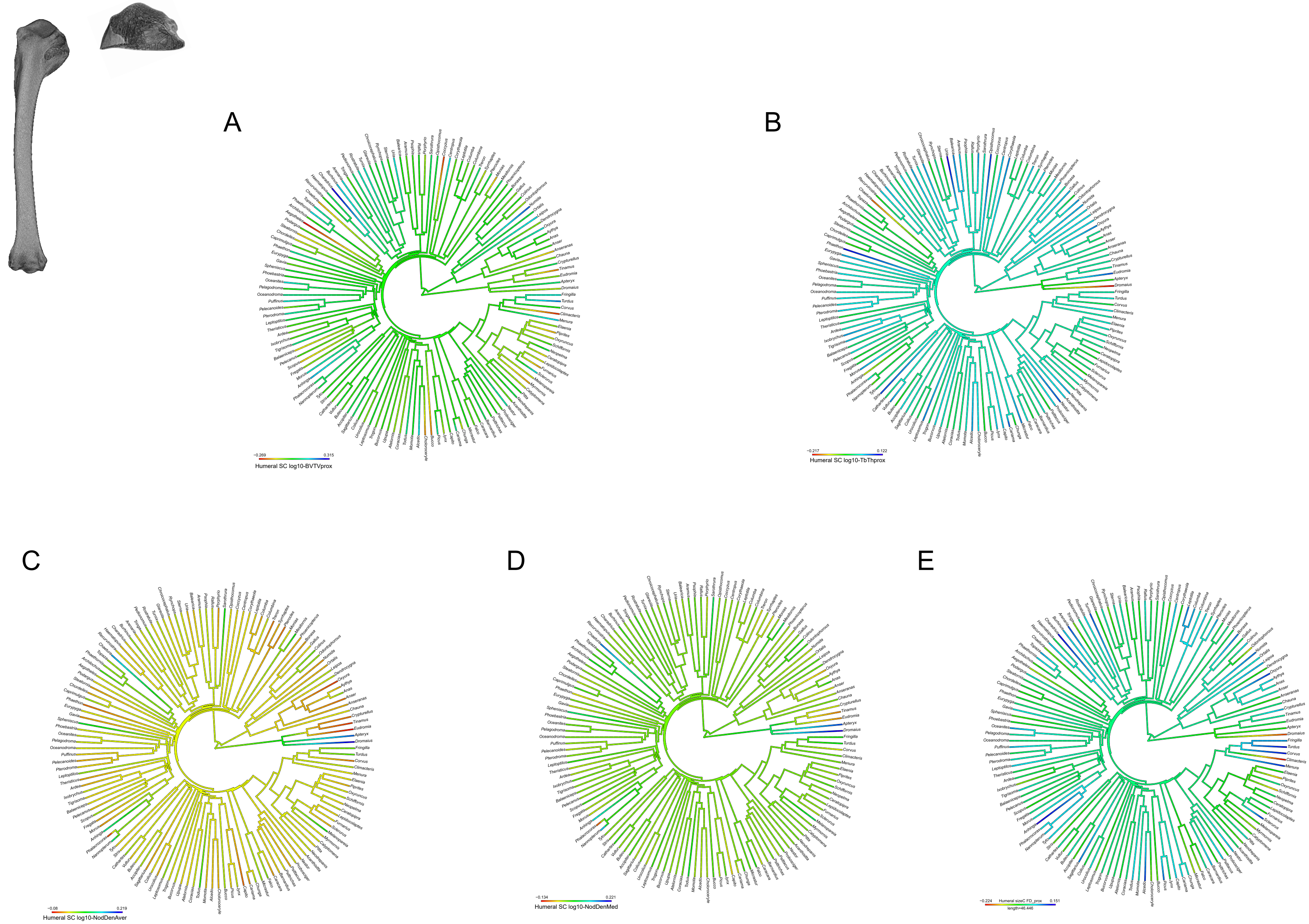
